# CENTRA: Knowledge-Based Gene Contexuality Graphs Reveal Functional Master Regulators by Centrality and Fractality

**DOI:** 10.1101/2025.06.30.662180

**Authors:** Frank Hause, Alice Wedler, Rene Keil, Laura Schian, Markus Glaß, Wiebke Günther, Oleksandr Sorokin, Wolfgang Hoehenwarter, Andrea Sinz, Stefan Hüttelmaier

**Affiliations:** Institute of Molecular Medicine, Section for Molecular Cell Biology, Faculty of Medicine, Martin Luther University Halle-Wittenberg, 06120 Halle, Germany; Center for Structural Mass Spectrometry, Martin Luther University Halle-Wittenberg, 06120 Halle, Germany; Department of Pharmaceutical Chemistry and Bioanalytics, Martin Luther University Halle-Wittenberg, 06120 Halle, Germany

**Author notes:** To whom correspondence should be addressed. Frank Hause; Stefan Hüttelmaier.

## Abstract

Deciphering gene function via context-aware approaches is limited by various means. Especially static gene sets used in enrichment analyses and the lack of single-gene resolution in such analyses restrains the flexible association of genes with specific context. Here, we introduce CENTRA (Centrality-based Exploration of Network Topologies from Regulatory Assemblies), a framework that models gene contextuality through topic-specific gene co-occurrence networks derived from curated gene sets and associated literature. Using Latent Dirichlet Allocation on 12,045 abstracts linked to MSigDB C2 gene sets, we uncovered 27 biological topics and constructed corresponding topic-specific networks that reflect distinct biological states, perturbation conditions, and disease-related regulatory programs. Graph-topological metrics, including centrality, local fractality, and perturbation sensitivity, were computed for each gene to capture structural relevance within these topic-specific contexts. We demonstrate that topological profiles distinguish well-characterized regulators, identify emerging functional candidates, and reveal context-specific roles. Thereby, our framework enables the prioritization of understudied genes by assessing the robustness of their topological signatures across topic-specific networks. To support exploration of these results, we developed a publicly accessible interactive browser application, CENTRA, which enables dynamic navigation of networks and their functional annotations. CENTRA provides an interpretable, scalable framework for investigating context-dependent gene function and hypothesis generation, offering a novel entry point beyond traditional enrichment approaches.

**Graphical Abstract:** 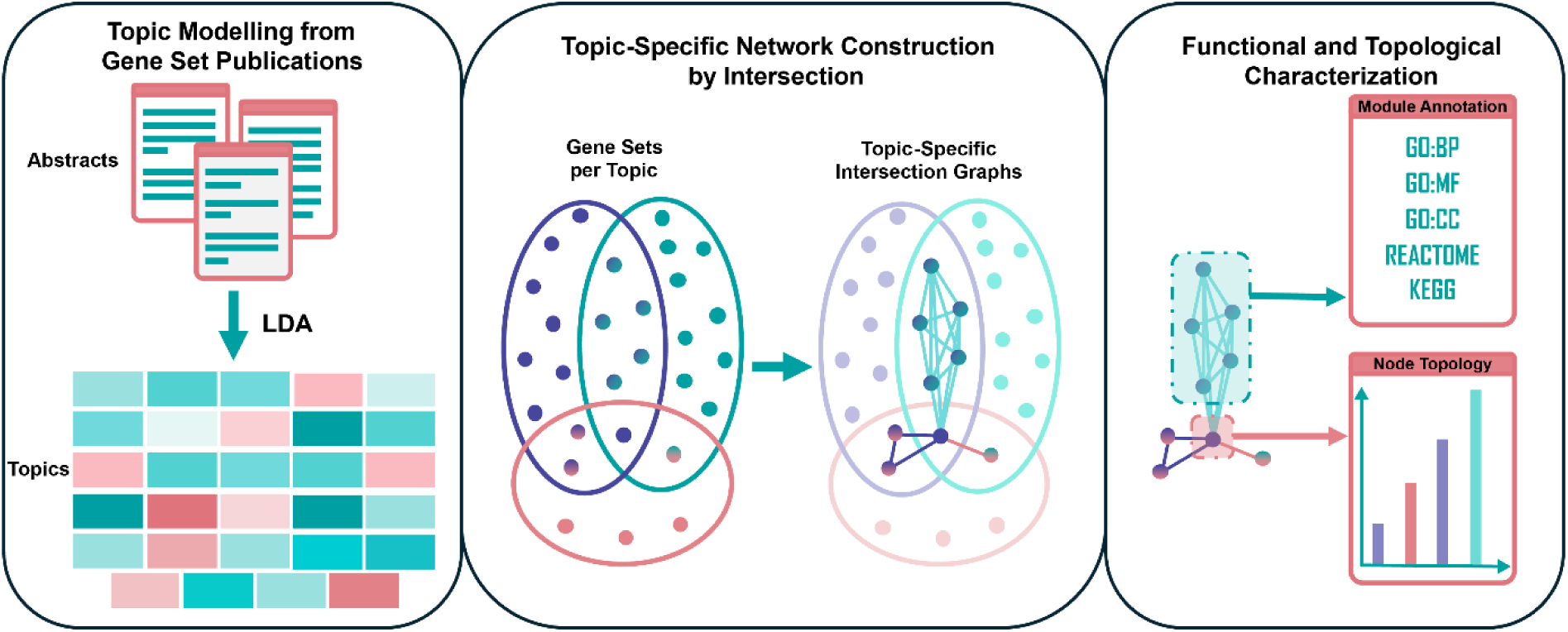

## Introduction

In most biological systems, genes and their products usually do not exhibit only one function exclusively. Gene function is rather highly context-dependent, shaped by cell type, developmental stage, physiological state, and disease condition.^1,2^ A canonical example of such context dependency is TP53, which under physiological conditions serves as a tumor suppressor by orchestrating senescence, DNA repair, and apoptosis.^3,4^ In contrast, when mutated in a tumorigenic context, the same gene can acquire gain-of-function properties that actively promote tumor progression, invasion, and resistance to therapy.^5–7^ This functional duality underscores the necessity of evaluating gene activity within the specific biological and pathological context in which it occurs.

The context-dependency of gene function presents a critical challenge for widely used functional enrichment analyses,^8^ particularly Overrepresentation Analysis^9^ (ORA) and Gene Set Enrichment Analysis^10^ (GSEA). While both methods have become standard tools for the downstream interpretation of differential gene or protein abundance data, they share two fundamental limitations: (i) They require a predefined set of genes to infer functional or pathway involvement, and (ii) they do not explicitly account for the biological context in which this set, or any individual gene, is relevant. ORA assigns functional significance by testing whether predefined gene sets, reportedly associated with specific pathways or conditions, are statistically overrepresented among a list of genes of interest, under the assumption that these genes act in concert within a shared biological process. GSEA, by contrast, assesses whether genes from predefined sets are disproportionately enriched at the top or bottom of a ranked list, typically ordered by fold-change, thus assigning functional relevance based on gene abundance shifts in a yet context-dependent experimental setting.

Both ORA and GSEA rely on reference gene sets that have been experimentally linked to specific perturbations, pathways, or disease states. These gene sets are curated in public repositories such as the Molecular Signatures Database (MSigDB)^11,12^, particularly its C2 collection, which contains curated gene sets from Canonical Pathways (CP) as well as Chemical and Genetic Perturbations (CGP). While these resources are invaluable for capturing broad biological themes, they are not designed to offer contextual relevance of individual genes. While researchers have the possibility to consult gene set annotations from different repositories when performing ORA or GSEA, the resulting enrichment terms are sometimes broad or generic, making biological interpretation difficult. This challenge is compounded by genes appearing in multiple gene sets linked to different pathways, blurring their specific functional context. More importantly, both approaches are fundamentally designed to assign function to sets of genes that show coordinated alteration under specific conditions, rather than to individual genes. As such, they cannot directly inform on the specific role or relevance of a single gene within a particular biological context. Gaining such gene-level insight typically requires integrating prior knowledge from literature; a process that is time-consuming, fragmented, and difficult to formalize within standard enrichment frameworks.

Topic modeling provides a computational strategy to systematically analyze large bodies of text and extract thematic patterns without relying on predefined categories or prior assumptions.^13,14^ It can be applied to entire publications or specific sections— such as abstracts or methods—to uncover latent semantic structures that reflect underlying biological concepts. A widely used approach in this domain is Latent Dirichlet Allocation (LDA),^15,16^ a generative probabilistic model that represents each document as a mixture of topics, and each topic as a probability distribution over words. By inferring these topic distributions, LDA identifies hidden thematic groupings that link related concepts across large and heterogeneous corpora.

In this study, we leverage LDA to annotate curated MSigDB gene sets with latent biomedical themes, using these topic assignments to construct topic-specific gene co-occurrence networks. Within each topic-specific network, genes are embedded according to their co-occurrence patterns, enabling the computation of structural properties such as centrality, local fractality, and robustness measures. This topological embedding reflects a gene’s relative contextual importance, revealing both well-established regulators and understudied genes whose structural roles suggest latent functional relevance. The resulting framework not only addresses a critical gap in the functional annotation of genes under specific perturbations or disease states but also offers an accessible platform, CENTRA (<u>C</u>entrality-based <u>E</u>xploration of <u>N</u>etwork <u>T</u>opologies from <u>R</u>egulatory <u>A</u>ssemblies), for hypothesis generation and exploratory analysis. We anticipate that this approach will support a deeper understanding of context-specific gene function and guide future studies in systems biology, network medicine, and translational research.

## Methods

### Construction of Topic-Specific Gene Co-Occurrence Networks

Gene sets and the abstracts of the reporting publications (for PubMed IDs [PMIDs] of included abstracts, see Supplementary File S1) were retrieved from the Curated Gene Set Collection C2 (CP and CGP) of the Molecular Signatures Database^11,12^ (MSigDB v2023.1.Hs) using the msigdbr package (Dolgalev, 2025). To compensate for missing PMIDs, gene set descriptions from Reactome^17^, the Kyoto Encyclopedia of Genes and Genomes^18^ (KEGG), and WikiPathways^19^ were retrieved using their respective application programming interfaces (APIs) or web services.

Abstracts corresponding to the annotated PMIDs were retrieved from PubMed using the NCBI E-utilities (rentrez package) (Winter, 2017). In case no abstract was available, the original gene set description was used. If multiple abstract annotations were present for a single gene set, all corresponding abstracts were concatenated into a single document prior to downstream processing. Abstracts and descriptions (hereafter referred to as documents) underwent a preprocessing pipeline using tm^20^ including (i) character normalization, (ii) removal of Greek and other non-Latin alphabet residues, (iii) lowercasing, (iv) punctuation and number filtering, (v) whitespace trimming, and (vi) stemming. Custom stopword removal was applied using a manually curated stopword list (see Supplementary File S2). Rare token (frequency < 5) were iteratively removed.

From all preprocessed documents, a document-term matrix was constructed. LDA^15,16^ was used to infer latent topics across the corpus. The final number of topics (k) was computed as function of assignment confidence of the preprocessed documents based on the posteriors of several LDA models (k = [10;50]). Several iterations were conducted to optimize interpretability and consistency. Topics were assigned human-readable labels based on the top 100 β-weighted terms from each topic. These were manually interpreted and ranked for internal consistency. The final model used k = 27 topics. A heuristic confidence threshold of 0.2 was used to retain only confident document-to-topic assignments while maximizing the number of classifiable documents.

For each topic, pairwise co-occurrence of gene symbols was inferred by finding intersections across every possible two gene sets; gene pairs found in shared sets formed unweighted and undirected edges in a network with nodes corresponding to genes and edges reflecting co-occurrence, using igraph^21^. Multiple overlaying edges were collapsed to unique connections to eventually form a knowledge-based topic-specific gene co-occurrence graph.

### Network-Level Analysis

Each topic-specific network was characterized by the following global topological metrics: node and edge counts, average degree, clustering coefficient, graph density, diameter, average path length, Louvain modularity, average betweenness and closeness centrality.

To compare the structural similarity of networks not sharing the same node structure, spectral similarity was assessed using Laplacian eigenvalues^22^ (top 20 components) of each network. Pairwise Euclidean distances between truncated spectra were computed to generate the corresponding distance matrix.

To assess node-level betweenness robustness (see below), local edge perturbation was simulated across 1,000 iterations (1‰ random edge rewiring per iteration) per network. For each iteration, betweenness centrality was recalculated. Per-gene variances in this metric were recorded, capturing the topological stability under minimal structural change possibly hinting at important roles of genes in a topic-specific network due to limited availability of those genes in the assigned gene sets.

### Module-Level Analysis

Louvain clustering was used to define modules within each topic network. For each module, the following topological metrics were computed: node and edge counts, average degree, graph density, and average path length.

Module-level functional enrichment was performed using g:Profiler^23^, restricted to annotations from Gene Ontology^24,25^ (GO), Reactome^17^, and KEGG^18^. Enrichment was computed using adjusted p-values (false discovery rate [FDR] < 0.05).

To visualize functional relationships among topics based on their most significant biological processes, a semantic similarity map was constructed using the top enriched GO:BP terms per topic. For each topic, the 100 most significant biological process terms (based on adjusted p-values) were selected. These terms were mapped into a two-dimensional space using classical multidimensional scaling (MDS) of pairwise semantic similarity scores derived from GOSemSim^26^ analysis.

The resulting MDS coordinates were used to calculate a centroid position for each topic, defined as the average semantic coordinates of its top 100 enriched GO terms.

These centroids provide a summary representation of the functional position of each topic within the semantic ‘space’ of the annotated network.

### Node-Level Analysis

For each node, the following topological metrics were computed: centrality measures (betweenness and eigenvector centrality). Additionally, local fractal dimension (LFD) was computed node-wise following Xiao, Chen, and Bogdan^27^. Robustness of betweenness centrality was assessed in an edge perturbation simulation by local edge perturbation (see above).

### CENTRA Implementation

To provide an accessible and interactive interface for exploring the resulting topic-specific networks, modules, and gene-level metrics, CENTRA was developed, a Shiny-based (Chang, Cheng, Allaire, et al., 2025) web browser application. CENTRA enables users to intuitively navigate and interpret the structural and functional properties of topic-specific co-occurrence networks derived from the curated gene set analysis. Designed to be low-barrier and hypothesis-generating, CENTRA allows researchers without advanced computational expertise to explore the topological metrics of different genes across diverse biological themes.

The application integrates the precomputed LDA topic assignments, gene co-occurrence networks, module clustering, functional enrichment results, and topological metrics at the network, module, and gene level. Users can interactively select topics, visualize network structure, explore functional module enrichment, and comparatively inspect gene-specific metric profiles across the analysis topic-specific networks. All visualizations are powered by networkD3 (Allaire, Ellis, Gandrud, et al., 2017) to ensure smooth interaction and responsive layout behavior.

CENTRA has been independently tested by several scientists with diverse backgrounds to evaluate its usability and interpretability. CENTRA is available for review at: [link] (*access upon request*).

## Results

### General Analysis Strategy

To establish a method that, unlike ORA and GSEA, supports functional annotation with single-gene granularity and contextual relevance, we leveraged the gene set-reporting literature associated with the MSigDB C2 cluster of curated gene sets. Specifically, we applied LDA to cluster the abstracts linked to these gene sets into distinct biomedical topics, each representing a characteristic biological process, condition, or functional theme. This clustering enabled us to reorganize gene sets and their constituent genes based on co-occurrence within shared topical contexts. By assigning each gene occurrence to one or more topics through its inclusion in gene sets with overlapping textual associations, we constructed knowledge-based, topic-specific gene co-occurrence networks, in which nodes represent genes and edges denote co-occurrence within sets assigned to the same topic (see Figure 1a-c).

**Figure 1:**
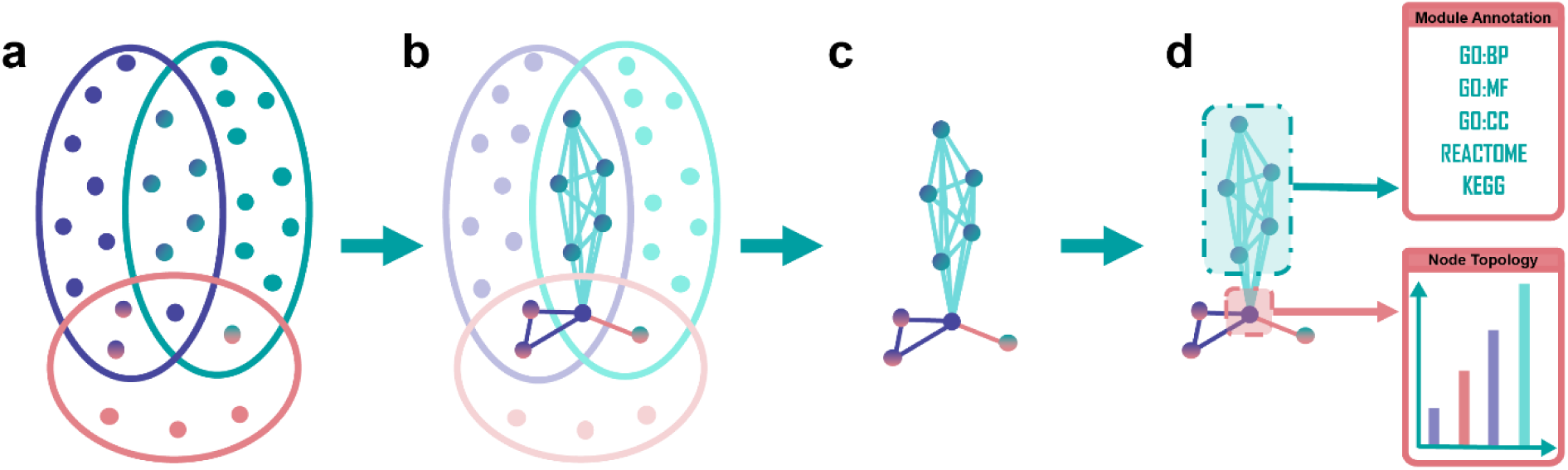
Construction of topic-specific gene co-occurrence networks. (**a**) Gene sets (colored ellipses) assigned to a topic via LDA exhibit partial overlaps in their constituent genes (colored dots). (**b**) Pairwise intersections were computed across all gene sets within a topic. Genes shared between sets were connected as undirected edges in an unweighted graph. (**c**) Recurrent edges from multiple overlapping gene sets were collapsed into single, unique interactions to generate a knowledge-based topic-specific gene co-occurrence network. (**d**) The resulting graph was further analyzed by identifying modules, performing functional annotation along with quantification of topological features at the node level.

The topological position of a gene within a topic-specific network serves as a proxy for its functional relevance within a given biological context.^28^ To capture this, we computed a set of interpretable graph metrics that reflect different aspects of a gene’s potential role within a particular context: A high betweenness centrality highlights genes that act as bridges between otherwise unconnected regions of the network, pointing to potential integrators, regulators, or “bottlenecks” across functional modules. Eigenvector centrality extends the idea of gene’s importance characterized by a short overall distance to other genes by not only counting connections but weighing them by the importance of neighboring genes, capturing influence within the broader network hierarchy. Finally, high values of LFD reflect an increasing structural complexity in the vicinity of a gene, highlighting genes embedded in tightly organized or hierarchically structured neighborhoods.^29^

Together, these metrics offer a multifaceted characterization of a gene’s contextual role, whether it is broadly connected, strategically positioned, or deeply embedded in a functionally cohesive cluster (see Figure 1d). These features can signal different forms of biological importance, from hub-like activity to local specialization.

To account for genes that appear in only a few gene sets but may nonetheless play important roles within a given topic-specific network, the structural stability of their topological metrics was assessed through topic-specific random edge rewiring. In this procedure, networks were perturbed by randomly reassigning a small fraction of edges, followed by recalculation of betweenness centrality. This approach enabled the estimation of each gene’s topological robustness across perturbations, serving as a proxy for stability, and thus potential reliability, of its inferred contextual relevance.

Importantly, the variance of key metrics such as betweenness centrality across perturbed networks may offer predictive cues for interpretation. For example, low absolute centrality values combined with high variance suggest that a gene’s network position is highly sensitive to small changes in connectivity—such as a few edges being added or removed. This indicates that the gene can occasionally assume a central or regulatory role depending on minor alterations in the network structure, potentially reflecting a functionally relevant but understudied gene in the given topic. Conversely, high metric values with high variance may point to an unstable topological position, where apparent centrality arises from configurations that are easily disrupted. Such cases call for cautious interpretation, as network prominence may not be reliably sustained. In contrast, genes with consistently high metric values and low variance are more likely to represent robust and biologically meaningful regulators within their respective contexts.

To make these data and insights widely accessible, we developed CENTRA, a publicly available, interactive web application that allows users to explore these networks and gene metrics in an intuitive manner. CENTRA supports topic selection, enrichment exploration, and node-level metric visualization offering a novel entry point for researchers interested in the context-dependent roles of genes in health and disease.

### Probabilistic Clustering of Gene Set Documents Reveals Topic-Specific Network Structures

To uncover latent thematic structure within the corpus of gene set-associated literature, probabilistic LDA topic modeling was applied to 12,045 abstracts associated with 6,366 gene sets from the C2 collection of MSigDB. This non-deterministic approach yielded 27 distinct topics reflecting diverse biological processes, perturbation conditions, and disease-related regulatory programs. These topics are characterized by distinct token distributions that capture their semantic composition (see Figure 2a). As a surrogate for this semantic composition, the β weight reflects the relative importance of a token within a given topic. For example, in the topic “DNA Damage Response and Repair Mechanisms” the tokens “repair,” “damage,” and “dna” yield the highest β weights; “Lipid Metabolism and Membrane Phospholipid Biosynthesis” is enriched for “lipid,” “phospholipid,” and “metabolism”; while “Neurodegenerative Diseases and Mitochondrial Dysfunction” prominently feature “mitochondria,” “degeneration,” and “neuron”.

**Figure 2:**
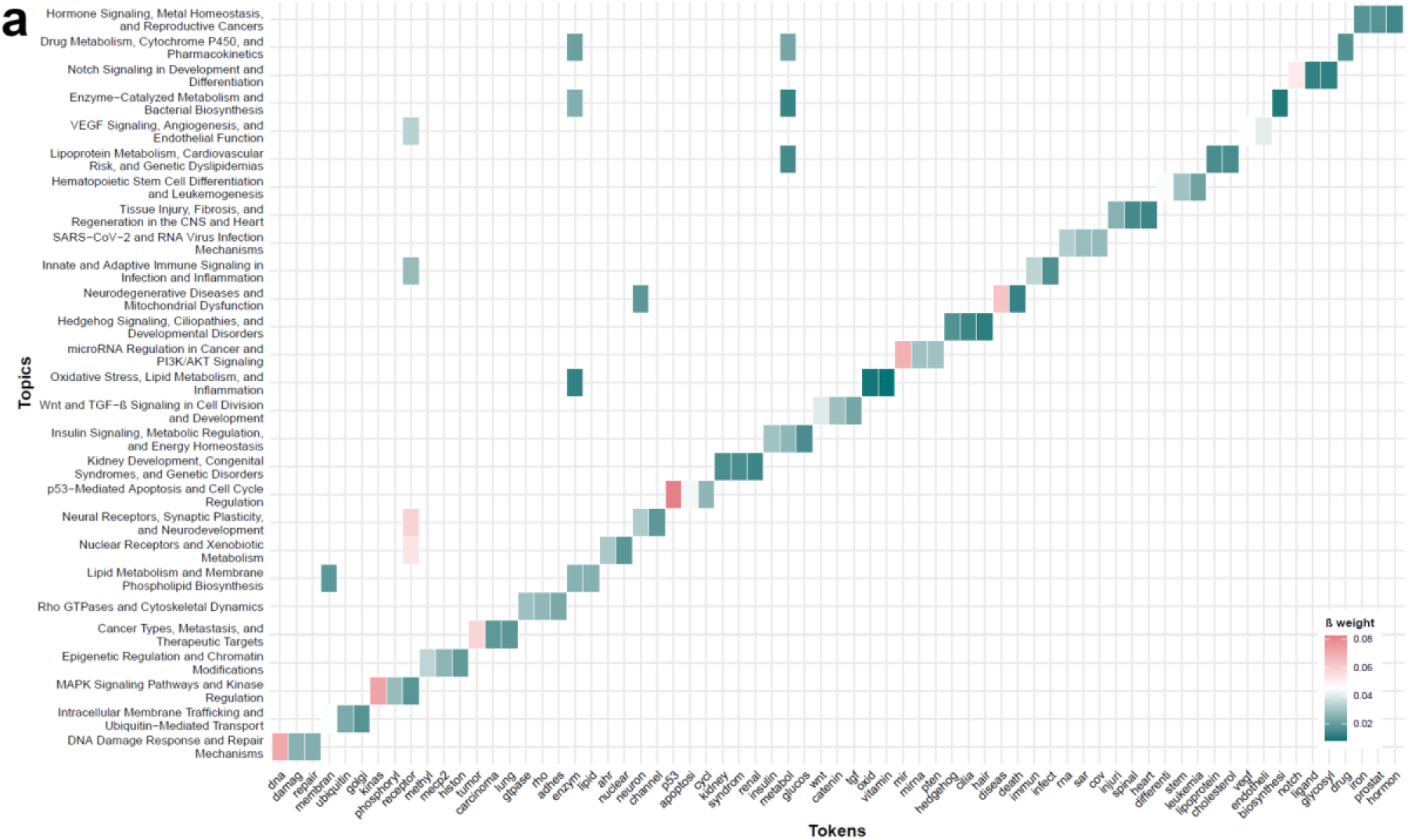
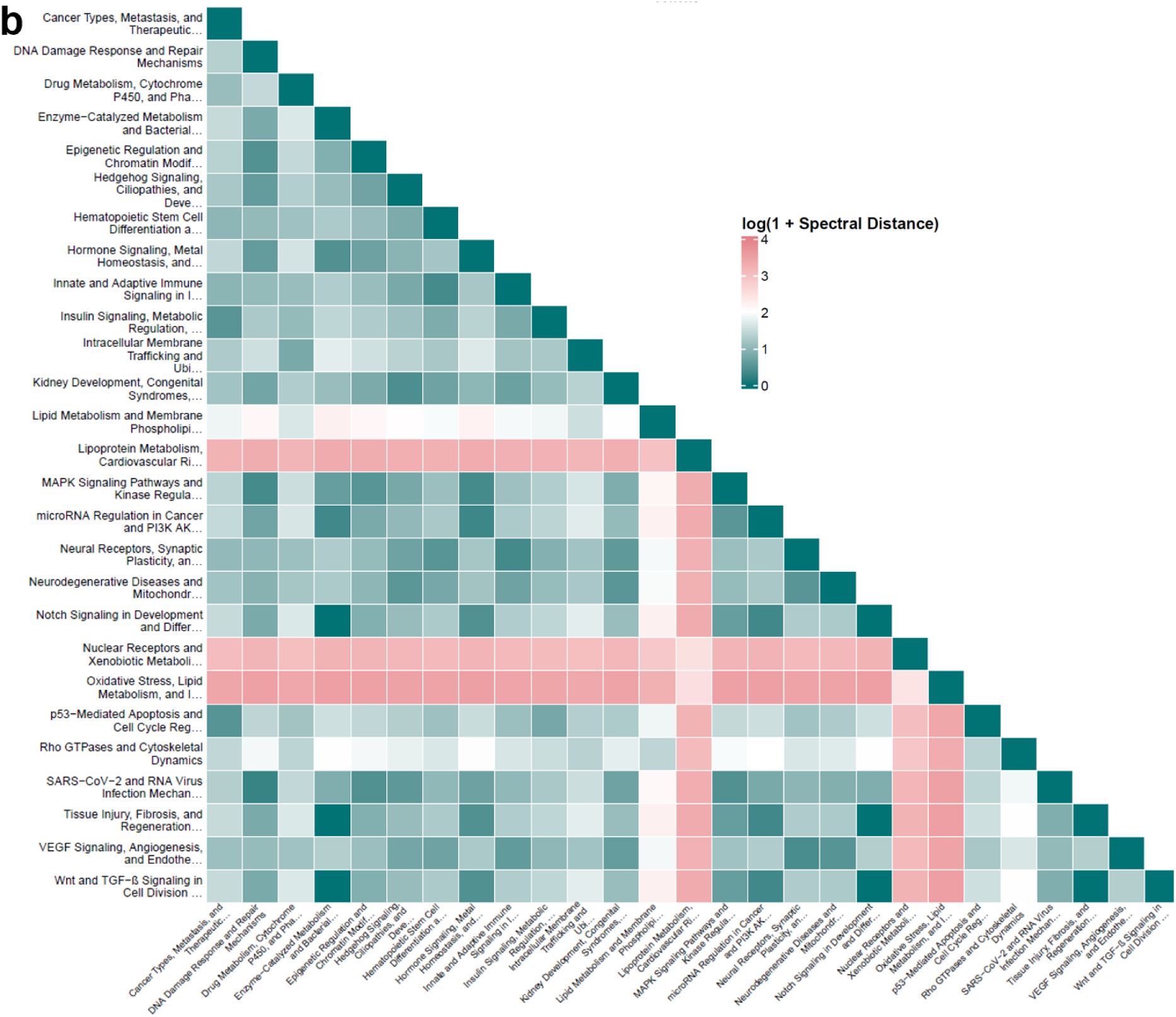
Topic modeling and network-level stratification of MSigDB C2 gene sets. (**a**) Token–topic distribution for the 27 topics derived from LDA modeling of 12,045 abstracts and gene set descriptions from the MSigDB C2 collection. Shown are the three highest β-weighted tokens per topic across all topics, with cell color representing the respective β-weight within the LDA model. (**b**) Spectral distance matrix indicating the structural similarity of the topic-specific gene co-occurrence networks. Values reflect log-transformed spectral distances, highlighting both closely related and structurally divergent topic-specific networks.

Documents were classified to a topic based on the posterior probability distribution of the LDA model. A threshold of 0.2 for the maximum posterior probability was chosen, balancing interpretability of topic content with the number of documents confidently assigned. Using this criterion, 2,052 documents were classified to a respective topic. Topic assignment was unevenly distributed, with the largest number of documents assigned to “Cancer Types, Metastasis, and Therapeutic Targets” (n = 276), and the fewest to “Oxidative Stress, Lipid Metabolism, and Inflammation” (n = 12). It can be assumed that these stark differences mirror broader trends in biomedical research, where cancer continues to represent a dominant focus of scientific activity. A principal component analysis (PCA) of the document–topic assignment confidence matrix further supports the discriminability of topic structures, with documents forming partially separable clusters in 2D space according to their topic assignments (see Supplementary Figure S1). This degree of separation is consistent with expectations for LDA, which models topics as distinct yet potentially overlapping distributions. Indeed, while topics are clearly distinguishable, they are not fully orthogonal, a property that reflects the shared thematic content of many biological abstracts. This is further illustrated at the token level, where token such as “receptor” rank among the top three β weights in five different topics, underscoring the biological interconnectedness of the thematic structures inferred.

Within each topic, gene co-occurrence networks were constructed by identifying shared genes across all pairwise combinations of gene sets assigned to that topic. These topic-specific networks varied considerably in size and topology, with key properties including node count (range: 48 to 12,922), edge count (range: 248 to 2,531,635), edge density (range: 0.017 to 0.537), and average degree (range: 10.3 to 391.8) (see Supplementary Table S1). Structural divergence between networks was quantified using pairwise spectral distances (see Figure 2b). Topics with overlapping biological scope, such as “Lipid Metabolism and Membrane Phospholipid Biosynthesis” and “Lipoprotein Metabolism, Cardiovascular Risk, and Genetic Dyslipidemias”, exhibited low spectral distances, indicating structurally similar architectures. In contrast, networks derived from topically and mechanistically divergent contexts, e.g., “Oxidative Stress, Lipid Metabolism, and Inflammation” versus “Wnt and TGF-β Signaling in Cell Division and Development”, showed pronounced spectral separation. These findings demonstrate that meaningful gene co-occurrence networks can be derived by clustering the gene set documents using LDA, resulting in unique networks that inherently reflect the varying degrees of gene co-occurence, regulatory modularity, and pathway specificity and thereby preserving the structural relationships of the embedded gene sets.

### Module Detection Reveals Functionally Unique Assemblies within Topic-Specific Networks

To assess whether topic-specific gene co-occurrence networks exhibit internally coherent substructures, Louvain community detection was applied to each network. This algorithm partitions the network into modules—densely interconnected groups of genes—based solely on topological properties. As the method is unsupervised and independent of prior biological knowledge, the resulting modules reflect purely emergent structural patterns derived from gene co-occurrence within each topic.

Despite substantial variation in size and connectivity across topic networks, modular structure was consistently detected. An illustrative example is provided in Figure 3a, which displays the network “Oxidative Stress, Lipid Metabolism, and Inflammation”. This network comprises 75 genes that are part of intersections among 12 gene sets assigned to this topic, making it the smallest of the 27 topic-specific graphs in terms of contributing documents. Nonetheless, three distinct modules were identified, each representing a subset of genes that exhibit a particular connection pattern within the topic-specific network.

**Figure 3:**
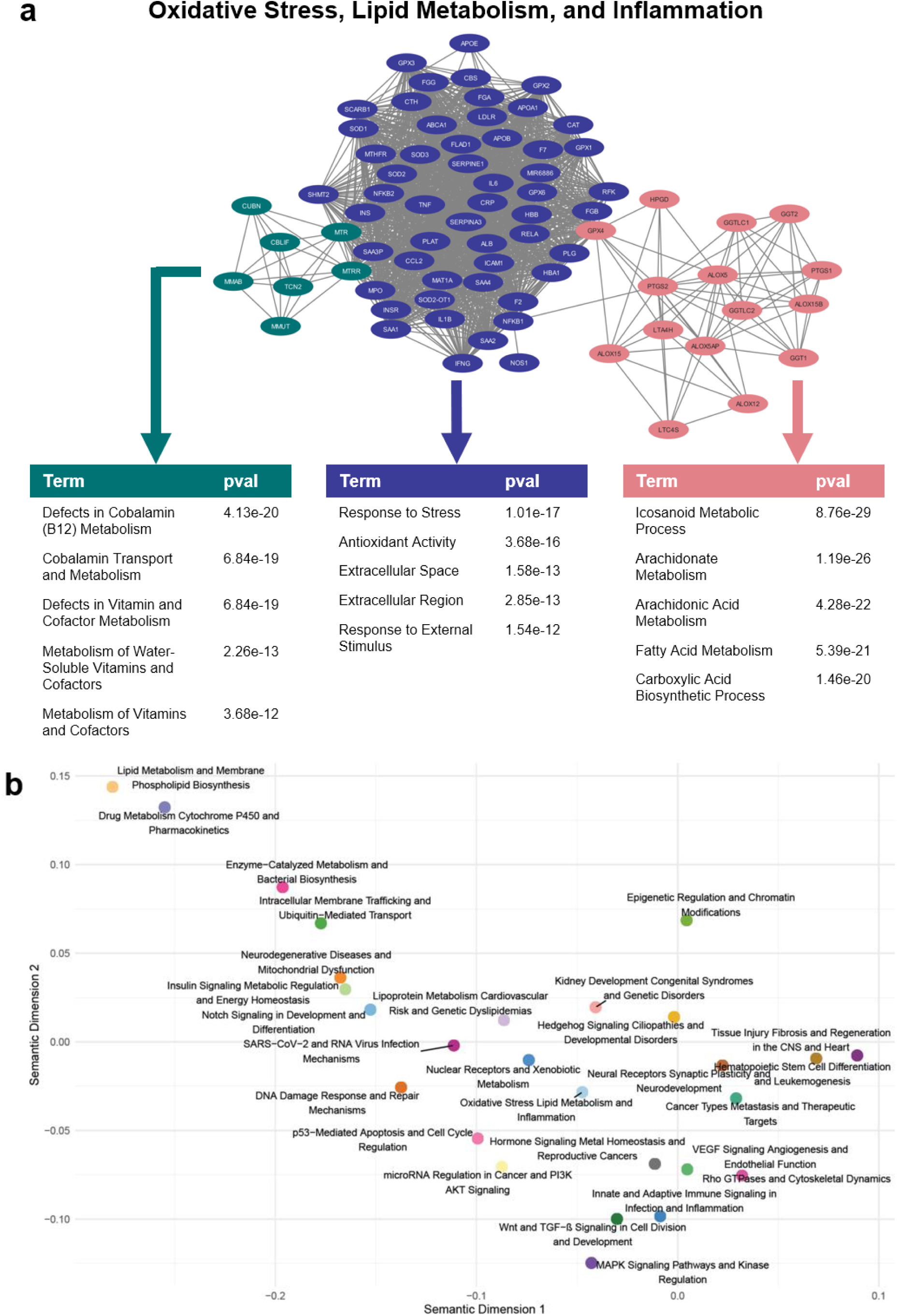
Functional annotation of Louvain-derived modules across topic-specific networks. (**a**) Example of a topic-specific network for “Oxidative Stress, Lipid Metabolism, and Inflammation”, the smallest network in the dataset (75 genes). Louvain clustering identifies three distinct modules, highlighted in different colors: the green module is functionally enriched for vitamin metabolism, the red module for fatty acid metabolism with a focus on inflammation, and the purple module for cellular stress response. (**b**) Semantic similarity map of the 27 topics based on the 100 most significant GO:BP terms of their modules. Each point represents one topic-specific network, positioned using multidimensional scaling of semantic distances among enriched terms.

Following module detection, functional characterization was performed using ORA on the genes within each module, utilizing gene sets from GO^24,25^, Reactome^17^, and KEGG^18^. The semantic similarity map in Figure 3b reveals clear separation of topics based on the functional profiles of their annotated modules. Each point represents the centroid of a topic, summarizing its most enriched GO biological process terms. The spatial distribution highlights distinct clusters of biologically related topics: for example, “Lipid Metabolism and Phospholipid Biosynthesis” appears near “Drug Metabolism Cytochrome P450 and Pharmacokinetics”, reflecting shared metabolic processes, while signaling and development-related topics form a separate cluster encompassing “MAPK Signaling Pathways and Kinase Regulation”, “Wnt and TGF-β Signaling in Cell Division and Development”, and “Innate and Adaptive Immune Signaling in Infection and Inflammation”. This separation reflects the functional coherence and diversity captured by the modular annotations and underscores the biological specificity embedded in each topic’s network structure.

### Centrality and Fractality Metrics Uncover Context-Dependent Importance of Genes across Topic-Specific Networks

The structural position of a gene within a topic-specific co-occurrence network provides a proxy for its contextual importance. It is assumed that this topological embedding reflects how prominently a gene contributes to the functional architecture within a given biological theme. To quantify this embedding, a suite of node-level topological metrics was calculated across all topic networks, including betweenness centrality (bottleneck genes), eigenvector centrality (hubs of important genes), and LFD (hierarchical regulators). In addition, the robustness of selected metrics was estimated by their variances under perturbations by randomized edge rewiring, capturing the structural stability of a gene’s influence.

To illustrate the interpretive value of node-level topological metrics, we now discuss three representative genes that each exhibit a pronounced signal in one of the assessed metrics: GPX4 displays exceptionally high betweenness centrality in the network “Oxidative Stress, Lipid Metabolism, and Inflammation” (see Figure 4a and 4b), highlighting its role as a topological bottleneck that mediates communication between otherwise weakly connected regions. This elevated betweenness suggests that GPX4 frequently lies on the shortest paths between other genes, enabling it to coordinate processes across distinct network modules. Consistently, GPX4 connects modules related to oxidative stress response and fatty acid metabolism (see Figure 3a), reflecting its functional role as a glutathione peroxidase that reduces lipid hydroperoxides and protects cells from ferroptotic death.^30^ Likewise remarkable and consistent is the exceedingly high eigenvector centrality of GPX4 in the network “Neural Receptor, Synaptic Plasticity, and Neurodevelopment” (see Figure 4a). This suggests a high relevance of the gene in neural development, function and response to injury. In agreement, GPX4 upregulation is observed in astrocytes upon brain injury and appears to protect cells from apoptosis.^31^

**Figure 4:**
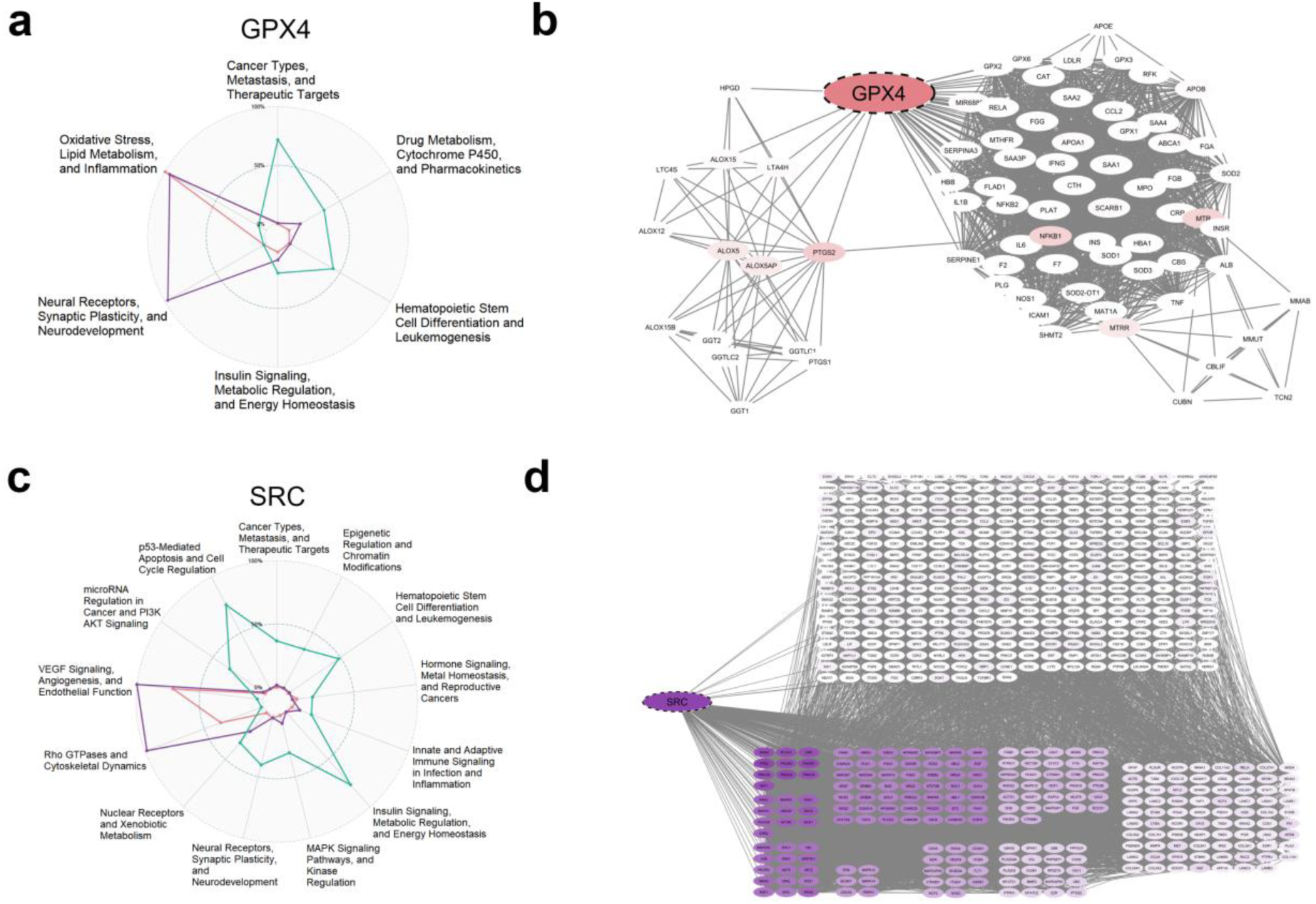
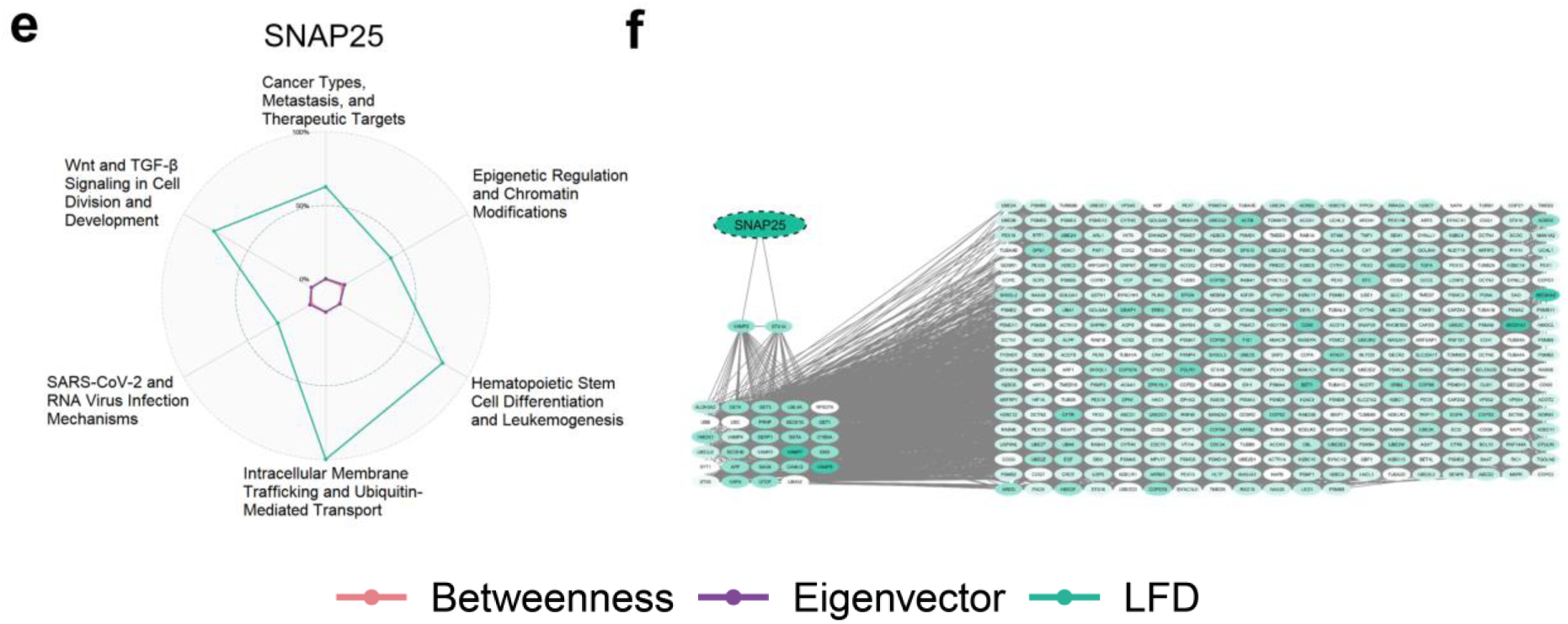
Topological metrics across topic-specific networks reveal context-dependent gene relevance. For each of the three selected metrics, betweenness, eigenvector centrality, and LFD, one representative gene with exceptionally high values in a specific network is shown. The left panels display radar plots of the metric values for the gene across all topic-specific networks, highlighting its contextual specificity; these plots can also be accessed interactively via the CENTRA web application. The right panels depict the local network neighborhood of the gene within the respective topic, colored by topology metric where a more intense color accounts for a higher metric score. (**a–b**) GPX4 shows high betweenness in the “Oxidative Stress, Lipid Metabolism, and Inflammation” network, acting as a bottleneck bridging two main node communities. (**c–d**) SRC is a hub of eigenvector centrality in the network “VEGF Signaling, Angiogenesis, and Endothelial Function”, indicating central influence through connections to other highly central nodes. (**e–f**) SNAP25 has the highest LFD in “Intracellular Membrane Trafficking and Ubiquitin-Mediated Transport”, embedded in a highly branched neighborhood with exponential growth in local connectivity as distance from SNAP25 increases linearly.

SRC displays prominently high eigenvector centrality in the network “VEGF Signaling, Angiogenesis, and Endothelial Function” (see Figure 4c). This metric reflects not just the number of connections, but the influence of those connections: SRC is linked predominantly to other genes with a high eigenvector centrality rather than less central nodes (see Figure 4d). This positions SRC within the network’s core of regulatory influence, suggesting a central role in propagating biological signals through densely connected hubs. This is consistent with SRC’s well-characterized function as a non-receptor tyrosine kinase that integrates growth factor signaling and promotes angiogenic responses, endothelial cell migration, and vascular remodeling which are hallmark processes of the biological theme captured by this network.^32,33^

SNAP25 demonstrates a uniquely high LFD within the network representing “Intracellular Membrane Trafficking and Ubiquitin-Mediated Transport” (see Figure 4e). Unlike classical centrality metrics, LFD captures the local structural complexity around a node, quantifying the density and scaling behavior of its immediate topological environment. SNAP25 resides in a region where connectivity expands rapidly with distance: from just 2 direct neighbors to 29 at two steps and 373 at three steps corresponding to an exponential neighborhood growth (see Figure 4f). This fractal-like embedding supports SNAP25’s function in vesicle trafficking and membrane fusion, processes that require tight coordination across multiple functional scales within the endomembrane system. While classically associated with synaptic function, SNAP25 also plays broader roles in intracellular vesicle docking and transport, aligning with its structural position in this network.^34^

To infer the functional relevance of genes in understudied or ambiguous contexts, network perturbation analysis was used to identify structurally sensitive nodes whose importance may be concealed under static conditions. Genes of particular interest in this framework are those with low degree, low baseline values in centrality or complexity measures, but high variance under minimal edge rewiring. This pattern suggests that the gene’s topological position is highly responsive to minor structural changes in the network, indicating latent importance in specific contexts and potentially reflecting gaps in existing biological knowledge or annotation. Such genes may serve context-specific roles that are easily disrupted or overlooked in bulk analyses, making topological variance a useful proxy for identifying underexplored functional relevance.

WFDC21P (see Figure 5), a pseudogene-derived long non-coding RNA (lncRNA) with minimal annotation, shows low betweenness centrality but high betweenness variance in “Hematopoietic Stem Cell Differentiation and Leukemogenesis”. This suggests a structurally unstable role, potentially acting as a transient bridge between regulatory modules. As a member of the WFDC family, known for roles in immune modulation and cancer^35–37^, WFDC21P may function as a non-coding regulator, such as a miRNA sponge or scaffold for RNA–protein complexes. In line, WFDC21P has been reported to impact on glycolysis and STAT3 signaling in cancer.^38^ Its topological volatility may imply latent involvement in epigenetic and transcriptional regulation or the modulation of metabolic control during differentiation or transformation.

**Figure 5:**
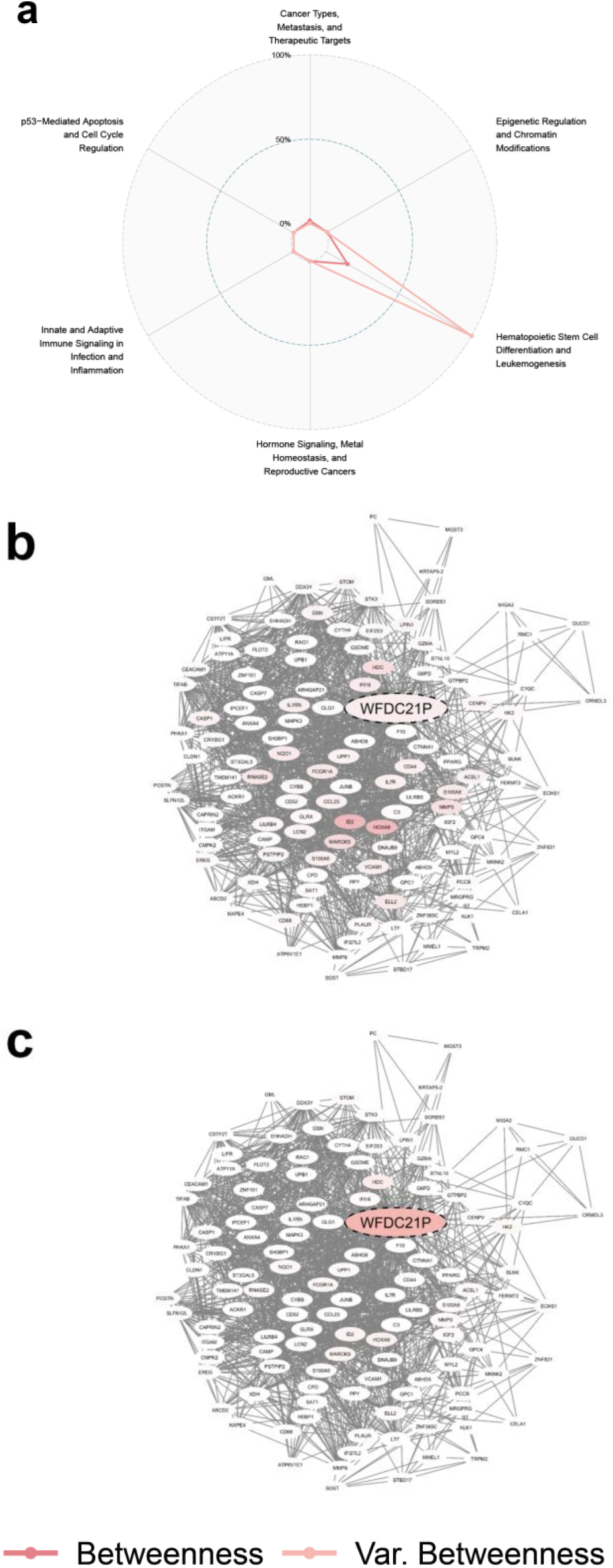
Topological variance reveals latent functional relevance of structurally sensitive genes. While conventional centrality metrics highlight statically embedded hubs, variance-based network perturbation analysis identifies genes whose importance is structurally unstable — low in static centrality but highly responsive to subtle changes in network topology. This pattern suggests potential functional relevance in specific or under-annotated contexts. (**a–c**) WFDC21P, a pseudogene-derived lncRNA, exhibits low betweenness centrality but high betweenness variance in the network “Hematopoietic Stem Cell Differentiation and Leukemogenesis”. (**a**) Radar plot shows metric and variance values across all networks; (**b**) network embedding of WFDC21P colored by betweenness; (**c**) same network colored by variance of betweenness under edge rewiring.

To facilitate broad access to the contextual and topological insights generated by this framework, all analyses have been integrated into CENTRA, an interactive web browser application designed to support exploratory investigation. CENTRA enables researchers to navigate topic-specific networks, inspect the modular architecture of functional annotations, and examine topological properties of individual genes across both well-characterized and understudied contexts. Additionally, the app allows users to display global network characteristics, browse enrichment tables linking modules to functional terms from GO^24,25^, Reactome^17^, and KEGG^18^. At the gene level, comprehensive topology metrics can be filtered, visualized, and downloaded for all genes across all networks. Furthermore, high-resolution radar plots summarizing gene-specific topological profiles can be interactively generated, customized by selected metrics and networks, and exported for external use. By making these layers of information accessible through an intuitive interface, CENTRA aims to support hypothesis generation, guide experimental prioritization, and encourage a more nuanced understanding of gene function as it emerges from literature-derived, context-aware network structures.

## Discussion

This study presents a scalable and interpretable framework for modeling the contextual relevance of genes based on literature-derived, topic-specific co-occurrence networks. By constructing networks where edges reflect shared occurrence across curated gene sets, and by calculating graph metrics such as centrality, fractality, and perturbation robustness, the approach systematically captures how genes contribute to distinct biological themes. The analysis demonstrates that well-established regulators exhibit interpretable topological signatures, while topological variance highlights potentially understudied genes with likely context-specific roles. To make these results accessible, the browser application CENTRA (Centrality-based Exploration of Network Topologies from Regulatory Assemblies) enables researchers to interactively explore the networks, functional modules, and gene-level metrics presented here, and to generate new hypotheses beyond the original analysis. Unlike conventional approaches, like ORA and GSEA, which operate on static gene lists and predefined annotations, this framework and its implementation in CENTRA offer a dynamic, topology-aware, and context-sensitive view on potential functions of single genes and proteins.

LDA is a generative probabilistic model designed to identify hidden (i.e., latent) thematic structures within large collections of data. Originally developed for topic modeling in natural language processing, LDA assumes that documents are mixtures of topics, and that topics are distributions over words.^15^ By inferring these latent topic structures, LDA can reveal semantic groupings without requiring prior labeling or supervision. In biomedical research, LDA has been applied to organize and analyze large bodies of scientific literature, allowing for the discovery of coherent biological themes across abstracts, full texts, or curated document sets.^39–42^ In the present analysis, LDA was employed to cluster the abstracts and descriptions associated with MSigDB C2 gene sets, thereby inferring biologically meaningful topics based on the semantic content of gene set documents rather than pre-assigned functional labels.

More recently, LDA and related topic modeling approaches have been adapted to single-cell transcriptomic data. Here, the aim is to uncover latent expression programs that define distinct cellular states, differentiation trajectories, or disease microenvironments.^43–45^ By interpreting cells as distributions over latent expression programs, and genes as features contributing to these programs (rather than documents as distributions over word topics), LDA can uncover hidden biological processes that govern cellular states, developmental trajectories, or pathological transitions. Applying LDA to gene set-associated literature in this study and integrating the resulting latent topics into network-based analyses offers a unique advantage: it captures a structured, scalable view of biological context without relying on fixed pathway definitions. Through this approach, CENTRA enables researchers to explore gene function within dynamically inferred thematic landscapes, bridging the strengths of probabilistic topic modeling and topological network analysis.

While LDA provides a useful probabilistic framework for clustering based on textual content, it is not the only method available for organizing functional biological information into coherent themes. Semantic similarity measures, particularly those based on GO^24,25^, offer an alternative approach by quantifying the functional relatedness between biological terms based on the structure of curated ontologies. GoSemSim^26^ is a widely used tool for this purpose. It computes the semantic similarity between GO terms by analyzing their position within the hierarchical GO graph and measuring the degree of shared ancestry between terms. Rather than relying on direct gene overlap, GoSemSim^26^ evaluates how much biological meaning is shared between two processes based on their common ancestors, thus allowing for the comparison of distinct but related annotations.

In the present study, GoSemSim^26^ was employed to map the functional relationships between topic-specific networks. Specifically, for each topic, the most significantly enriched GO:BP terms were extracted from module-level annotations, and pairwise semantic similarities were calculated. Multidimensional scaling of these similarities allowed the construction of a semantic landscape that visualizes how topics are positioned relative to one another based on their biological content. This analysis was performed to illustrate that the topic-specific gene co-occurrence networks, constructed via latent topic modeling and gene set co-occurrence, lead to functionally distinct assemblies. The centroids representing each topic were computed based on the semantic coordinates of their top enriched GO terms, providing a summary view of the biological coherence and separation achieved by the underlying network construction.

While GoSemSim^26^ provides an effective means to compare the semantic proximity of biological processes, the present framework and CENTRA extend beyond purely functional similarity. By embedding genes into topic-specific co-occurrence networks and analyzing their topological properties, the analysis captures not only what biological processes are enriched, but also how individual genes are structurally organized within these contexts. This integration of functional annotation and network architecture enables a deeper, context-aware exploration of gene function that cannot be achieved through semantic similarity measures alone.

Other tools have previously explored the intersection of literature mining and gene function interpretation yet differ notably in scope and resolution. GeneTopics^46^, for instance, applies topic modeling to gene-associated literature to identify dominant semantic themes and assign relevancy scores, offering an alternative to curated annotations through automated topic-based summaries. However, its primary focus lies in summarizing gene sets through representative keywords and literature, without modeling network structure or providing single-gene metrics. Similarly, the Gene-set Cohesion Analysis Tool (GCAT)^47^ employs Latent Semantic Indexing to assess the functional coherence of gene sets, generating cohesion scores based on textual similarity. While it displays network graphs, its core utility lies in measuring the overall functional coherence of the gene set as a whole, rather than providing granular topological metrics—such as centrality or fractality—for individual genes to characterize their precise roles within distinct, topic-specific contexts. While effective for evaluating overall group consistency, GCAT does not offer structural insights at the gene level or explore context-dependent roles across multiple biological themes.

Another framework for deriving gene-function relations, ProtSemNet^48^, constructs bipartite semantic networks with proteins and topics as vertices to evaluate protein group coherence, capturing topic-level bridging patterns qualitatively. Yet, it lacks the granularity needed to assess the topological relevance of individual genes within diverse functional contexts, as its primary metrics are focused on the overall coherence of a gene set as measured by shortest-distance subgraphs rather than a suite of systematic node-level topological measures for individual genes. In contrast, CENTRA directly embeds genes into topic-specific co-occurrence networks, quantifies their topological roles using a comprehensive set of graph metrics, and integrates this with functional enrichment to support hypothesis generation at single-gene resolution. This combination of semantic modeling and structural profiling offers a flexible and interpretable framework for exploring gene function beyond the group-level focus of earlier approaches.

Although the CENTRA framework uses modular enrichment on networks constructed from annotated gene sets, the analytical approach does not reiterate or recycle the original annotations. Instead, it introduces two independent abstraction layers: first, gene sets are clustered based on the semantic content of their reporting abstracts using unsupervised topic modeling, not by curated functional categories; second, co-occurrence networks are formed purely from structural gene intersections within each topic, without reference to any annotation. Louvain clustering is then applied to the resulting graphs—again independently of functional input—and enrichment is conducted only afterward using a distinct set of knowledge resources (GO^24,25^, Reactome^17^, KEGG^18^). Functional patterns thus emerge post hoc from structurally inferred gene modules and reflect the organization of genes in a latent, topic-specific context. This decoupling between source, structure, and annotation allows the recovery of both known and novel associations without direct reuse of the original semantic content.

While the integration of functional annotation with topic-specific co-occurrence networks offers a scalable view of gene contextuality, some limitations of the current framework must be acknowledged. First, the current analysis is based on gene sets and abstracts from curated databases, limiting coverage to documented biological processes. Although incorporating additional MSigDB^11–12^ collections such as Hallmark, Immune Signatures, or GO-derived sets could broaden thematic diversity, the resulting networks would still be constrained by literature availability and reporting bias. However, the CENTRA framework is designed to uncover latent, hidden structures within the existing body of knowledge, supporting hypothesis generation even in contexts where explicit annotations are sparse. Second, topological metric robustness was assessed by random edge rewiring, capturing general sensitivity of node-level betweenness centrality but not modeling regulatory changes such as differential expression, alternative splicing, or post-translational modification. More biologically realistic perturbation, such as introducing pseudo-nodes, assigning edge weights, or using directed interactions, could better reflect biological dynamics. Yet, such modifications would fundamentally alter the intuitive simplicity of topic-specific gene co-occurrence networks, potentially obscuring their accessibility for exploratory hypothesis generation, which remains a primary aim of the present framework. Third, the quality of gene set annotations is heterogeneous. To reduce semantic noise, the analysis included only those abstracts that could be confidently assigned to topics. While this approach inevitably excludes some biologically meaningful but ambiguously described gene sets, stricter inclusion criteria were favored to maximize clarity and robustness in the interpretation of topological measures across topics. By emphasizing precision over inclusivity, the resulting networks better support coherent functional interpretation at the cost of descriptive depth. Fourth, empirical validation and integration of experimental data remain open challenges. The current analysis provides opportunities for plausible hypotheses regarding context-dependent gene importance, particularly for poorly annotated genes, based on their structural properties and perturbation sensitivity within topic-specific gene co-occurrence networks. However, these insights remain computational inferences and require experimental validation.

Adapting the framework to incorporate user-provided experimental data would not constitute a simple extension of CENTRA. It would necessitate the construction of fundamentally different network architectures, moving from unweighted, topic-specific gene co-occurrence networks toward weighted, data-integrated systems. Such an approach would be incommensurable with the current model, which prioritizes interpretability, intuitive navigation, and consistency with curated biological knowledge. Future developments may explore hybrid models, but CENTRA as presented here is deliberately designed as a knowledge-centric exploration platform, supporting hypothesis generation rather than direct experimental integration.

In summary, the present analysis and CENTRA establish a flexible, interpretable, and accessible framework for context-aware exploration of gene function. By integrating semantic topic modeling with topological structure, the approach introduces a novel analytic dimension that moves beyond static gene lists toward literature-derived, network-based representations of biological knowledge. As functional genomics continues to scale in complexity, tools like CENTRA offer a conceptual bridge between curated domain knowledge and data-driven discovery. Looking ahead, this framework contributes to the groundwork for integrative models that combine prior knowledge with experimental data, enabling future platforms to dynamically synthesize context, structure, and function into interpretable biological insights. CENTRA thus marks a step toward a new class of exploratory tools that treat gene function not as fixed annotation, but as an emergent property of particular biological contexts.

## Acknowledgements

Author Contributions: F.H. conceived and developed the outline of this research, implemented the codes, and performed data analysis and method evaluations with the help from R.K., A.W., L.S., M.G., W.G., O.S., and W.H. F.H. wrote the paper with the help from all other authors. A.S. and S.H. supervised the work and revised the manuscript.

## Supplementary data

Supplementary Data are available at *to be indicated upon publication*.

## Data Availability

CENTRA is available for review at: [link] (*access upon request*).

## Supplementary Information

**Figure S1:**
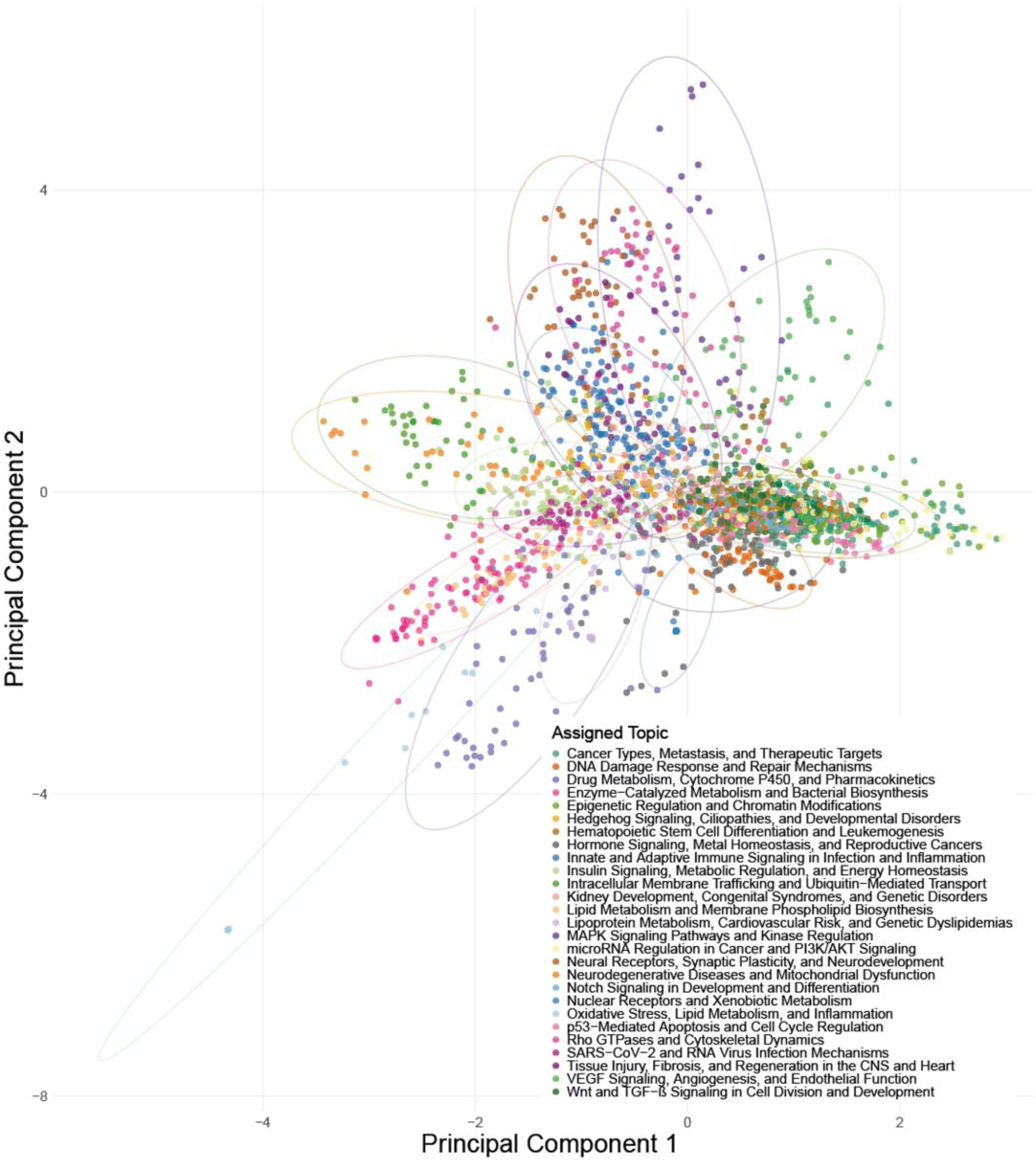
Principal Component Analysis (PCA) of the document–topic assignment matrix. Each dot in the PCA represents one document, colored by its dominant topic, and positioned based on the full probability distribution of this particular document across all 27 topics. Ellipses indicate 95% confidence intervals for each topic cluster.

**Table S1:** Topological properties of the 27 topic-specific gene co-occurrence networks. Each network was constructed by identifying shared genes across all pairwise combinations of gene sets within a given topic.

| Topic | Assigned Documents | Nodes | Edges | Avg. Degree | Clustering Coef. | Graph Density | Diameter | Modularity | Avg. Betweenness | Avg. Closeness | Avg. Path Length |
| --- | --- | --- | --- | --- | --- | --- | --- | --- | --- | --- | --- |
| Hematopoietic Stem Cell Differentiation and Leukemogenesis | 168 | 8,956 | 1,174,574 | 262.299 | 0.704 | 0.029 | 7 | 0.547 | 6488.517 | 0.002 | 2.455 |
| Neurodegenerative Diseases and Mitochondrial Dysfunction | 39 | 605 | 44,237 | 146.238 | 0.938 | 0.242 | 6 | 0.286 | 346.969 | 0.032 | 2.241 |
| Cancer Types, Metastasis, and Therapeutic Targets | 276 | 12,922 | 2,531,635 | 391.833 | 0.444 | 0.030 | 6 | 0.444 | 8426.451 | < 0.001 | 2.305 |
| Hedgehog Signaling, Ciliopathies, and Developmental Disorders | 51 | 743 | 29,085 | 78.291 | 0.714 | 0.106 | 8 | 0.548 | 627.747 | 0.024 | 2.817 |
| VEGF Signaling, Angiogenesis, and Endothelial Function | 47 | 592 | 13,543 | 45.753 | 0.663 | 0.077 | 6 | 0.547 | 412.260 | 0.032 | 2.510 |
| Kidney Development, Congenital Syndromes, and Genetic Disorders | 40 | 1,301 | 21,004 | 32.289 | 0.622 | 0.025 | 8 | 0.732 | 1458.610 | 0.014 | 3.311 |
| microRNA Regulation in Cancer and PI3K/AKT Signaling | 99 | 3,725 | 118,177 | 63.451 | 0.429 | 0.017 | 6 | 0.499 | 3206.227 | 0.009 | 2.758 |
| p53-Mediated Apoptosis and Cell Cycle Regulation | 107 | 5,524 | 781,842 | 283.071 | 0.751 | 0.051 | 6 | 0.538 | 3561.920 | 0.001 | 2.292 |
| Tissue Injury, Fibrosis, and Regeneration in the CNS and Heart | 40 | 935 | 8,921 | 19.082 | 0.677 | 0.020 | 8 | 0.638 | 888.759 | 0.048 | 3.218 |
| Innate and Adaptive Immune Signaling in Infection and Inflammation | 159 | 5,355 | 538,210 | 201.012 | 0.612 | 0.038 | 6 | 0.517 | 3693.405 | 0.003 | 2.389 |
| Hormone Signaling, Metal Homeostasis, and Reproductive Cancers | 99 | 4,037 | 209,396 | 103.738 | 0.728 | 0.026 | 8 | 0.618 | 3324.777 | 0.008 | 2.677 |
| DNA Damage Response and Repair Mechanisms | 70 | 1,736 | 142,020 | 163.618 | 0.928 | 0.094 | 6 | 0.369 | 1159.657 | 0.014 | 2.383 |
| MAPK Signaling Pathways and Kinase Regulation | 37 | 577 | 31,713 | 109.924 | 0.948 | 0.191 | 8 | 0.157 | 321.000 | 0.038 | 2.296 |
| Epigenetic Regulation and Chromatin Modifications | 121 | 7,549 | 574,897 | 152.311 | 0.523 | 0.020 | 6 | 0.593 | 5859.306 | 0.004 | 2.567 |
| Insulin Signaling, Metabolic Regulation, and Energy Homeostasis | 93 | 2,600 | 292,295 | 224.842 | 0.830 | 0.087 | 6 | 0.500 | 1785.952 | 0.002 | 2.387 |
| Rho GTPases and Cytoskeletal Dynamics | 62 | 1,397 | 189,459 | 271.237 | 0.818 | 0.194 | 5 | 0.420 | 726.631 | 0.002 | 2.044 |
| SARS-CoV-2 and RNA Virus Infection Mechanisms | 69 | 3,313 | 216,500 | 130.697 | 0.727 | 0.039 | 7 | 0.599 | 2905.826 | 0.008 | 2.784 |
| Notch Signaling in Development and Differentiation | 49 | 405 | 6,538 | 32.286 | 0.774 | 0.080 | 9 | 0.637 | 134.244 | 0.091 | 2.668 |
| Neural Receptors, Synaptic Plasticity, and Neurodevelopment | 58 | 1,428 | 177,918 | 249.185 | 0.827 | 0.175 | 7 | 0.257 | 868.653 | 0.009 | 2.261 |
| Wnt and TGF-β Signaling in Cell Division and Development | 76 | 2,301 | 75,362 | 65.504 | 0.626 | 0.028 | 7 | 0.674 | 1963.628 | 0.018 | 2.791 |
| Drug Metabolism, Cytochrome P450, and Pharmacokinetics | 53 | 300 | 5,133 | 34.220 | 0.740 | 0.114 | 8 | 0.470 | 220.933 | 0.032 | 2.603 |
| Nuclear Receptors and Xenobiotic Metabolism | 17 | 336 | 16,067 | 95.637 | 0.875 | 0.285 | 4 | 0.399 | 132.202 | 0.002 | 1.789 |
| Lipid Metabolism and Membrane Phospholipid Biosynthesis | 47 | 312 | 5,819 | 37.301 | 0.717 | 0.120 | 9 | 0.563 | 337.337 | 0.021 | 3.362 |
| Lipoprotein Metabolism, Cardiovascular Risk, and Genetic Dyslipidemias | 19 | 48 | 248 | 10.333 | 0.857 | 0.220 | 2 | 0.535 | 1.458 | 0.122 | 1.220 |
| Intracellular Membrane Trafficking and Ubiquitin-Mediated Transport | 51 | 789 | 23,158 | 58.702 | 0.774 | 0.074 | 8 | 0.702 | 886.009 | 0.013 | 3.404 |
| Oxidative Stress, Lipid Metabolism, and Inflammation | 12 | 75 | 1,490 | 39.733 | 0.964 | 0.537 | 4 | 0.098 | 24.960 | 0.009 | 1.675 |
| Enzyme-Catalyzed Metabolism and Bacterial Biosynthesis | 93 | 596 | 10,821 | 36.312 | 0.953 | 0.061 | 9 | 0.653 | 344.240 | 0.055 | 3.652 |

## References

1. Eguchi Y, Bilolikar G, Geiler-Samerotte K. Why and how to study genetic changes with context-dependent effects. Curr Opin Genet Dev. 2019 Oct;58-59:95-102. doi: 10.1016/j.gde.2019.08.003.

2. Banerji A. An attempt to construct a (general) mathematical framework to model biological "context-dependence". Syst Synth Biol. 2013;7(4): 221–227. doi: 10.1007/s11693-013-9122-6.

3. Harris CC. Structure and function of the p53 tumor suppressor gene: clues for rational cancer therapeutic strategies. J Natl Cancer Inst. 1996 Oct 16;88(20):1442–55. doi: 10.1093/jnci/88.20.1442.

4. Wang H, Guo M, Wei H, et al. Targeting p53 pathways: mechanisms, structures, and advances in therapy. Signal Transduct Target Ther. 2023 Mar 1;8(1):92. doi: 10.1038/s41392-023-01347-1.

5. Dittmer D, Pati S, Zambetti G, Chu S, Teresky AK, Moore M, Finlay C, Levine AJ. Gain of function mutations in p53. Nat Genet. 1993 May;4(1):42–6. doi: 10.1038/ng0593-42.

6. Shi Y, Ren X, Cao S, et al. *TP53* gain-of-function mutation modulates the immunosuppressive microenvironment in non-HPV-associated oral squamous cell carcinoma. J Immunother Cancer. 2023 Aug;11(8):e006666. doi: 10.1136/jitc-2023-006666.

7. Zhao M, Wang T, Gleber-Netto FO, et al. Mutant p53 gains oncogenic functions through a chromosomal instability-induced cytosolic DNA response. Nat Commun. 2024 Jan 2;15(1):180. doi: 10.1038/s41467-023-44239-2.

8. Garcia-Moreno A, López-Domínguez R, Villatoro-García JA, et al. Functional Enrichment Analysis of Regulatory Elements. Biomedicines. 2022 Mar 3;10(3):590. doi: 10.3390/biomedicines10030590.

9. Draghici S, Khatri P, Martins RP, et al. Global functional profiling of gene expression. Genomics. 2003 Feb;81(2):98–104. doi: 10.1016/s0888-7543(02)00021-6.

10. Subramanian A, Tamayo P, Mootha VK, et al. Gene set enrichment analysis: a knowledge-based approach for interpreting genome-wide expression profiles. Proc Natl Acad Sci U S A. 2005 Oct 25;102(43):15545–50. doi: 10.1073/pnas.0506580102.

11. Liberzon A, Subramanian A, Pinchback R, et al. Molecular signatures database (MSigDB) 3.0. Bioinformatics. 2011 Jun 15;27(12):1739–40. doi: 10.1093/bioinformatics/btr260.

12. Liberzon A, Birger C, Thorvaldsdóttir H, et al. The Molecular Signatures Database (MSigDB) hallmark gene set collection. Cell Syst. 2015 Dec 23;1(6):417–425. doi: 10.1016/j.cels.2015.12.004.

13. Dinsa EF, Das M, Abebe TU. A topic modeling approach for analyzing and categorizing electronic healthcare documents in Afaan Oromo without label information. Sci Rep. 2024 Dec 30;14(1):32051. doi: 10.1038/s41598-024-83743-3.

14. Grubbs AE, Sinha N, Garg R, et al. Use of topic modeling to assess research trends in the journal Gynecologic Oncology. Gynecol Oncol. 2023 May;172:41–46. doi: 10.1016/j.ygyno.2023.03.001.

15. Blei D, Ng AY, Jordan MI. Latent Dirichlet Allocation. J Mach Learn Res. 2003; 3: 993–1022. Doi: 10.5555/944919.944937.

16. Hagg LJ, Merkouris SS, O’Dea GA, et al. Examining Analytic Practices in Latent Dirichlet Allocation Within Psychological Science: Scoping Review. J Med Internet Res. 2022 Nov 8;24(11):e33166. doi: 10.2196/33166.

17. Milacic M, Beavers D, Conley P, et al. The Reactome Pathway Knowledgebase 2024. Nucleic Acids Res. 2024 Jan 5;52(D1):D672–D678. doi: 10.1093/nar/gkad1025.

18. Kanehisa M, Goto S. KEGG: kyoto encyclopedia of genes and genomes. Nucleic Acids Res. 2000 Jan 1;28(1):27–30. doi: 10.1093/nar/28.1.27.

19. Agrawal A, Balcı H, Hanspers K, et al. WikiPathways 2024: next generation pathway database. Nucleic Acids Res. 2024 Jan 5;52(D1):D679–D689. doi: 10.1093/nar/gkad960.

20. Feinerer I, Hornik K, Meyer D. Text Mining Infrastructure in R. J Stat Softw. 2008; 25(5):1–54. doi: doi:10.18637/jss.v025.i05.

21. Csardi G, Nepusz T. The igraph software package for complex network research. InterJournal. 2006; Complex Systems:1695.

22. McGraw PN, Menzinger M. Laplacian spectra as a diagnostic tool for network structure and dynamics. Phys Rev. 2008; 77(3): 0311002. doi: 10.1103/PhysRevE.77.031102.

23. Kolberg L, Raudvere U, Kuzmin I, et al. g:Profiler-interoperable web service for functional enrichment analysis and gene identifier mapping (2023 update). Nucleic Acids Res. 2023 Jul 5;51(W1):W207–W212. doi: 10.1093/nar/gkad347.

24. Ashburner M, Ball CA, Blake JA, et al. Gene ontology: tool for the unification of biology. Nat Genet. 2000 May;25(1):25–29. doi: 10.1038/75556.

25. The Gene Ontology Consortium. The Gene Ontology knowledgebase in 2023. Genetics. 2023 May 4;224(1):iyad031. doi: 10.1093/genetics/iyad031.

26. Yu G. Gene Ontology Semantic Similarity Analysis Using GOSemSim. Methods Mol Biol. 2020;2117:207–215. doi: 10.1007/978-1-0716-0301-7_11.

27. Xiao X, Chen H, Bogdan P. Deciphering the generating rules and functionalities of complex networks. Sci Rep. 2021 Nov 25;11(1):22964. doi: 10.1038/s41598-021-02203-4.

28. Ghorbani M, Jonckheere EA, Bogdan P. Gene Expression Is Not Random: Scaling, Long-Range Cross-Dependence, and Fractal Characteristics of Gene Regulatory Networks. Front Physiol. 2018 Oct 22;9:1446. doi: 10.3389/fphys.2018.01446.

29. Ozgür A, Vu T, Erkan G, et al. Identifying gene-disease associations using centrality on a literature mined gene-interaction network. Bioinformatics. 2008 Jul 1;24(13):i277–85. doi: 10.1093/bioinformatics/btn182.

30. Hu Q, Zhang Y, Lou H, et al. GPX4 and vitamin E cooperatively protect hematopoietic stem and progenitor cells from lipid peroxidation and ferroptosis. Cell Death Dis. 2021 Jul 15;12(7):706. doi: 10.1038/s41419-021-04008-9.

31. Savaskan NE, Borchert A, Bräuer AU, Kuhn H. Role for glutathione peroxidase-4 in brain development and neuronal apoptosis: specific induction of enzyme expression in reactive astrocytes following brain injury. Free Radic Biol Med. 2007 Jul 15;43(2):191–201. doi: 10.1016/j.freeradbiomed.2007.03.033.

32. Galvagni F, Pennacchini S, Salameh A, et al. Endothelial cell adhesion to the extracellular matrix induces c-Src-dependent VEGFR-3 phosphorylation without the activation of the receptor intrinsic kinase activity. Circ Res. 2010 Jun 25;106(12):1839–48. doi: 10.1161/CIRCRESAHA.109.206326.

33. Patel A, Sabbineni H, Clarke A, et al. Novel roles of Src in cancer cell epithelial-to-mesenchymal transition, vascular permeability, microinvasion and metastasis. Life Sci. 2016 Jul 15;157:52–61. doi: 10.1016/j.lfs.2016.05.036.

34. Zhao Y, Fang Q, Sharma S, Jakhanwal S, Jahn R, Lindau M. All SNAP25 molecules in the vesicle-plasma membrane contact zone change conformation during vesicle priming. Proc Natl Acad Sci USA. 2024 Jan 9;121(2):e2309161121. doi: 10.1073/pnas.2309161121.

35. Bingle CD, Vyakarnam A. Novel innate immune functions of the whey acidic protein family. Trends Immunol. 2008 Sep;29(9):444–53. doi: 10.1016/j.it.2008.07.001.

36. Larsen M, Ressler SJ, Gerdes MJ, et al. The WFDC1 gene encoding ps20 localizes to 16q24, a region of LOH in multiple cancers. Mamm Genome. 2000 Sep;11(9):767–73. doi: 10.1007/s003350010135.

37. Bingle L, Cross SS, High AS, et al. WFDC2 (HE4): a potential role in the innate immunity of the oral cavity and respiratory tract and the development of adenocarcinomas of the lung. Respir Res. 2006 Apr 6;7(1):61. doi: 10.1186/1465-9921-7-61.

38. Guan YF, Huang QL, Ai YL, Chen QT, Zhao WX, Wang XM, Wu Q, Chen HZ. Nur77-activated lncRNA WFDC21P attenuates hepatocarcinogenesis via modulating glycolysis. Oncogene. 2020 Mar;39(11):2408–2423. doi: 10.1038/s41388-020-1158-y.

39. Kavvadias S, Drosatos G, Kaldoudi E. Supporting topic modeling and trends analysis in biomedical literature. J Biomed Inform. 2020 Oct;110:103574. doi: 10.1016/j.jbi.2020.103574.

40. Lu A, Li K, Su G, et al. Revealing Academic Evolution and Frontier Pattern in the Field of Uveitis Using Bibliometric Analysis, Natural Language Processing, and Machine Learning. Ocul Immunol Inflamm. 2024 Oct;32(8):1564–1579. doi: 10.1080/09273948.2023.2262028.

41. Nguyen TT, Nguyen HT, Do HP, et al. Characterizing the Development of Research Landscapes in Substance Use and HIV/AIDS During 1990 to 2021. Subst Abuse. 2023 Jun 5;17:11782218231177515. doi: 10.1177/11782218231177515.

42. Montes-Escobar K, de la Hoz-M J, Castillo-Cordova P, et al. Glioblastoma: a comprehensive approach combining bibliometric analysis, Latent Dirichlet Allocation, and HJ-Biplot: Glioblastoma insights and trends: a 49-year bibliometric analysis. Neurosurg Rev. 2024 May 10;47(1):209. doi: 10.1007/s10143-024-02440-x.

43. Pancheva A, Wheadon H, Rogers S, et al. Using topic modeling to detect cellular crosstalk in scRNA-seq. PLoS Comput Biol. 2022 Apr 8;18(4):e1009975. doi: 10.1371/journal.pcbi.1009975.

44. Adossa NA, Rytkönen KT, Elo LL. Dirichlet process mixture models for single-cell RNA-seq clustering. Biol Open. 2022 Apr 15;11(4):bio059001. doi: 10.1242/bio.059001.

45. Wu X, Teo YV, Neretti N, et al. Mouse blood cells types and aging prediction using penalized Latent Dirichlet Allocation. BMC Genomics. 2024 Sep 18;23(Suppl 4):866. doi: 10.1186/s12864-024-10763-8.

46. Wang V, Xi L, Enayetallah A, et al. GeneTopics—interpretation of gene sets via literature-driven topic models. BMC Syst Biol. 2013;7 Suppl 5(Suppl 5):S10. doi: 10.1186/1752-0509-7-S5-S10.

47. Xu L, Furlotte N, Lin Y, et al. Functional cohesion of gene sets determined by latent semantic indexing of PubMed abstracts. PLoS One. 2011 Apr 14;6(4):e18851. doi: 10.1371/journal.pone.0018851.

48. Zheng B, Lu X. Novel metrics for evaluating the functional coherence of protein groups via protein semantic network. Genome Biol. 2007;8(7):R153. doi: 10.1186/gb-2007-8-7-r153.

