## Supplementary File S1 for "CENTRA: Knowledge-Based Gene Contexuality Graphs Reveal Functional Master Regulators by Centrality and Fractality"

| | | | | |
|---|---|---|---|---|
| 14576184 | 16121031 | 15033539 | 16116475 | 17252012 |
| 20584986 | 15657362 | 12137940 | 17234778 | 17450142 |
| 17001320 | 15688035 | 17952126 | 12414654 | 18948947 |
| 12471561 | 17483311 | 14525773 | 19029983 | 15674340 |
| 12197474 | 33428749 | 15711547 | 18710942 | 12900543 |
| 15388584 | 12642603 | 18849966 | 18794116 | 16551631 |
| 20085707 | 18443585 | 18454173 | 18172325 | 16491124 |
| 18413731 | 17213802 | 19061838 | 17234794 | 17119049 |
| 18838534 | 17353267 | 15735755 | 18199539 | 17234770 |
| 15252187 | 18757432 | 16501609 | 16877703 | 17548472 |
| 16611997 | 18490921 | 18316590 | 18272964 | 15681441 |
| 15592509 | 16651414 | 12379459 | 16314830 | 14604967 |
| 15199222 | 17234781 | 18710938 | 10737792 | 11830491 |
| 17486063 | 15070671 | 16990782 | 15592499 | 18701473 |
| 17043641 | 18838536 | 16909116 | 16983338 | 17636019 |
| 17287851 | 16397028 | 16314847 | 18849962 | 16652140 |
| 15831697 | 18339861 | 15856024 | 34197623 | 18757399 |
| 17621274 | 17562867 | 18212061 | 17344918 | 20406972 |
| 20123981 | 16617321 | 18519664 | 18506891 | 17072329 |
| 17409456 | 16273092 | 17430594 | 17704800 | 17525748 |
| 16116230 | 14769913 | 17242180 | 17213801 | 16862181 |
| 18381438 | 32416070 | 10756030 | 18794137 | 18172260 |
| 17322878 | 17409441 | 18676852 | 14676830 | 11741835 |
| 15824734 | 18660816 | 16288205 | 12791645 | 16007187 |
| 22208948 | 16341039 | 17404577 | 17043659 | 14522256 |
| 21320499 | 17195838 | 12480690 | 17409444 | 16565084 |
| 15756019 | 18931683 | 17146443 | 16207825 | 18381450 |
| 17213814 | 18593951 | 21364935 | 15608684 | 16434974 |
| 11384963 | 15635089 | 16424048 | 18381411 | 21873988 |
| 16954472 | 16862182 | 18339859 | 17709385 | 15817677 |
| 12393420 | 15674348 | 12915101 | 14519204 | 17724461 |
| 15303102 | 19414752 | 17242199 | 17525749 | 20624283 |
| 18264110 | 20336062 | 16107850 | 15095275 | 19832978 |
| 16331256 | 20956565 | 15757903 | 15735721 | 15897907 |
| 17438134 | 17016442 | 18344982 | 16849537 | 12695333 |
| 12479369 | 17187432 | 14573703 | 12947006 | 12086890 |
| 15042541 | 18413726 | 12824457 | 17260020 | 12637319 |
| 15520196 | 17483317 | 18923524 | 12771951 | 15958647 |
| 19010895 | 12447701 | 14981515 | 12885910 | 17952124 |
| 17934481 | 18849970 | 12704389 | 17072333 | 17785439 |
| 17353275 | 16799620 | 15711545 | 19884340 | 20060365 |
| 16331246 | 20421412 | 18701503 | 15870273 | 18438415 |
| 17704777 | 12969979 | 16607279 | 11423116 | 18245477 |
| 15980968 | 11711622 | 16160012 | 17452451 | 18794900 |
| 15806164 | 16769770 | 14684422 | 17621275 | 27280975 |
| 16091735 | 17308078 | 15336447 | 19189975 | 15307835 |
| 17043639 | 18195040 | 15452378 | 17546029 | 25721503 |
| 17072343 | 18288184 | 20385362 | 9861020 | 16627760 |
| 15778709 | 14701746 | 17001317 | 28498607 | 17968324 |
| 15331443 | 18157142 | 17409404 | 17713554 | 28119430 |
| 14707115 | 20087356 | 16909099 | 18339860 | 20421419 |

| | | | | |
|---|---|---|---|---|
| 20713442 | 16607286 | 12673201 | 18724357 | 15940259 |
| 17538624 | 18317448 | 20385360 | 17178841 | 17623797 |
| 16849555 | 17726467 | 17170726 | 18451147 | 18026104 |
| 14973112 | 17173069 | 12411303 | 12228721 | 18762803 |
| 16849584 | 17210682 | 20655465 | 18212054 | 15033468 |
| 18427549 | 17160024 | 11779835 | 17086209 | 15213097 |
| 18711403 | 17717066 | 20124474 | 18281487 | 18227149 |
| 18316615 | 11991981 | 11468180 | 16288221 | 14499499 |
| 18574468 | 16007192 | 12411319 | 14508521 | 18757430 |
| 18710934 | 17210724 | 15295046 | 18519682 | 16116477 |
| 16785517 | 15310760 | 29346386 | 16210406 | 17060456 |
| 15735688 | 20609354 | 16585179 | 11137995 | 16885346 |
| 16532023 | 17440070 | 18724378 | 17289871 | 17682065 |
| 20308323 | 17704799 | 16998498 | 16715129 | 17430890 |
| 18591254 | 15897887 | 16474848 | 15578091 | 20584988 |
| 12198119 | 28446439 | 18413733 | 12096084 | 17114343 |
| 16105978 | 18037961 | 17088532 | 15592526 | 12531789 |
| 17998939 | 20385766 | 14512514 | 14562044 | 16205638 |
| 12824169 | 12791650 | 18923165 | 19047146 | 17145885 |
| 18632613 | 19596788 | 19723656 | 17928865 | 16186804 |
| 16892093 | 17252022 | 32663312 | 11172053 | 19364821 |
| 17039238 | 17483333 | 16612386 | 17922013 | 16247484 |
| 18591936 | 16044152 | 16278680 | 12692265 | 18955504 |
| 16912112 | 15331438 | 17009164 | 17440068 | 16908595 |
| 17878328 | 16547494 | 11773596 | 28855211 | 16462763 |
| 18632645 | 16914566 | 17332353 | 15031206 | 18488029 |
| 17130830 | 15205324 | 18593902 | 17676035 | 16247469 |
| 18381431 | 16983347 | 19620289 | 15386376 | 16909124 |
| 17220874 | 18381439 | 17468757 | 15750623 | 15870693 |
| 16741228 | 15492243 | 16760442 | 17998334 | 16424014 |
| 17960246 | 12354774 | 16247483 | 18246078 | 18985860 |
| 11487589 | 18337816 | 18059283 | 15318170 | 12036489 |
| 12907719 | 19074870 | 17409405 | 17486088 | 16314831 |
| 21177505 | 11902584 | 16932341 | 15103417 | 19858293 |
| 19074876 | 18542059 | 15592519 | 11313881 | 18701491 |
| 12021175 | 16205646 | 15120960 | 20478528 | 18690214 |
| 15985538 | 15459216 | 16799640 | 16293602 | 18451145 |
| 16501597 | 16407832 | 33096892 | 18838539 | 20211141 |
| 19108712 | 23636981 | 12663442 | 14734480 | 14615372 |
| 18701499 | 12610208 | 17297441 | 19805515 | 15475267 |
| 12438227 | 14625302 | 15146237 | 15579653 | 15288478 |
| 17072321 | 15688027 | 12648972 | 11402317 | 18042934 |
| 16899599 | 16732325 | 15710396 | 15592527 | 17332321 |
| 33829040 | 17483324 | 19188439 | 16710476 | 19289500 |
| 15931389 | 18172295 | 19564420 | 16619041 | 15897889 |
| 12433680 | 14673169 | 16007138 | 15769904 | 14527998 |
| 19139271 | 17099726 | 12446780 | 17470548 | 12606941 |
| 17873875 | 17533367 | 17873912 | 19307312 | 16819509 |
| 15548615 | 16507773 | 12234922 | 17891185 | 12914697 |
| 10521349 | 17237784 | 11416145 | 11309484 | 18676831 |
| 18075594 | 19074878 | 17101797 | 15642117 | 16434967 |

| | | | | |
|---|---|---|---|---|
| 15897883 | 18039849 | 16998502 | 12213819 | 16166251 |
| 15608679 | 20538915 | 16357323 | 17597811 | 15940270 |
| 17533364 | 15657340 | 17363560 | 12969976 | 18281475 |
| 17369842 | 12057921 | 15735737 | 11532376 | 18381418 |
| 16449969 | 15531917 | 18989307 | 17237808 | 16799638 |
| 10888876 | 18391978 | 16551867 | 16751803 | 17891175 |
| 10464095 | 18794111 | 12130493 | 18413734 | 16774935 |
| 17515606 | 16540645 | 17486082 | 16936776 | 17873882 |
| 15210650 | 18223687 | 18339881 | 11027337 | 15485929 |
| 15565109 | 19451229 | 14566342 | 17142811 | 17967875 |
| 16532004 | 16007190 | 18794130 | 17694077 | 16741146 |
| 15349906 | 17293870 | 12540486 | 15735679 | 17483347 |
| 18245461 | 15608688 | 18632628 | 11493461 | 12893766 |
| 18037878 | 16568090 | 18641651 | 12646203 | 11781229 |
| 17452975 | 15190254 | 15869869 | 16491115 | 19433444 |
| 17151600 | 20708159 | 18500333 | 17210693 | 17339329 |
| 15699330 | 15591113 | 27191501 | 17699763 | 18066064 |
| 15105423 | 17308118 | 15731117 | 18071306 | 16953217 |
| 11279020 | 10741968 | 15558012 | 17464315 | 15735714 |
| 12885785 | 26763932 | 18600261 | 16849564 | 15738394 |
| 15695336 | 10644742 | 18288197 | 16140920 | 20074520 |
| 19635812 | 19332562 | 16507782 | 15674343 | 19364815 |
| 18451154 | 17189429 | 16785998 | 12185249 | 20479124 |
| 17220891 | 12623842 | 18285459 | 12766259 | 18829567 |
| 11861364 | 17325667 | 17440110 | 16772382 | 17287849 |
| 16127159 | 12091409 | 18509334 | 17440165 | 17563751 |
| 11468144 | 15824735 | 17603471 | 16618808 | 18443592 |
| 18504433 | 19917725 | 18829530 | 15339848 | 12198161 |
| 17603561 | 17409432 | 18451862 | 19010930 | 18644872 |
| 20346151 | 29625051 | 18519693 | 17297461 | 20516219 |
| 16532037 | 15374877 | 15637593 | 20399149 | 18006812 |
| 12356685 | 32573711 | 18593922 | 15302897 | 16533756 |
| 12702554 | 15750621 | 18212050 | 15664994 | 18852285 |
| 17242205 | 15661559 | 16127434 | 17213809 | 17965720 |
| 11522623 | 15601821 | 20144757 | 12670911 | 19752192 |
| 18245465 | 10969808 | 15515172 | 17724462 | 17308109 |
| 18815592 | 17452985 | 11438693 | 15558013 | 11807556 |
| 19074894 | 17322921 | 17213808 | 16508012 | 15864312 |
| 17308126 | 15831586 | 18413728 | 16785988 | 18339882 |
| 18332864 | 17237765 | 18794119 | 18426912 | 16636670 |
| 16990345 | 20159609 | 17873908 | 17211412 | 16531451 |
| 16205643 | 15302890 | 12808457 | 17667943 | 17971902 |
| 20123980 | 18245467 | 15827134 | 12855579 | 17922014 |
| 16618720 | 11755394 | 14606959 | 16909110 | 19103759 |
| 11509180 | 15897246 | 18922927 | 18483246 | 17510386 |
| 19289498 | 17694086 | 17761758 | 12802282 | 20609350 |
| 18676865 | 17934524 | 18632618 | 16651410 | 17363561 |
| 11981038 | 15608685 | 16832351 | 17377532 | 25681390 |
| 16369491 | 18281483 | 15024077 | 18339843 | 11910354 |
| 17072341 | 18559539 | 16909125 | 12682234 | 16670265 |
| 18757419 | 18245484 | 18724358 | 16740760 | 18981216 |

| | | | | |
|---|---|---|---|---|
| 17353439 | 15749026 | 17213807 | 15271793 | 19564408 |
| 18438429 | 16331255 | 15735734 | 15516973 | 11279127 |
| 12228720 | 19805517 | 11016956 | 18981221 | 15531915 |
| 12469122 | 17377531 | 16007217 | 17597760 | 18381434 |
| 16909112 | 15198980 | 18829555 | 15187151 | 12198164 |
| 11070096 | 18281472 | 16467079 | 17468756 | 17173050 |
| 18490920 | 14530283 | 18451135 | 18625718 | 12618007 |
| 20837710 | 16832346 | 16540638 | 19043405 | 19158274 |
| 16314843 | 17413002 | 21844125 | 17606841 | 15637585 |
| 20547756 | 15471956 | 12591738 | 20038531 | 17767167 |
| 17934472 | 15856012 | 12717449 | 15722553 | 18039842 |
| 15155749 | 12832290 | 15824737 | 16394013 | 19414603 |
| 17283130 | 15026540 | 11867723 | 16799645 | 17297437 |
| 17483325 | 28847964 | 16478745 | 18413723 | 15189810 |
| 20404087 | 12839967 | 17906636 | 17260014 | 18245496 |
| 18701482 | 18794802 | 17206142 | 17173048 | 16467078 |
| 11799067 | 17404570 | 17334370 | 18663361 | 17954559 |
| 16449971 | 17072322 | 15548687 | 17898786 | 16449976 |
| 17330098 | 17200670 | 20227041 | 17483357 | 15107491 |
| 15184677 | 12640676 | 11937641 | 17875932 | 17452977 |
| 18679425 | 15516975 | 18469815 | 18519671 | 17486081 |
| 33263276 | 17143286 | 18172294 | 12418965 | 15755900 |
| 18974135 | 19273610 | 18285465 | 16788156 | 17392792 |
| 18381423 | 18544741 | 17443180 | 16247478 | 15721472 |
| 18316601 | 12789274 | 19687298 | 17440099 | 15735726 |
| 17213805 | 16707453 | 18974123 | 15895078 | 19703992 |
| 17334389 | 11139609 | 18676871 | 17283122 | 17440082 |
| 17406368 | 18026132 | 15033914 | 15608674 | 17438126 |
| 15562319 | 17982488 | 17001308 | 17998335 | 19047173 |
| 21159642 | 17709396 | 18378698 | 19074901 | 17975138 |
| 15741219 | 33766982 | 18542061 | 17389037 | 16751804 |
| 18264134 | 16158056 | 17403900 | 17496919 | 16247446 |
| 14973077 | 15580292 | 19047182 | 16574658 | 20541704 |
| 15226186 | 17952090 | 17599052 | 15273285 | 16135788 |
| 16007141 | 16407838 | 20231363 | 16041372 | 15635450 |
| 16247447 | 11516994 | 11992124 | 16077073 | 19307311 |
| 16702952 | 17470557 | 11904358 | 15051823 | 17440055 |
| 17728400 | 16912175 | 15608639 | 16606835 | 19580510 |
| 18519667 | 19289501 | 16116479 | 15084694 | 17047040 |
| 12732648 | 11867738 | 18762810 | 15561778 | 16585155 |
| 15492259 | 18505969 | 16103878 | 17371845 | 17255260 |
| 17724341 | 18568025 | 11982916 | 15940248 | 17334365 |
| 16527888 | 16407836 | 17297473 | 17923686 | 16007088 |
| 29045844 | 17283119 | 16818636 | 20956564 | 12419474 |
| 18171984 | 18641660 | 17438526 | 11823860 | 15897868 |
| 17334401 | 19074882 | 17533371 | 16079796 | 18504438 |
| 15834410 | 16150706 | 12663452 | 17483315 | 15297311 |
| 18413748 | 16607285 | 18185580 | 17297478 | 12032322 |
| 19074895 | 19010892 | 14517214 | 16109776 | 18235444 |
| 33429950 | 11786909 | 17363581 | 20129251 | 16006652 |
| 17500590 | 16862184 | 18772890 | 12923195 | 12200464 |

| | | | | |
|---|---|---|---|---|
| 17234769 | 18974140 | 18805970 | 18197893 | 21116052 |
| 17210681 | 17016446 | 16236521 | 18174920 | 24847263 |
| 17429401 | 12016162 | 12563308 | 16286565 | 29378302 |
| 14737109 | 18281482 | 16146838 | 12649599 | 25394486 |
| 15829979 | 15592518 | 12951584 | 17332888 | 31191253 |
| 12058064 | 16311603 | 16642045 | 19549592 | 27807401 |
| 17452456 | 17938208 | 15625120 | 15590995 | 19903023 |
| 15100412 | 20805357 | 11721960 | 12943727 | 19464758 |
| 15608677 | 16382050 | 12468433 | 11208606 | 12805290 |
| 18779318 | 17618270 | 16352814 | 17309947 | 15645263 |
| 17297452 | 12381414 | 12529654 | 10746853 | 26074767 |
| 18070364 | 18483247 | 12032780 | 12023610 | 24529521 |
| 18806801 | 17178890 | 10717473 | 17445565 | 26136647 |
| 12095419 | 20584981 | 19075268 | 14710779 | 20101724 |
| 17700529 | 12719586 | 12130514 | 8835634 | 18252769 |
| 18632634 | 18559513 | 12393465 | 10571983 | 15314242 |
| 16595783 | 18536717 | 11090075 | 10331992 | 2793832 |
| 17409455 | 15619625 | 15160934 | 10667806 | 8071227 |
| 17908789 | 14517420 | 11607818 | 12216939 | 8444803 |
| 18397753 | 14639605 | 19855079 | 12563298 | 9811644 |
| 17043662 | 19010928 | 17554387 | 17172287 | 10931327 |
| 18199530 | 15930281 | 15001769 | 15141068 | 10984043 |
| 17486072 | 12393520 | 10204118 | 17215350 | 11447132 |
| 18381945 | 16728703 | 10959047 | 16116284 | 12562791 |
| 18381452 | 11861292 | 15489912 | 15670154 | 14450717 |
| 18560354 | 17237761 | 11171368 | 14522973 | 15134748 |
| 16951684 | 18948956 | 11984870 | 15232608 | 15489439 |
| 16116430 | 18264109 | 12640114 | 15750290 | 15695810 |
| 12138103 | 12679051 | 14623871 | 18675468 | 15809294 |
| 18724390 | 17043644 | 11703922 | 9848086 | 16237198 |
| 11439330 | 18757440 | 12665527 | 18035450 | 16277754 |
| 17456585 | 17404513 | 10890911 | 25450339 | 16452451 |
| 33458693 | 18025224 | 15551862 | 29890840 | 16995900 |
| 17099727 | 18754011 | 15561584 | 25767489 | 17085508 |
| 16424387 | 12673210 | 15733744 | 11484003 | 17190829 |
| 15845533 | 9826724 | 12829246 | 15177383 | 17827295 |
| 16140955 | 17074815 | 15980064 | 17568632 | 18199744 |
| 12738660 | 15897880 | 10766253 | 22842534 | 18304669 |
| 17283147 | 21596316 | 15856064 | 21663966 | 11479276 |
| 16652150 | 22159717 | 12754525 | 16677790 | 28700839 |
| 19150350 | 9799792 | 11557972 | 14557245 | 28512398 |
| 18316609 | 18424520 | 14716305 | 24051203 | 30879475 |
| 18381416 | 18375759 | 9671776 | 30741722 | 30870681 |
| 12604609 | 10557362 | 10905472 | 10527810 | 23415570 |
| 15897875 | 11731493 | 12368907 | 31655116 | 16924260 |
| 17551146 | 16135226 | 11479627 | 18648507 | 16723044 |
| 19451226 | 11929526 | 15058305 | 20332425 | 28669745 |
| 16261164 | 18326572 | 14679176 | 16394273 | 32344665 |
| 16007176 | 16109926 | 9687539 | 11551913 | 16372325 |
| 12554760 | 17873039 | 16002531 | 28057414 | 16603792 |
| 18754008 | 17214741 | 11850323 | 27994051 | 17051205 |

| | | | | |
|---|---|---|---|---|
| 16909017 | 23803157 | 16124859 | 14556242 | 18086859 |
| 14569202 | 15654015 | 17222174 | 11736900 | 12838335 |
| 15310460 | 15725726 | 15564123 | 16054233 | 11413485 |
| 12595144 | 15322527 | 10517845 | 11130178 | 15111104 |
| 17434459 | 12838338 | 16595667 | 28286127 | 11299192 |
| 27649160 | 19148180 | 17462988 | 25766766 | 11698232 |
| 19230774 | 22429822 | 14996803 | 16858867 | 8125957 |
| 16026864 | 25526085 | 12644495 | 12516863 | 15000521 |
| 11239414 | 15601572 | 11172168 | 19308322 | 9149541 |
| 11870681 | 9894605 | 11477111 | 10838567 | 10997573 |
| 16713195 | 23070002 | 10719286 | 17893748 | 9235901 |
| 30837838 | 18304445 | 17976144 | 17337257 | 7861177 |
| 29626651 | 20533903 | 11686879 | 21665970 | 14572802 |
| 26264610 | 15719025 | 17133123 | 11436317 | 10331643 |
| 28148298 | 24278747 | 18274560 | 16564093 | 11080247 |
| 28878620 | 21145844 | 18274559 | 12482854 | 10508738 |
| 17409386 | 18023214 | 12093011 | 12087109 | 12972622 |
| 17566607 | 16397578 | 17364144 | 11500517 | 14517314 |
| 26388731 | 21914490 | 10811837 | 16766224 | 21338471 |
| 25991442 | 22465226 | 15049952 | 17719544 | 22129055 |
| 32119873 | 26231733 | 15762980 | 24184513 | 22002090 |
| 28057298 | 22075987 | 9534029 | 20018209 | 19172794 |
| 24619348 | 18238895 | 11913070 | 22158834 | 19475418 |
| 27815720 | 12949498 | 12192541 | 28784662 | 19434403 |
| 29476642 | 14737178 | 11396701 | 20937905 | 26228554 |
| 21722302 | 9916983 | 17192582 | 21959040 | 20807542 |
| 16472115 | 11520933 | 15132715 | 19826053 | 17988210 |
| 18519638 | 24769728 | 11577986 | 22562166 | 11805843 |
| 21834058 | 18208375 | 14682358 | 11734571 | 16799263 |
| 31319884 | 19770840 | 15630019 | 17158541 | 18981594 |
| 29605155 | 20075940 | 12677003 | 16474624 | 16552170 |
| 25071440 | 14636671 | 15818466 | 16110317 | 10747208 |
| 29299811 | 15292194 | 15363605 | 11103787 | 16701909 |
| 30242016 | 15205388 | 11891122 | 12850530 | 16096350 |
| 23361386 | 14660610 | 11687826 | 17149381 | 12763850 |
| 15771591 | 14626496 | 12471249 | 15746962 | 11294886 |
| 15814328 | 14622984 | 15960985 | 15551095 | 15736582 |
| 11905820 | 11679404 | 15084255 | 12461567 | 14659803 |
| 15304636 | 11328810 | 25374504 | 10358768 | 19403607 |
| 16200082 | 10908295 | 17169551 | 15886119 | 28596175 |
| 12776207 | 9812980 | 26651291 | 15752555 | 14690046 |
| 12974477 | 5288244 | 15948971 | 12033741 | 11910893 |
| 15224093 | 16499623 | 12588368 | 15364057 | 9597126 |
| 16181333 | 15590624 | 15978322 | 15771577 | 16034094 |
| 15653312 | 19205777 | 15468170 | 15886112 | 12234363 |
| 11244040 | 17413274 | 16881963 | 15457440 | 15071551 |
| 15745856 | 12270949 | 16306523 | 15322145 | 14519386 |
| 15224094 | 11716909 | 11276000 | 15626471 | 10798271 |
| 15653315 | 16823481 | 15270885 | 12119152 | 9150551 |
| 20010787 | 18382419 | 14737121 | 9697839 | 12810109 |
| 26615431 | 16101641 | 15656799 | 11163197 | 14519398 |

| | | | | |
|---|---|---|---|---|
| 15882774 | 15771581 | 25568220 | 19321146 | 9417034 |
| 11050419 | 17496928 | 26082779 | 19567913 | 17636923 |
| 11860281 | 9765254 | 21811895 | 19665965 | 11572082 |
| 10964748 | 15046611 | 25705421 | 16286919 | 12921709 |
| 12367625 | 12051620 | 23110467 | 19767723 | 12125070 |
| 10664064 | 17675482 | 17635716 | 23258413 | 12796473 |
| 14556710 | 14607270 | 26324800 | 35006429 | 7698997 |
| 15115723 | 15719031 | 23911436 | 34949589 | 10617638 |
| 15056568 | 11071626 | 24924409 | 32829024 | 15036264 |
| 15728677 | 14982876 | 20027185 | 34970269 | 15381249 |
| 12500939 | 10403855 | 23564577 | 34504660 | 11331578 |
| 15923648 | 15156182 | 23564578 | 18672201 | 12736199 |
| 7908906 | 16484590 | 25309880 | 19273156 | 15229478 |
| 8242749 | 9834202 | NA | 11805847 | 12845610 |
| 16936746 | 12801837 | 15102328 | 19017683 | 29079659 |
| 15838516 | 11160144 | 14551435 | 19246813 | 19007773 |
| 17568790 | 12244301 | 12022921 | 14522402 | 18647649 |
| 9832503 | 22286129 | 12670965 | 15146239 | 15261671 |
| 19020303 | 27990019 | 17169919 | 11524405 | 19325624 |
| 10385618 | 32226546 | 17030798 | 16817756 | 20110990 |
| 12809600 | 12015613 | 18625006 | 8087839 | 15687999 |
| 12925736 | 12397359 | 18391471 | 9759502 | 19924646 |
| 16212951 | 12724733 | 10613873 | 16038189 | 16203612 |
| 15838523 | 12507418 | 12644499 | 18043895 | 19696797 |
| 11099028 | 18802415 | 18520135 | 16550168 | 22193161 |
| 7973727 | 15000146 | 19270684 | 16393888 | 15947972 |
| 9889196 | 16555243 | 24892271 | 11226815 | 14747944 |
| 17018296 | 11078609 | 21106488 | 10575324 | 16508724 |
| 9029154 | 16699851 | 27577879 | 9698084 | 17028294 |
| 15876871 | 12621137 | 15500860 | 11504799 | 11431710 |
| 17426725 | 12951588 | 16034365 | 10901707 | 15284264 |
| 12573239 | 10505543 | 23847622 | 12399158 | 7585656 |
| 15568976 | 16042571 | 21349428 | 18463198 | 16248975 |
| 9618481 | 15349822 | 17530315 | 12386121 | 14700549 |
| 26650195 | 9196022 | 24381786 | 10859160 | 15694859 |
| 20935501 | 12737309 | 18926762 | 15788214 | 11389469 |
| 12495636 | 11459867 | 21155715 | 16048561 | 12197905 |
| 11872087 | 15573119 | 22410872 | NA | 14967450 |
| 12069927 | 15310786 | 12189384 | 15514566 | 11252954 |
| 17169327 | 11223406 | 26115564 | 10976516 | 16829981 |
| 11244042 | 15711891 | 23612460 | 15173095 | 17000658 |
| 8576262 | 11170304 | 22710174 | 11489791 | 16377102 |
| 17537990 | 16326109 | 19362700 | 11379777 | 14744244 |
| 18662984 | 15479695 | 19497760 | 24281184 | 12851486 |
| 10852130 | 11477132 | 18573338 | 27709007 | 15864276 |
| 14625285 | 19167459 | 19122387 | 27232112 | 16729045 |
| 17371236 | 18568040 | 18280611 | 20195289 | 16729043 |
| 16144840 | 24213116 | 18941233 | 20434429 | 10880430 |
| 18711432 | 20554751 | 19590578 | 26867495 | 9872057 |
| 19159344 | 26149458 | 18375758 | 22846601 | 15863494 |
| 15803148 | 26489954 | 18724932 | 12069966 | 10490623 |

| | | | | |
|---|---|---|---|---|
| 9642287 | 18092812 | 1512203 | 11102867 | 29290584 |
| 11275267 | 12571226 | 10092508 | 8253761 | 24582806 |
| 11015622 | 21867484 | 10542045 | 14517336 | 27779865 |
| 15788404 | 12893295 | 12431977 | 10329735 | 15771582 |
| 869535 | 1902669 | 14663079 | 14734546 | 12415312 |
| 12360215 | 14506308 | 15522832 | 12802054 | 15156179 |
| 14630197 | 15033580 | 1649829 | 12807000 | 14985712 |
| 10586892 | 9535909 | 10938271 | 15623507 | 12360214 |
| 16470226 | 10751659 | 11823218 | 15635094 | 9927518 |
| 11079101 | 10873290 | 10473374 | 24364251 | 11905810 |
| 11449275 | 12566218 | 17636255 | 25509331 | 15738953 |
| 12145654 | 15094344 | 16707670 | 19837036 | 14670296 |
| 12488490 | 15955302 | 15470257 | 26563290 | 15629410 |
| 11708889 | 15033579 | 7639721 | 33112703 | 18086882 |
| 15466916 | 15033578 | 11341914 | 32156170 | 17571595 |
| 11244038 | 15639402 | 15520012 | 10978349 | 18430459 |
| 18322649 | 12154354 | 11042448 | 18065535 | 15902993 |
| 10376013 | 11253051 | 11274177 | 18819910 | 12589470 |
| 10577498 | 16700615 | 11099377 | 9596633 | 16935872 |
| 10224668 | 12762887 | 10473568 | 16407450 | 17395553 |
| 11228151 | 16095998 | 11551958 | 21252347 | 15459747 |
| 17050193 | 11057895 | 9506957 | 11513957 | 17041811 |
| 17308335 | 9643506 | 12512856 | 15350605 | 18951640 |
| 11304556 | 12084351 | 17253960 | 15613448 | 16288298 |
| 15485887 | 12675513 | 12716890 | 15982836 | 15336977 |
| 9545263 | 18482986 | 21138417 | 10707049 | 14526190 |
| 12821647 | 14583094 | 22899863 | 14596811 | 12372277 |
| 9417101 | 11157967 | 17928217 | 12943720 | 15242639 |
| 10601359 | 15610825 | 14568616 | 15080147 | 24739785 |
| 10755615 | 12049631 | 19661164 | 12963732 | 19525941 |
| 10586078 | 9677355 | 17239763 | 18708085 | 26920688 |
| 9581795 | 12750367 | 12847091 | 16412784 | 25191533 |
| 14756624 | 12967709 | 11390981 | 14735553 | 15695335 |
| 15179030 | 12121990 | 10318803 | 17900507 | 29427371 |
| 11573199 | 15537651 | 9756849 | 15882431 | 11807533 |
| 15459662 | 7813456 | 10742590 | 12359826 | 17081970 |
| 11229600 | 12751784 | 9151776 | 14556773 | 30941017 |
| 12235283 | 18395526 | 9988768 | 16147539 | 15996541 |
| 12640026 | 8892297 | 10520990 | 15372473 | 11738471 |
| 15688067 | 11150302 | 15304505 | 17784964 | 15996546 |
| 14519391 | 12949079 | 16099108 | 26875496 | 11839795 |
| 15108811 | 9851611 | 11030741 | 23719536 | 25270767 |
| 14744429 | 12446213 | 14706853 | 26077918 | 16522639 |
| 11082269 | 10713147 | 12511560 | 26064136 | 12844200 |
| 10698680 | 17922848 | 3161730 | 23799571 | 14711353 |
| 11882383 | 12466271 | 12716912 | 20708941 | 10615904 |
| 15053919 | 14982633 | 10837462 | 22534646 | 11371514 |
| 11146675 | 6798961 | 10748143 | 18583599 | 18056765 |
| 15034601 | 8662716 | 10854428 | 24925320 | 19136482 |
| 19675644 | 8995384 | 10854427 | 22056910 | 11225594 |
| 16632608 | 11294844 | 12655644 | 26366162 | 18572189 |

| | | | | |
|---|---|---|---|---|
| 14993306 | 15721837 | 15987739 | 15319346 | 16939974 |
| 18310071 | 17543969 | 18635948 | 15987776 | 16856873 |
| 14635525 | 15983048 | 34249946 | 9107551 | 15499580 |
| 11282018 | 15322192 | 34646033 | 4395686 | 10812966 |
| 12867054 | 18070909 | 11749383 | 15298956 | 10764727 |
| 10196204 | 15039466 | 12191611 | 16612326 | 10985348 |
| 9748166 | 10586030 | 12947395 | 18157157 | 13475371 |
| 11742412 | 15831826 | 12676795 | 11920679 | 16894175 |
| 15624019 | 11298294 | 11369511 | 11436301 | 11514508 |
| 16288289 | 12438359 | 12390245 | 27304501 | 16321959 |
| 10567225 | 9504340 | 14597384 | 24556838 | 5835946 |
| 9445476 | 18793215 | 12566928 | 25242279 | 21953451 |
| 9609110 | 10462516 | 12849693 | 26269525 | 14702404 |
| 9609111 | 15639739 | 20602996 | 26185979 | 17408485 |
| 15229476 | 16848789 | 22367796 | 26658722 | 11004450 |
| 9346240 | 11728310 | 23085193 | 24698685 | 15528644 |
| 17344645 | 10608501 | 22296764 | 26046644 | 11921398 |
| 18291668 | 14762510 | 19483709 | 26700098 | 18021072 |
| 11343120 | 14976004 | 23664135 | 24842496 | 19247843 |
| 12791379 | 15604886 | 15298336 | 23465396 | 22221911 |
| 12511876 | 14977592 | 15604203 | 23421405 | 26610024 |
| 19079156 | 15629883 | 15028942 | 23694989 | 26610025 |
| 15971106 | 15173832 | 14614204 | 19812304 | 23124073 |
| 18549797 | 15265790 | 15383650 | 16756488 | 16493424 |
| 18243016 | 10975856 | 10841026 | 27027448 | 24718315 |
| 17182536 | 12540948 | 15016298 | 28283069 | 24707857 |
| 18838301 | 14757320 | 11023702 | 15607802 | 25526305 |
| 19079167 | 12909723 | 12894244 | 16109839 | 19767725 |
| 17161619 | 12324235 | 16822996 | 15567854 | 20643110 |
| 18483500 | 11302930 | 12753399 | 12205470 | 26655628 |
| 22383755 | 9694817 | 9263329 | 11062493 | 27291964 |
| 22339650 | 1634517 | 10643996 | 15607806 | 25879280 |
| 21372217 | 12226220 | 10347209 | 11895890 | 26549800 |
| 25217450 | 8765137 | 11602344 | 12530970 | 23846113 |
| 21034406 | 12415297 | 11041354 | 16177799 | 22430786 |
| 25587654 | 11427694 | 16246839 | 8942070 | 24840700 |
| 24069565 | 9735948 | 16750612 | 16002670 | 23702978 |
| 16204702 | 9602501 | 15841168 | 9143705 | 25145756 |
| 23735217 | 14509570 | 16001050 | 14710949 | 23215645 |
| 24606160 | 15652868 | 16001072 | 11905813 | 25330206 |
| 10851053 | 10652548 | 16899407 | 11836514 | 25879288 |
| 11114747 | 11378299 | 15009714 | 12766758 | 27012466 |
| 8682149 | 15803158 | 11224709 | 12752668 | 26121197 |
| 30267440 | 14715912 | 15721476 | 15618404 | 26482951 |
| 34492272 | 15857686 | 14695152 | 9862693 | 22819539 |
| 32203403 | 9697848 | 15557758 | 11390470 | 21575908 |
| 11890722 | 11802770 | 10843728 | 11399424 | 29148036 |
| 10770277 | 1682209 | 12487150 | 16699811 | 25816776 |
| 7591091 | 12628346 | 10725110 | 14704852 | 16971555 |
| 10417174 | 1883197 | 10570027 | 11529497 | 17589524 |
| 7916951 | 2943218 | 10490505 | 10851172 | 16895466 |

| | | | | |
|---|---|---|---|---|
| 15578053 | 15987956 | 12459728 | 29477730 | 11741871 |
| 12215643 | 16878994 | 11407945 | 25588886 | 15697206 |
| 17506672 | 20145624 | 12946833 | 26104206 | 10572115 |
| 12379883 | 24733839 | 16702400 | 21488979 | 10913097 |
| 12741677 | 19652915 | 10740269 | 23927593 | 15901685 |
| 16870043 | 11557990 | 12648469 | 27606286 | 12700258 |
| 15948711 | 16269360 | 14586404 | 27617233 | 16788179 |
| 11902573 | 10944513 | 12545153 | 27732847 | 16428816 |
| 10944551 | 22875451 | 14580692 | 20300064 | 23279921 |
| 12867060 | 12488531 | 12079267 | 15318166 | 16458304 |
| 16112428 | 22735517 | 15774796 | 10647931 | 20023024 |
| 12407699 | 19941039 | 15823750 | 16557281 | 17055078 |
| 15942304 | 19306880 | 11707511 | 16551846 | 17055079 |
| 11895903 | 16760654 | 23340292 | 16946003 | 17010456 |
| 16189154 | 15182704 | 16495942 | 17062879 | 16631601 |
| 17625570 | 15136768 | 12938733 | 16242838 | 10537203 |
| 18593892 | 17480229 | 26501339 | 16189702 | 9603882 |
| 19629074 | 15182701 | 15767499 | 11165748 | 9098055 |
| 23201355 | 16518630 | 16972273 | 12094241 | 19016485 |
| 22154278 | 18631137 | 18047737 | 16339096 | 20660314 |
| 21855164 | 18598214 | 29090092 | 15122209 | 19580826 |
| 21753699 | 18550795 | 12814656 | 14634372 | 19439500 |
| 22736493 | 18946022 | 20495568 | 14685170 | 9461500 |
| 21850578 | 16554437 | 24781149 | 15611513 | 10217486 |
| 23733083 | 18096427 | 28468938 | 17211469 | 10518530 |
| 30941048 | 18311174 | 28838811 | 17287871 | 11007789 |
| 29128428 | 14691134 | 19663908 | 12351585 | 12196148 |
| 32326537 | 16469879 | 23352769 | 12878745 | 12195810 |
| 15288257 | 16584177 | 27103567 | 15724144 | 12055304 |
| 15300010 | 10580130 | 26297973 | 16551847 | 11283349 |
| 15288258 | 11821425 | 31100053 | 11900250 | 17506685 |
| 15288260 | 12626393 | 24548101 | 15082523 | 16245042 |
| 15585829 | 10350056 | 31587020 | 9422516 | 23042036 |
| 12464310 | 15466199 | 15189334 | 10835690 | 23922391 |
| 15272268 | 10207017 | 26487845 | 10453277 | 15828220 |
| 11896183 | 25727146 | 27546335 | 11750244 | 15828221 |
| 12417415 | 11439191 | 27581196 | 8069858 | 15828222 |
| 12015972 | 16557269 | 27829983 | 11527574 | 15828223 |
| 10028969 | 11747320 | 21674642 | 12711111 | 16476485 |
| 12681948 | 11313928 | 24522549 | 11720739 | 16601267 |
| 11436314 | 15116721 | 25761946 | 12616528 | 14999402 |
| 11436316 | 12505356 | 22926193 | 11807782 | 10529898 |
| 11436323 | 10714958 | 20736035 | 16064138 | 15673477 |
| 17276014 | 16697662 | 25611507 | 11463388 | 16511590 |
| 17884153 | 16915296 | 27911343 | 15109562 | 16360030 |
| 16922398 | 17409411 | 28823929 | 11559746 | 11921433 |
| 18018633 | 26037915 | 29147906 | 11905738 | 12543708 |
| 9111134 | 19584092 | 31054074 | 12243753 | 17346171 |
| 12200473 | 19240372 | 30370500 | 21193867 | 10198776 |
| 12684507 | 12135761 | 30937538 | 24508914 | 18081658 |
| 11278778 | 26646590 | 29720658 | 17251915 | 18488143 |

| | | | | |
|---|---|---|---|---|
| 11067870 | 12045215 | 12700767 | 11856310 | 9763449 |
| 16299351 | 12084906 | 14657486 | 12867427 | 10213783 |
| 9292932 | 11730319 | 11082272 | 16042598 | 9565594 |
| 16261175 | 11063930 | 10574699 | 10476961 | 9553086 |
| 11941303 | 10497333 | 15325584 | 10476960 | 10459012 |
| 17960151 | 12374568 | 16099218 | 10497122 | 12893879 |
| 17952897 | 12477927 | 16200202 | 10937989 | 10085287 |
| 18093537 | 11148128 | 15325588 | 10937990 | 10397773 |
| 14699405 | 21747944 | 15928708 | 11014182 | 11445562 |
| 11058677 | 20478527 | 15978794 | 11014183 | 12853481 |
| 15661024 | 19399032 | 11099404 | 15208624 | 9446565 |
| 14647478 | 25915839 | 16125113 | 18272355 | 10359592 |
| 27055367 | 25485532 | 19653858 | 19120475 | 9614185 |
| 25584786 | 18172298 | 19802558 | 19539500 | 11208059 |
| 20889378 | 28968951 | 20034776 | 18549796 | 10379359 |
| 27060771 | 16338359 | 21187343 | 19615405 | 11029060 |
| 32565065 | 18497808 | 21801009 | 19120476 | 10788491 |
| 18245809 | 16919475 | 24343578 | 18591413 | 8621431 |
| 15695097 | 16181324 | 25948417 | 19076341 | 9647643 |
| 17051207 | 19483713 | 18358634 | 18439848 | 9427746 |
| 11592845 | 23495936 | 25048860 | 18984593 | 9843576 |
| 17085779 | 28583440 | 11313942 | 19224920 | 12600315 |
| 14705136 | 29071527 | 19422821 | 32668226 | 10097106 |
| 10398674 | 22215586 | 17229979 | 19239894 | 9214619 |
| 15890824 | 10087917 | 15549172 | 16968219 | 9647644 |
| 27581199 | 10652088 | 15549171 | 16939780 | 9452464 |
| 31618844 | 10747959 | 15833268 | 17471261 | 12669082 |
| 31795242 | 11007775 | 15109615 | 17447862 | 10873817 |
| 30710214 | 18046402 | 16555051 | 16766188 | 10982406 |
| 23439666 | 17368028 | 12793984 | 17560162 | 9817754 |
| 28643372 | 16212506 | 12419208 | 10499798 | 10564279 |
| 29194372 | 12475201 | 16286563 | 10784442 | 9707576 |
| 18420416 | 10924740 | 16581295 | 19880312 | 11739776 |
| 16844381 | 17259602 | 12505055 | 18573085 | 10557242 |
| 21980554 | 19299134 | 12372844 | 31835031 | 11251103 |
| 25766616 | 15063851 | 12019279 | 21190829 | 9305917 |
| 17466621 | 19948674 | 15173164 | 19051051 | 9710615 |
| 10195903 | 16914536 | 17362841 | 20383543 | 9813082 |
| 14532116 | 9550618 | 25332897 | 17451884 | 10359608 |
| 31417367 | 16352913 | 24273531 | 11782468 | 11208143 |
| 29261664 | 17557941 | 21951628 | 23070004 | 9852078 |
| 21196165 | 19721811 | 18793332 | 16882042 | 10683148 |
| 23479001 | 12138151 | 23692861 | 10839363 | 9507000 |
| 28507517 | 11826292 | 24760442 | 11737951 | 9628864 |
| 15528202 | 11385088 | 16968212 | 11115874 | 10036234 |
| 15640354 | 11045400 | 17618621 | 15342965 | 9199167 |
| 20205843 | 18056784 | 11093772 | 10872468 | 11278762 |
| 29998397 | 11533299 | 17893666 | 11306253 | 11555414 |
| 28714865 | 10601204 | 16688755 | 11104523 | 9614193 |
| 15342004 | 10986234 | 16428379 | 11237004 | 11035026 |
| 15611724 | 11152757 | 12136096 | 11252968 | 11323436 |

| | | | | |
|---|---|---|---|---|
| 12068098 | 23959882 | 20134025 | 11171088 | 18431594 |
| 10397763 | 23467090 | 21283798 | 15122199 | 18566824 |
| 12011104 | 19284629 | 21781017 | 15928679 | 10073616 |
| 10652340 | 25820328 | 22896795 | 15084594 | 10070156 |
| 11305904 | 17287577 | 22928510 | 14656980 | 10099706 |
| 1495423 | 22570381 | 24548561 | 11905831 | 18655910 |
| 8457204 | 14715493 | 24651541 | 15286727 | 11773613 |
| 10366774 | 12471498 | 25152334 | 15343367 | 11001488 |
| 19239890 | 11082195 | 25469537 | 15099566 | 15522987 |
| 10978320 | 12358600 | 25483301 | 12669022 | 17956998 |
| 17537823 | 15073291 | 25757376 | 15489916 | 14586738 |
| 19733268 | 15340795 | 22892070 | 12133805 | 13678960 |
| 16531502 | 15184572 | 12042308 | 14647476 | 11741095 |
| 12777052 | 15883781 | 18094055 | 15308211 | 15602010 |
| 10998348 | 4297098 | 23885123 | 15345216 | 16104843 |
| 11469811 | 16262699 | 17275323 | 14617194 | 14641020 |
| 12604236 | 9858571 | 17457343 | 19132916 | 15338053 |
| 12832414 | 24375100 | 16932750 | 12195013 | 14724572 |
| 15642792 | 14534577 | 17395581 | 19290929 | 12391145 |
| 16413106 | 11252892 | 16497588 | 17395537 | 10942706 |
| 16547389 | 14557817 | 14698224 | 16799472 | 11032799 |
| 17459698 | 14769828 | 12734364 | 17187070 | 11292584 |
| 17475203 | 11252746 | 14751764 | 16922370 | 12102686 |
| 17926129 | 12163231 | 17637696 | 16922408 | 12403808 |
| 22982583 | 11836504 | 17507094 | 16922367 | 13679513 |
| 22633058 | 12466190 | 11101877 | 16816840 | 11953326 |
| 20143161 | 10021351 | 16924467 | 16405856 | 11207565 |
| 22092713 | 9847239 | 15749753 | 12944364 | 11349009 |
| 21833341 | 15063168 | 15823385 | 17588513 | 12671039 |
| 16452431 | 10519551 | 11078440 | 15292141 | 14659695 |
| 9695921 | 19758997 | 11272200 | 17458653 | 15204437 |
| 18929068 | 8093006 | 15759102 | 12689814 | 17156122 |
| 18305268 | 10934147 | 12480546 | 16339747 | 17464284 |
| 12835292 | 16642010 | 11390407 | 12693609 | 11454785 |
| 17943287 | 26735394 | 9011569 | 16870827 | 18039131 |
| 8048228 | 26160612 | 7491105 | 19120701 | 12920582 |
| 16932712 | 15975920 | 15258147 | 11165670 | 17200452 |
| 15459673 | 15671031 | 9974390 | 17158339 | 12439638 |
| 15972354 | 33925597 | 11325516 | 17200683 | 16845885 |
| 10430616 | 20495575 | 15585596 | 15269340 | 16413486 |
| 12965173 | 27353478 | 9719467 | 11773611 | 10904843 |
| 10025913 | 22169974 | 11507694 | 18040024 | 18357777 |
| 15380523 | 17525755 | 8072542 | 15026027 | 18545064 |
| 24607224 | 15820555 | 8786033 | 10221985 | 19357408 |
| 24385952 | 16650994 | 12796471 | 10911373 | 10051305 |
| 23261954 | 16771626 | 15889095 | 18552454 | 14635791 |
| 21481196 | 17182900 | 15889096 | 18703404 | 14745971 |
| 23587289 | 17950242 | 14617043 | 11709400 | 12446114 |
| 20696704 | 18005931 | 8550403 | 15725705 | 14530294 |
| 19439032 | 18946474 | 1194238 | 9458738 | 15102436 |
| 17108952 | 20094051 | 18435614 | 12060106 | 12967557 |

| | | | | |
|---|---|---|---|---|
| 11433294 | 10068641 | 18458083 | 7927516 | 18426796 |
| 23256519 | 17368312 | 18287005 | 1702903 | 14742429 |
| 19482104 | 14634621 | 15485873 | 16181335 | 16687408 |
| 18838381 | 9390557 | 18772128 | 20171863 | 14630918 |
| 23151663 | 14679297 | 18094042 | 17157489 | 20022965 |
| 28522145 | 15502848 | 8327893 | 19218463 | 16931790 |
| 25808920 | 15210742 | 7641683 | 11742988 | 18840095 |
| 32888647 | 19817485 | 8997178 | 16413484 | 18048764 |
| 28242765 | 17562450 | 14514350 | 12049731 | 25821458 |
| 18832364 | 19941817 | 10449730 | 16982218 | 18038217 |
| 16291214 | 21642957 | 17900700 | 11163244 | 34611326 |
| 20103563 | 19276888 | 10903473 | 12209154 | 16391004 |
| 17709369 | 20933024 | 16098514 | 4561027 | 12704645 |
| 21039332 | 21543634 | 29053970 | 15218528 | 10205060 |
| 11048892 | 21757760 | 25561175 | 15723341 | 12440953 |
| 21352852 | 17283063 | 30844724 | 9422506 | 22094468 |
| 2825027 | 10934204 | 26378229 | 17599937 | 21347367 |
| 9450929 | 15657177 | 25567907 | 16107722 | 15231740 |
| 18691557 | 10777492 | 28066558 | 17961083 | 20603001 |
| 19481063 | 22986504 | 27648300 | 15924268 | 16756494 |
| 19126755 | 12192414 | 29236692 | 19448080 | 18160255 |
| 12161432 | 15691767 | 17470430 | 15901243 | 11809809 |
| 10879540 | 16459297 | 28686218 | 30056832 | 15592430 |
| 18237417 | 22981989 | 24529380 | 15920196 | 17698587 |
| 18043734 | 10966113 | 11724780 | 27558536 | 26393581 |
| 20418862 | 14614827 | 21427764 | 23686171 | 11034610 |
| 18296446 | 23079593 | 5661997 | 17514195 | 20708099 |
| 15965470 | 19906846 | 24469809 | 11160393 | 26545300 |
| 9819428 | 16339192 | 23348840 | 12008020 | 28630109 |
| 18077452 | 20395968 | 23530209 | 1350383 | 11429545 |
| 15652348 | 21356043 | 23351793 | 9495808 | 23931059 |
| 19567869 | 15564374 | 21816947 | 26045554 | 26339618 |
| 19524514 | 9069263 | 25047614 | 17997397 | 26772212 |
| 23064382 | 592399 | 13978319 | 7903167 | 22030238 |
| 24465591 | 641056 | 8918697 | 28303017 | 23000944 |
| 19671665 | 9732867 | 10648173 | 8139656 | 25060581 |
| 17991742 | 11779870 | 18187424 | 11823786 | 16603771 |
| 11751416 | 11323433 | 8306975 | 24003239 | 23667505 |
| 10486198 | 11744733 | 1081164 | 23795651 | 19074509 |
| 18066052 | 1998122 | 8163536 | 26139588 | 20859064 |
| 18816852 | 10497269 | 1376928 | 17716973 | 19149539 |
| 25560970 | 20525283 | 9468499 | 25689043 | 17572487 |
| 18412118 | 16698963 | 16849466 | 23187890 | 10519914 |
| 18394579 | 19416660 | 19065780 | 16762929 | 9646295 |
| 21497760 | 19075029 | 18596693 | 33298899 | 8387282 |
| 10662633 | 15152008 | 11606584 | 16549788 | 14706825 |
| 10668974 | 19741254 | 15489917 | 22751019 | 3317417 |
| 24529385 | 19737925 | 7929028 | 17230199 | 12941277 |
| 20435029 | 9487141 | 18818202 | 19379695 | 22013193 |
| 23973942 | 20705608 | 6335036 | 12724779 | 21878990 |
| 19575673 | 18195019 | 1550550 | 16495441 | 20533886 |

| | | | | |
|---|---|---|---|---|
| 12819131 | 15837415 | 17221863 | 17533373 | 22547309 |
| 12941276 | 23746838 | 15146185 | 11238451 | 21312325 |
| 23482940 | 9282108 | 21924235 | 18228324 | 21478859 |
| 16382132 | 18555778 | 38372588 | 7629134 | 20858899 |
| 22326026 | 13905658 | 24191001 | 1396589 | 21799911 |
| 9362062 | 19597488 | 16760263 | 10454565 | 22363759 |
| 20399632 | 19818705 | 11136719 | 11907280 | 16446366 |
| 22389062 | 22048312 | 18977199 | 19362144 | 14684739 |
| 15998802 | 14615387 | 19608861 | 30354722 | 11970895 |
| 17574030 | 11282294 | 17356064 | 18839291 | 22826439 |
| 20145001 | 8578462 | 20703077 | 24179613 | 19494126 |
| 15314642 | 15118080 | 17525332 | 20522543 | 22986494 |
| 23567335 | 15664996 | 12086618 | 22382365 | 8034666 |
| 12169685 | 22653444 | 20622854 | 23706667 | 2344612 |
| 11384985 | 23945166 | 21498573 | 21533068 | 23212477 |
| 12620239 | 25040720 | 18367646 | 9878398 | 2188731 |
| 19851329 | 17955020 | 21478295 | 16169847 | 21413788 |
| 12620240 | 18786386 | 18079394 | 14744247 | 2188730 |
| 12011067 | 18775307 | 19962665 | 17254765 | 22134922 |
| 12598906 | 16746585 | 20009532 | 15557120 | 19410545 |
| 17254298 | 34884868 | 18317450 | 8861902 | 19660450 |
| 15788703 | 33321098 | 16339315 | 15494733 | 20080577 |
| 20816209 | 15987702 | 17363375 | 17395582 | 17339380 |
| 12670394 | 27049945 | 23637185 | 16214811 | 19410539 |
| 10849438 | 30844107 | 16702223 | 18989317 | 19629040 |
| 30621321 | 6773467 | 23873287 | 12408825 | 22196163 |
| 28350208 | 18926908 | 10469566 | 15525529 | 21453242 |
| 31113848 | 17145306 | 24119662 | 18632669 | 21473702 |
| 28029388 | 8717519 | 25364732 | 10898789 | 19061483 |
| 28571533 | 12495737 | 23732476 | 16510874 | 22286106 |
| 29858187 | 16280322 | 19854180 | 18573912 | 20064600 |
| 23406282 | 16717127 | 23378591 | 11080158 | 23268465 |
| 9566895 | 20554521 | 8001155 | 12408824 | 21375475 |
| 10712901 | 10430017 | 22130221 | 14749774 | 16325574 |
| 10339564 | 25062251 | 10562278 | 16141059 | 22201797 |
| 11046155 | 15007061 | 23783758 | 15475613 | 12944365 |
| 10995389 | 28146471 | 10825291 | 18608124 | 19243136 |
| 10801458 | 26872272 | 9288971 | 17052452 | 20418328 |
| 7641697 | 9736607 | 11044372 | 19521399 | 20074029 |
| 15304631 | 15339664 | 25303525 | 23606632 | 15571815 |
| 19339544 | 26042225 | 16799259 | 2738071 | 18234723 |
| 19704417 | 8953040 | 21438114 | 2775660 | 19619488 |
| 20728359 | 19092055 | 21711675 | 2713493 | 15459103 |
| 20965423 | 1177317 | 27245231 | 2592373 | 10647780 |
| 29078414 | 27092250 | 22664329 | 3009023 | 19759537 |
| 24732012 | 21445324 | 21679683 | 22183981 | 10581160 |
| 23216249 | 26829388 | 27974214 | 16151858 | 17318175 |
| NA | 12719478 | 10374692 | 15687324 | 10421629 |
| 28797581 | 17113138 | 12876557 | 21907012 | 11336703 |
| 20188139 | 16604071 | 16598076 | 16846473 | 25840006 |
| 28546457 | 19696148 | 24705298 | 9587028 | 23730680 |

| | | | | |
|---|---|---|---|---|
| 12756558 | 1976634 | 11336673 | 12511868 | 23561633 |
| 20414203 | 12239347 | 15469835 | 21851428 | 23038248 |
| 31374202 | 21376233 | 15848799 | 26048987 | 11080164 |
| 22237395 | 25081058 | 15093606 | 23325218 | 15621527 |
| 26708048 | 17210635 | 17300218 | 18082604 | 16901784 |
| 31048490 | 10535941 | 20937854 | 25099582 | 17158953 |
| 25493225 | 11427533 | 20026667 | 17768402 | 8876179 |
| 28572265 | 11389444 | 17707230 | 21734703 | 22902626 |
| 17887919 | 17591695 | 23150253 | 16289108 | 25435140 |
| 26746812 | 19135894 | 23153826 | 12496284 | 22466610 |
| 27932076 | 21095583 | 23063748 | 12565875 | 32152397 |
| 31116084 | 16751101 | 18631128 | 12629551 | 17290000 |
| 29117491 | 22045334 | 15491402 | 19758793 | 12660172 |
| 22983739 | 11278251 | 10747205 | 18606873 | 20951344 |
| 21384229 | 11158580 | 8978277 | 17101212 | 22617422 |
| 2301971 | 14718519 | 25887776 | 11461910 | 11713476 |
| 22838182 | 20937913 | 8677749 | 17646408 | 11136974 |
| 22165913 | 10757800 | 11443118 | 15546612 | 10831834 |
| 21388382 | 20061380 | 19616933 | 17909001 | 11940650 |
| 14742321 | 16027725 | 20347046 | 17122878 | 1361170 |
| 10747039 | 22344298 | 15958495 | 19909365 | 15458387 |
| 15964013 | 9739761 | 22101327 | 16597617 | 12048180 |
| 22055184 | 11904382 | 15837422 | 12791257 | 16885393 |
| 12486230 | 24402227 | 18614053 | 12141425 | 24452601 |
| 8434121 | 16246722 | 18377699 | 9858549 | 25843800 |
| 15790808 | 19023283 | 20148673 | 10982855 | 18787071 |
| 9357549 | 23912815 | 28539435 | 15863030 | 18444242 |
| 15574587 | 27312108 | 25303530 | 17505008 | 16493026 |
| 8242752 | 24699078 | 25597633 | 18757403 | 21159781 |
| 9843217 | 19754430 | 17510434 | 11325814 | 15084259 |
| 10669743 | 26085183 | 23850489 | 9154000 | 11964479 |
| 2342578 | 23732108 | 10873660 | 20101236 | 11094066 |
| 19528227 | 23657496 | 19274700 | 19103595 | 23580232 |
| 8242751 | 19489725 | 23954741 | 11003564 | 21196578 |
| 16166375 | 19436320 | 19339991 | 9538690 | 23918930 |
| 2392154 | 23624913 | 27288742 | 18381441 | 14966295 |
| 17694070 | 21307119 | 17037979 | 11056689 | 24047697 |
| 22781841 | 11283727 | 21680538 | 17552943 | 26780985 |
| 7079182 | 15021893 | 17567461 | 9266968 | 21164521 |
| 24214338 | 26864683 | 10441588 | 21080029 | 17667983 |
| 23687512 | 26423937 | 3893466 | 18406375 | 26715362 |
| 24589714 | 10727776 | 18110463 | 20032467 | 11777915 |
| 16403636 | 26054742 | 2729894 | 11012776 | 28890329 |
| 23938249 | 15001665 | 224930 | 19799112 | 16943287 |
| 21243710 | 7938166 | 21917992 | 18698078 | 23707940 |
| 23587238 | 17450176 | 15496458 | 19276253 | 10102273 |
| 19657014 | 19345195 | 15956359 | 17475912 | 22412893 |
| 23166392 | 17613433 | 14534365 | 7692230 | 20978166 |
| 24059801 | 12482742 | 17916740 | 30552988 | 14576824 |
| 1536007 | 11208594 | 29980625 | 17668209 | 19276113 |
| 7623835 | 10551886 | 21354435 | 8104191 | 29533771 |

| | | | | |
|---|---|---|---|---|
| 25425107 | 9662327 | 16533948 | 24141787 | 10684278 |
| 27730540 | 8050359 | 9458043 | 17996710 | 9188632 |
| 15824087 | 9020173 | 16102042 | 11376695 | 7474100 |
| 15866051 | 14990995 | 7997267 | 11242102 | 7474080 |
| 14701673 | 10228158 | 12005431 | 18285462 | 18839070 |
| 26700816 | 20655481 | 15601831 | 14976220 | 20156840 |
| 27869526 | 17364682 | 17488973 | 10930435 | 19356150 |
| 22511764 | 15269338 | 8548048 | 19442142 | 177353 |
| 27547294 | 20227374 | 17041586 | 22216413 | 11008000 |
| 22876196 | 26131740 | 25789972 | 9839614 | 8605876 |
| 16840535 | 8665925 | 16012167 | 15944408 | 17969444 |
| 11779849 | 8522583 | 25201414 | 9716487 | 8621447 |
| 29056338 | 11737941 | 25643323 | 14512302 | 15607020 |
| 26523980 | 11146276 | 25642960 | 12133942 | 19059436 |
| 24291305 | 9861669 | 19357644 | 9885206 | 18929502 |
| 25007762 | 19412884 | 20483915 | 21349098 | 14525967 |
| 10913147 | 17560368 | 25642963 | 23436998 | 19022706 |
| 21389279 | 19165215 | 18539610 | 20890289 | 18613828 |
| 26052839 | 21755468 | 9837724 | 26581522 | 19302047 |
| 12239572 | 19239886 | 17962301 | 14527418 | 22595692 |
| 24167160 | 25565029 | 12139939 | 19457864 | 20410258 |
| 24920680 | 15741316 | 12791985 | 20144995 | 7632928 |
| 10358014 | 21801010 | 17928296 | 18624796 | 16775004 |
| 20921137 | 14723851 | 17965729 | 22252131 | 16061793 |
| 18612045 | 10075717 | 21804533 | 15075377 | 16049494 |
| 24040102 | 17442700 | 15485900 | 11836526 | 16229832 |
| 27731402 | 2105456 | 10212258 | 19900460 | 11756463 |
| 24028821 | 17000779 | 21052091 | 16113677 | 25343031 |
| 21536589 | 15087129 | 22977173 | 11081636 | 24068461 |
| 16783365 | 9452474 | 16396909 | 19432801 | 21421921 |
| 27791111 | 17306374 | 8824192 | 20359876 | 16157178 |
| 17395749 | 11815463 | 18585355 | 17943301 | 19575639 |
| 18827341 | 9914469 | 21257310 | 11331907 | 25233431 |
| 11447289 | 12475787 | 22902404 | 20414202 | 21436057 |
| 16682955 | 17478680 | 11395778 | 27193484 | 23839779 |
| 23493540 | 16990140 | 15163793 | 20646998 | 23846822 |
| 23493538 | 8825650 | 23565096 | 25535373 | 25708460 |
| 26199742 | 9701566 | 15075292 | 25713138 | 23881164 |
| 24666553 | 21220505 | 9593755 | 20676083 | 15069402 |
| 13093749 | 21382416 | 21502959 | 23142665 | 24316379 |
| 18220710 | 20823909 | 16987893 | 28992441 | 16413926 |
| 8213607 | 21704024 | 26741528 | 16777091 | 19846511 |
| 25639270 | 19703651 | 29452096 | 25043001 | 20213925 |
| 5823111 | 20919643 | 9759480 | 10508650 | 16287098 |
| 9778043 | 29747811 | 10550206 | 17084981 | 18626067 |
| 8288595 | 11408587 | 20871616 | 19118207 | 19851335 |
| 10597240 | 10679020 | 19268590 | 28106780 | 9211984 |
| 15014447 | 18385516 | 19369211 | 14754902 | 17189949 |
| 15084262 | 11739401 | 23149936 | 17624957 | 20519495 |
| 7929342 | 20937827 | 9880493 | 8041786 | 16630821 |
| 11988738 | 11285137 | 20965415 | 10620603 | 8906794 |

| | | | | |
|---|---|---|---|---|
| 21664576 | 21599891 | 19703008 | 19962410 | 12526811 |
| 16136078 | 25436572 | 15364901 | 24842111 | 11208076 |
| 8906787 | 21647776 | 15211354 | 23149385 | 9384576 |
| 8898207 | 21647775 | 17057362 | 14715917 | 9811606 |
| 22326621 | 10755665 | 8252621 | 15738989 | 9564030 |
| 17641202 | 22791193 | 19330724 | 15886331 | 19933846 |
| 8609415 | 23620664 | 23511973 | 14603310 | 21613978 |
| 17168669 | 17215883 | 24388755 | 34758268 | 21177493 |
| 19339967 | 15157936 | 24071720 | 15111103 | 7478553 |
| 28642194 | 12827356 | 28610953 | 17244524 | 8626428 |
| 23376167 | 16675045 | 23732472 | 9724731 | 19219045 |
| 21921683 | 16908340 | 16212495 | 1828394 | 12072434 |
| 20485518 | 14530973 | 29364559 | 9508775 | 19115315 |
| 22586411 | 17374385 | 31894645 | 9448290 | 24835590 |
| 25732612 | 19164095 | 17035353 | 8001816 | 23382213 |
| 23770565 | 17439925 | 29222186 | 8078588 | 20805990 |
| 12126930 | 16825786 | 15102545 | 1828392 | 25114211 |
| 9893266 | 12836025 | 16126164 | 8246996 | 17496916 |
| 11324705 | 16720382 | 19262508 | 23842645 | 23443682 |
| 21779496 | 19946895 | 15744310 | 10773440 | 15664191 |
| 22569528 | 10871282 | 20649548 | 9190208 | 9001234 |
| 19091303 | 24518591 | 22817841 | 8497257 | 23208509 |
| 22733995 | 16849525 | 21629705 | 7859739 | 14684163 |
| 22800433 | 12006644 | 22146081 | 1828393 | 12475979 |
| 18593949 | 8703035 | 19575678 | 12360190 | 24828152 |
| 21329875 | 15314156 | 22265687 | 20584987 | 9442023 |
| 21372320 | 9038378 | 19682549 | 22500797 | 9009191 |
| 21164013 | 24951116 | 19076454 | 21552327 | 8625412 |
| 16600872 | 8906793 | 20533901 | 15930137 | 12807427 |
| 21164014 | 18779368 | 19008463 | 18760361 | 14672936 |
| 34782609 | 9070211 | 18430751 | 15662836 | 25740432 |
| 34235568 | 12147692 | 24366263 | 15242646 | 22001063 |
| 34495539 | 11894096 | 20079335 | 17301054 | 11585921 |
| 18952054 | 10521801 | 25463276 | 18206966 | 21402876 |
| 26733807 | 8384622 | 19021507 | 17098746 | 20006983 |
| 24150225 | 11478803 | 20522547 | 23290524 | 21031079 |
| 18511691 | 24814316 | 20873749 | 18782863 | 10383463 |
| 23493374 | 16782899 | 17996737 | 16980297 | 18322270 |
| 19628041 | 15713485 | 19767391 | 25314029 | 12775774 |
| 20385708 | 16537482 | 19720617 | 10597293 | 11357136 |
| 22262897 | 24823486 | 28765093 | 12740388 | 15908424 |
| 23259945 | 23626807 | 20214493 | 16735510 | 11448771 |
| 18989361 | 15301554 | 28301740 | 19245792 | 12832477 |
| 26214522 | 7513258 | 20100523 | 10713156 | 16571882 |
| 25424712 | 20015050 | 24440477 | 20029837 | 15994232 |
| 22231481 | 8208547 | 24362565 | 9790970 | 19649571 |
| 25579386 | 20670691 | 24911204 | 23394999 | 23650620 |
| 15951480 | 11420688 | 22986124 | 23001041 | 1378856 |
| 18974038 | 21664574 | 25088301 | 22836579 | 20338884 |
| 1390320 | 8689559 | 19716793 | 11593029 | 16585540 |
| 15583024 | 9316394 | 22172991 | 12414957 | 16618810 |

| | | | | |
|---|---|---|---|---|
| 12884866 | 23362348 | 10744726 | 24814062 | 17956313 |
| 18585455 | 11866432 | 11679632 | 24202393 | 9090381 |
| 18414735 | 11986231 | 16061658 | 22589270 | 27977397 |
| 19483673 | 21898486 | 15023535 | 24651010 | 23798008 |
| 22179609 | 18724371 | 23230144 | 22020331 | 25177298 |
| 22373579 | 23362349 | 14599765 | 19451217 | 24508507 |
| 10802669 | 10958687 | 10583946 | 17254968 | 10767286 |
| 23306437 | 24842903 | 11900249 | 11564866 | 21464226 |
| 21293379 | 19640841 | 19008000 | 16337592 | 29321502 |
| 11955432 | 15533827 | 1370859 | 15287722 | 3693402 |
| 23505375 | 25125655 | 18938244 | 17210787 | 27284042 |
| 23727112 | 25356737 | 17509148 | 12189133 | 22555564 |
| 10572166 | 20804727 | 27587538 | 7816143 | 20385596 |
| 22529269 | 18593473 | 18043707 | 9628874 | 15581572 |
| 8422676 | 16368887 | 2105931 | 9162092 | 12948489 |
| 12023295 | 12386158 | 19561302 | 22325352 | 16378245 |
| 10757784 | 18974129 | 9590179 | 12134156 | 1656221 |
| 19001859 | 11971902 | 24098485 | 7588633 | 10579907 |
| 23333306 | 16365048 | 24388663 | 8846784 | 19644473 |
| 11030616 | 27749937 | 33447881 | 9030721 | 21430697 |
| 10959836 | 27841855 | 11060288 | 8774846 | 10866302 |
| 17470781 | 10837024 | 25977809 | 12171911 | 9326929 |
| 25512557 | 11518718 | 20200447 | 9195981 | 17611497 |
| 15451442 | 24667410 | 16720576 | 8622688 | 18954143 |
| 11493912 | 10964583 | 16469703 | 9564042 | 17604717 |
| 19859661 | 23576639 | 17973576 | 8622669 | 19487573 |
| 17531985 | 18239137 | 1840504 | 15866172 | 9467011 |
| 21496649 | 17948123 | 7522482 | 11062067 | 10555148 |
| 20795950 | 15531924 | 14983052 | 19462008 | 16949365 |
| 17352659 | 12374742 | 16402120 | 15563468 | 17940011 |
| 17671086 | 12897056 | 12839923 | 17395589 | 22221002 |
| 12446846 | 14986688 | 26105537 | 10842357 | 21427766 |
| 12842859 | 16530045 | 10733595 | 11067851 | 23980886 |
| 16530044 | 16921404 | 9824166 | 12096136 | 12558974 |
| 15107403 | 16429119 | 11883935 | 22722607 | 10918587 |
| 8687460 | 11112321 | 10722742 | 16764946 | 12089343 |
| 12391150 | 10487760 | 14729966 | 19096775 | 19196647 |
| 8649374 | 16024779 | 9529249 | 17513417 | 20861159 |
| 18374649 | 11535832 | 8552081 | 19096781 | 20192774 |
| 8001118 | 12370315 | 15572696 | 9780000 | 20043944 |
| 18094723 | 19520913 | 27812880 | 12200445 | 20559318 |
| 12050117 | 20368433 | 9704925 | 9214649 | 17605038 |
| 10713164 | 24060861 | 22336108 | 10559261 | 16828287 |
| 16256737 | 18097461 | 20583212 | 19365405 | 18216768 |
| 15569487 | 29455642 | 11234019 | 17433286 | 18216767 |
| 20156974 | 27573755 | 7588608 | 19204114 | 18726176 |
| 10206645 | 22901806 | 18365017 | 21983142 | 17330069 |
| 10551863 | 21335236 | 8259215 | 9598355 | 15269334 |
| 21632553 | 12589057 | 8521522 | 20004982 | 16125054 |
| 18469519 | 18316408 | 25907612 | 23140633 | 2018971 |
| 10934472 | 16912174 | 21993244 | 18848778 | 8704248 |

| | | | | |
|---|---|---|---|---|
| 12411949 | 11896061 | 18487508 | 29723737 | 16503871 |
| 1672265 | 21102611 | 17681183 | 25818528 | 19416966 |
| 12681240 | 17591783 | 19473982 | 31277234 | 10217147 |
| 16543958 | 14563852 | 21289308 | 26036474 | 10358075 |
| 9295288 | 17768054 | 19884659 | 32178475 | 15987244 |
| 15561706 | 12192048 | 26415504 | 23850828 | 11237865 |
| 11554294 | 11031238 | 16702404 | 32542921 | 24105466 |
| 1670772 | 7597053 | 18172008 | 11447113 | 17994015 |
| 16574391 | 23468639 | 25737250 | 11821947 | 26239616 |
| 18616426 | 22231403 | 24344134 | 24937142 | 20673872 |
| 16153702 | 25659039 | 26279303 | 2993863 | 19331822 |
| 17183653 | 11826263 | 16829519 | 23993095 | 22357867 |
| 16846370 | 17696772 | 17974977 | 11579234 | 18566290 |
| 20505756 | 19453276 | 26561776 | 15710605 | 13678582 |
| 16819296 | 26172624 | 25728608 | 23954936 | 22365832 |
| 15110706 | 25833379 | 15471878 | 21779505 | 18615013 |
| 24096405 | 25815272 | 10913193 | 18623629 | 12207013 |
| 23473600 | 26187467 | 17483331 | 24671035 | 23856247 |
| 21941108 | 22708562 | 15572663 | 18454133 | 20223231 |
| 19738629 | 26235645 | 22084245 | 8900285 | 22374676 |
| 21248841 | 24652853 | 15539407 | 10811826 | 21532586 |
| 25232742 | 25175772 | 20606012 | 10804209 | 22778133 |
| 15753315 | 21883762 | 20554524 | 24452471 | 17589498 |
| 18474358 | 24457024 | 11114724 | 11575290 | 26580584 |
| 18426888 | 19775879 | 23814047 | 12591178 | 8700862 |
| 20351093 | 23800009 | 22231447 | 18094957 | 16166372 |
| 21490058 | 19940261 | 29362479 | 15457548 | 15331439 |
| 10669757 | 23061719 | 28462526 | 19706539 | 22759635 |
| 9727485 | 24367090 | 33033617 | 23217742 | 26749280 |
| 20805998 | 25547115 | 27246931 | 18547144 | 21343615 |
| 21602795 | 16418168 | 14574414 | 18701067 | 19114653 |
| 19723635 | 20080666 | 23303910 | 21729787 | 25645944 |
| 20080834 | 11781575 | 7991565 | 27819291 | 22412390 |
| 10935637 | 26280536 | 21330364 | 24012004 | 19858498 |
| 20495621 | 11378386 | 25402849 | 20071533 | 11203699 |
| 21551231 | 11545734 | 20466531 | 19258013 | 8929538 |
| 19714243 | 18757404 | 22251903 | 7539918 | 20139099 |
| 19773441 | 19345329 | 22694141 | 9000051 | 19235904 |
| 1764995 | 26910647 | 23898197 | 10202154 | 25452107 |
| 9187150 | 18327259 | 16855591 | 19788888 | 25562167 |
| 8681805 | 26183061 | 20937701 | 9840812 | 20206228 |
| 19701457 | 18199536 | 8164745 | 9632793 | 26716895 |
| 20170518 | 26862215 | 27790713 | 18187047 | 16751105 |
| 8293978 | 19440552 | 28202288 | 14668441 | 21931630 |
| 7918097 | 12297295 | 32460002 | 11959977 | 26778333 |
| 9653148 | 25867063 | 28055293 | 10526574 | 23403054 |
| 20674646 | 23013792 | 32362820 | 16710296 | 15703179 |
| 16738315 | 11035045 | 29330095 | 11751371 | 25260729 |
| 16253240 | 25884169 | 25239922 | 12466191 | 18541707 |
| 16738331 | 22427670 | 25309333 | 17047378 | 22422618 |
| 20462400 | 23073177 | 31027363 | 18288277 | 12000792 |

| | | | | |
|---|---|---|---|---|
| 16901655 | 21317932 | 26526991 | 18086875 | 9535855 |
| 11857736 | 14702041 | 16246168 | 24607366 | 11278486 |
| 16728642 | 12914926 | 27313038 | 24388711 | 16204230 |
| 10215629 | 10435622 | 7711290 | 25354898 | 10470034 |
| 10911365 | 19293287 | 15277680 | 31956375 | 22340718 |
| 10572046 | 21814038 | 20435355 | 27707839 | 21098037 |
| 18767071 | 16213212 | 18632300 | 33195688 | 19513348 |
| 18850007 | 17499002 | 21377628 | 33767988 | 18006505 |
| 26004068 | 11923872 | 23765164 | 20405038 | 10924361 |
| 23333304 | 19250907 | 21183348 | 23716662 | 9988689 |
| 27048832 | 21986495 | 9590177 | 23444367 | 19723632 |
| 23431171 | 8319905 | 19109890 | 22293177 | 10753910 |
| 16839880 | 12654245 | 17036197 | 9792683 | 18429822 |
| 22682249 | 7878469 | 27314390 | 16202622 | 12221077 |
| 20228809 | 17121812 | 32354068 | 16431929 | 17546647 |
| 11099047 | 15103385 | 29768694 | 14595335 | 10940294 |
| 20660729 | 8463288 | 31871319 | 8543060 | 8662891 |
| 17189187 | 1629222 | 19490898 | 15507212 | 22228765 |
| 22864287 | 9108399 | 22539766 | 9497258 | 15480428 |
| 18719709 | 15121853 | 27875099 | 21868368 | 12473120 |
| 20691894 | 20925119 | 27676292 | 16472745 | 10873388 |
| 25356872 | 11266540 | 11905808 | 17198702 | 15504364 |
| 17145718 | 8955148 | 29385717 | 15085137 | 8702756 |
| 18278067 | 17395641 | 24622787 | 12906795 | 8662509 |
| 20673990 | 11440997 | 12636908 | 21576392 | 8816443 |
| 15186775 | 7608159 | 12577064 | 24914801 | 17151359 |
| 17719541 | 19329425 | 28098758 | 10678165 | 8617235 |
| 11684014 | 11601988 | 32377503 | 16943183 | 11018042 |
| 22362889 | 15871698 | 9614181 | 20101212 | 20803696 |
| 15525938 | 18957201 | 18075254 | 21397845 | 26363959 |
| 11511361 | 16648486 | 27740624 | 15944396 | 28664531 |
| 21625555 | 19305001 | 28196644 | 11779461 | 27548350 |
| 16415881 | 16855289 | 11809807 | 16292343 | 22103495 |
| 20870725 | 20417600 | 18311041 | 10958683 | 31208035 |
| 22611192 | 22593209 | 19846717 | 17244649 | 16190977 |
| 17591690 | 21987589 | 19269969 | 11948177 | 16254241 |
| 17349958 | 15339662 | 10967094 | 8798539 | 22931484 |
| 15053879 | 9649503 | 27556037 | 9199170 | 15546864 |
| 15866171 | 15737063 | 11306516 | 21730051 | 29596304 |
| 10710310 | 14730319 | 24613967 | 8756646 | 31514269 |
| 10673501 | 17112726 | 25482645 | 23875749 | 22260703 |
| 10570149 | 21878504 | 12456725 | 19628035 | 23648569 |
| 12064478 | 2862839 | 31303019 | 16762923 | 27314616 |
| 20959462 | 29101239 | 22619228 | 1660188 | 31344837 |
| 11447225 | 11560503 | 24256248 | 16636067 | 17222083 |
| 10673500 | 18450746 | 31454517 | 16507994 | 12778124 |
| 10884347 | 23354059 | 30513825 | 11896053 | 10072071 |
| 12393879 | 28814160 | 26555387 | 11145705 | 24128008 |
| 11239457 | 11106749 | 17504809 | 17237347 | 12732141 |
| 11546806 | 27629041 | 29189096 | 7559638 | 24969695 |
| 12851404 | 15774759 | 15556869 | 10975528 | 18237546 |

| | | | | |
|---|---|---|---|---|
| 12110184 | 25742728 | 11276226 | 30181355 | 21711248 |
| 16979770 | 11526392 | 11604131 | 11509732 | 21330321 |
| 11162552 | 19800266 | 18374575 | 12070128 | 20094046 |
| 8107774 | 12220513 | 16939971 | 12194817 | 21252999 |
| 12635176 | 10837065 | 18269210 | 10411507 | 27221039 |
| 19737400 | 11132103 | 20351173 | 15226403 | 15142631 |
| 8769096 | 12588352 | 8617720 | 33411331 | 36191627 |
| 11689693 | 20339092 | 21686281 | 11121404 | 10404636 |
| 19150423 | 20876458 | 23770676 | 29054983 | 10459005 |
| 9308965 | 18855616 | 11114305 | 23889475 | 12297041 |
| 3972795 | 21129147 | 18267069 | 29374774 | 30314968 |
| 1868835 | 18316480 | 20818427 | 17274988 | 29967253 |
| 1535687 | 21725049 | 19218920 | 12122009 | 21638049 |
| 9705341 | 12576332 | 18411265 | 11607814 | 31292270 |
| 9776743 | 22777353 | 8234276 | 30061385 | 18978815 |
| 20659008 | 2196092 | 21979375 | 23777859 | 26375550 |
| 18482001 | 16628190 | 8688489 | 32442405 | 22908275 |
| 3837185 | 7787779 | 16141325 | 24125182 | 30449325 |
| 18442490 | 25394790 | 12427531 | 16317043 | 19047365 |
| 18378697 | 25749719 | 12034743 | 33641239 | 12807903 |
| 9649439 | 22012064 | 12556884 | 34646012 | 25501036 |
| 16481321 | 18787208 | 20033988 | 9053841 | 17545517 |
| 18690213 | 11230154 | 32344597 | 23583981 | 9275162 |
| 10101234 | 19585539 | 25559847 | 22745804 | 24374327 |
| 9531558 | 9182763 | 32006019 | 25204415 | 24050403 |
| 15207230 | 15107406 | 32229307 | 24135138 | 12114214 |
| 15207236 | 22164214 | 14752036 | 18974108 | 26318331 |
| 17993265 | 12814943 | 28655883 | 20526349 | 18721752 |
| 20643813 | 18641354 | 32936320 | 24345920 | 10508857 |
| 22642575 | 21880786 | 20107438 | 17127394 | 19193720 |
| 12114023 | 27317874 | 16944206 | 7512720 | 1706616 |
| 24369424 | 23389849 | 27410728 | 20237318 | 22308027 |
| 25176256 | 26823759 | 24239741 | 18400509 | 24218551 |
| 3418702 | 26091038 | 22045002 | 26956865 | 7929212 |
| 26410597 | 7891679 | 10821263 | 26617336 | 22186019 |
| 8674114 | 14765127 | 27916698 | 16019425 | 8617750 |
| 23391857 | 19389811 | 28484373 | 16188692 | 16978839 |
| 24121435 | 11038178 | 26052920 | 8197119 | 27164096 |
| 22122054 | 25865393 | 30082454 | 23656643 | 28629526 |
| 24490730 | 12637528 | 26188105 | 12060844 | 24918289 |
| 19917616 | 16738314 | 30638948 | 7542747 | 29393890 |
| 22065625 | 20599712 | 30181353 | 17127320 | 26549969 |
| 17894445 | 16135801 | 30082452 | 23994464 | 25426117 |
| 15193314 | 26414766 | 30291150 | 22053285 | 27490140 |
| 25775268 | 10228168 | 26229154 | 9065403 | 22431075 |
| 11862453 | 18579750 | 29917041 | 11375957 | 21307776 |
| 20174677 | 8429029 | 27565685 | 9065402 | 19247306 |
| 21956940 | 11865982 | 30747242 | 11689702 | 15863038 |
| 25590339 | 12042069 | 22890707 | 10935490 | 15567848 |
| 10533284 | 2485589 | 16873410 | 21367659 | 7917292 |
| 16873067 | 19659435 | 25096182 | 20226962 | 16140923 |

| | | | | |
|---|---|---|---|---|
| 24588013 | 20439427 | 15574878 | 19927149 | 23139211 |
| 18056464 | 17937601 | 12530964 | 12787574 | 8553070 |
| 26134678 | 22289780 | 15472075 | 12671646 | 2418960 |
| 23298836 | 21677426 | 803282 | 25619719 | 22710176 |
| 23175443 | 22249808 | 9207787 | 24034695 | 7524079 |
| 23558953 | 20198122 | 21378989 | 16062169 | 17052215 |
| 25535896 | 23579596 | 21378985 | 23254191 | 1375264 |
| 22837387 | 20553370 | 16773578 | 15207811 | 2011584 |
| 20566684 | 21527729 | 11187898 | 22089421 | 19249086 |
| 25294908 | 23565489 | 21681853 | 29408302 | 4589993 |
| 26425723 | 21331644 | 11466430 | 25880691 | 20192804 |
| 9207791 | 20029269 | 16508304 | 15207812 | 8580068 |
| 8845844 | 22211890 | 10880446 | 29964125 | 15585754 |
| 15448004 | 21713385 | 25164209 | 8940083 | 18710302 |
| 11830541 | 23562747 | 25080992 | 23352614 | 22952392 |
| 17525745 | 20600241 | 11891328 | 21205088 | 10858654 |
| 16012724 | 21622208 | 11182080 | 18719096 | 20576942 |
| 19809159 | 22448029 | 22030550 | 22547065 | 19530173 |
| 8640234 | 21147620 | 21890822 | 21847121 | 10700176 |
| 9539778 | 22991412 | 21349854 | 25019382 | 22438566 |
| 21160078 | 19158955 | 11925448 | 26247733 | 7749328 |
| 10471491 | 27170215 | 16778208 | 24788754 | 6318122 |
| 8798788 | 22578774 | 18339869 | 18172307 | 17143297 |
| 11429702 | 18483217 | 11438922 | 25033798 | 17510365 |
| 9865713 | 17555450 | 29146722 | 27301673 | 18311776 |
| 19066716 | 8168137 | 11344305 | 22895187 | 16339511 |
| 28751768 | 20889717 | 11119607 | 22575959 | 10818153 |
| 16761019 | 20164854 | 14701863 | 20495530 | 10608864 |
| 29025582 | 19996456 | 18836481 | 27578174 | 16597699 |
| 31066629 | 1846706 | 23237809 | 18363568 | 12883891 |
| 27795560 | 16461907 | 25511451 | 19741192 | 23506886 |
| 17268528 | 18614017 | 25041739 | 26586761 | 12568659 |
| 30559310 | 15180923 | 26801977 | 22399681 | 25819438 |
| 24854988 | 19881549 | 26387946 | 16475976 | 16981043 |
| 19808698 | 25992381 | 9856458 | 16483568 | 14624363 |
| 24608088 | 10734310 | 8752100 | 7688988 | 18523892 |
| 10698507 | 10648629 | 29162704 | 17350321 | 14579113 |
| 28470536 | 15070743 | 27654855 | 19648010 | 10334869 |
| 21359601 | 15218027 | 25385546 | 19762341 | 15715662 |
| 15178581 | 7854453 | 11520916 | 21550381 | 16211368 |
| 14984498 | 12948444 | 8621434 | 16636301 | 12719981 |
| 11290575 | 15035987 | 19573017 | 30184463 | 9096318 |
| 14759363 | 23352452 | 17686986 | 12150994 | 18568018 |
| 15549094 | 22510884 | 9078261 | 17936557 | 14657019 |
| 20868367 | 31059601 | 1653651 | 9809982 | 11274402 |
| 20452772 | 24211266 | 17967490 | 19834456 | 10022869 |
| 21224387 | 27264673 | 7961713 | 7777546 | 12592392 |
| 20412773 | 25137548 | 25196463 | 21358634 | 10934479 |
| 21233212 | 11044625 | 11253356 | 7761852 | 17716241 |
| 11278283 | 9244301 | 10833508 | 8752208 | 14770312 |
| 20951342 | 14980219 | 11114882 | 9214508 | 12750889 |

| | | | | |
|---|---|---|---|---|
| 12759754 | 15774861 | 14744778 | 18391940 | 14729977 |
| 11279194 | 6305978 | 26832794 | 20378837 | 14623878 |
| 15522866 | 17597759 | 7891714 | 15059920 | 15126337 |
| 23453951 | 27002219 | 9687504 | 23236473 | 11463392 |
| 18725941 | 11969205 | 15870067 | 20531387 | 10733530 |
| 22438257 | 8078928 | 11100470 | 12897130 | 11511362 |
| 15498859 | 721806 | 1333888 | 9712883 | 11919562 |
| 23103841 | 25352115 | 7693660 | 18337750 | 19040420 |
| 15003515 | 23971743 | 11779505 | 19276069 | 11054413 |
| 26716518 | 164179 | 15564041 | 17146433 | 15377670 |
| 20807551 | 18718904 | 12482908 | 22252318 | 23059823 |
| 16219661 | 12359733 | 7737999 | 16140933 | 11382926 |
| 21932071 | 17170135 | 15761148 | 9671765 | 19615968 |
| 12907679 | 21173190 | 22074927 | 11896574 | 15781865 |
| 18391176 | 12216837 | 21350678 | 14968113 | 9858527 |
| 15728426 | 12221122 | 9821950 | 11814693 | 22012619 |
| 16460286 | 2332429 | 10653472 | 11559530 | 26773055 |
| 19783045 | 10805725 | 22074928 | 14688482 | 11350971 |
| 19278523 | 18524850 | 15384919 | 11278253 | 11988847 |
| 22277752 | 17556371 | 20577264 | 15580302 | 25429106 |
| 22934072 | 11733541 | 23236459 | 10973264 | 12771027 |
| 22394562 | 11279162 | 9724637 | 17284521 | 10984493 |
| 20890285 | 10559940 | 17332413 | 24739573 | 20096807 |
| 19926846 | 14527956 | 11191051 | 22802528 | 22988298 |
| 20871604 | 16399794 | 11566266 | 12692263 | 18187580 |
| 20107183 | 10224048 | 16923393 | 18431400 | 12170774 |
| 23604073 | 22647598 | 18365233 | 11574543 | 23139867 |
| 13175187 | 12646134 | 12612641 | 11812999 | 9063748 |
| 15173218 | 19896968 | 22421964 | 16772293 | 11971958 |
| 15173217 | 12536145 | 15814722 | 7784094 | 7834749 |
| 19923268 | 9113989 | 20019748 | 10891498 | 10807576 |
| 22732495 | 22984005 | 21737329 | 22802529 | 15607964 |
| 22918591 | 17928588 | 14739303 | 23485469 | 22172947 |
| 22812528 | 19074620 | 12181749 | 20460379 | 24096481 |
| 17031663 | 17263764 | 22017430 | 9780002 | 10524633 |
| 18031228 | 11564718 | 21173796 | 20817677 | 16249378 |
| 16095902 | 1499644 | 12893815 | 26122615 | 14614825 |
| 19616654 | 9989491 | 15950447 | 7566179 | 11070172 |
| 21892772 | 10375551 | 15976490 | 23703390 | 16341217 |
| 15209380 | 14685173 | 9926927 | 12606722 | 16857668 |
| 20164921 | 4754905 | 16728594 | 10777207 | 19521997 |
| 16608850 | 4507727 | 18198340 | 16135520 | 9799232 |
| 23441475 | 21550412 | 24366874 | 15577913 | 16144834 |
| 23026158 | 18321209 | 19822145 | 15273740 | 15226270 |
| 20551227 | 10727395 | 13298683 | 22766503 | 18420935 |
| 8162991 | 8988173 | 12203114 | 23365256 | 22085962 |
| 24868212 | 22371483 | 18344994 | 15289308 | 22884633 |
| 23736009 | 9162057 | 10716435 | 15735003 | 9501179 |
| 19037882 | 10648798 | 15665293 | 17442733 | 9704927 |
| 21736948 | 9850080 | 19439913 | 20353938 | 12004135 |
| 20565253 | 22735262 | 17540029 | 18676979 | 23954429 |

| | | | | |
|---|---|---|---|---|
| 21596310 | 15824739 | 11672531 | 22721435 | 19531029 |
| 24916645 | 21637926 | 23166394 | 19170759 | 2295606 |
| 19843474 | 24384674 | 19480567 | 22772988 | 27155576 |
| 21829232 | 16936726 | 19128516 | 20417083 | 18436705 |
| 26344767 | 18852463 | 21044011 | 28602209 | 16365312 |
| 24636259 | 22961379 | 22909387 | 21405107 | 27067637 |
| 21716289 | 22722852 | 15493035 | 19389739 | 17335404 |
| 16080119 | 23150874 | 22463982 | 21216617 | 11104681 |
| 12473678 | 15643423 | 22443931 | 17289573 | 15353562 |
| 24555100 | 22445173 | 16978057 | 8404537 | 17313374 |
| 20392956 | 25131816 | 23306458 | 12734757 | 16336962 |
| 22679399 | 24468964 | 22083510 | 18690016 | 15298169 |
| 21425435 | 24680896 | 17068183 | 17581921 | 16806605 |
| 15696166 | 23882114 | 20207225 | 12845533 | 16011460 |
| 9620804 | 19771207 | 19755485 | 17693766 | 20430952 |
| 15156186 | 16420676 | 18029452 | 18195086 | 21111842 |
| 23045283 | 24469049 | 23248145 | 16461757 | 22658984 |
| 23442798 | 15475956 | 18035408 | 18400693 | 8382952 |
| 12732176 | 20028861 | 20027182 | 12110519 | 19153612 |
| 23613270 | 22606298 | 20300647 | 12838422 | 16537653 |
| 21906677 | 25633035 | 12923055 | 18767960 | 16782012 |
| 9566865 | 26208636 | 22394165 | 18682559 | 15220349 |
| 25123279 | 21679462 | 22122341 | 764868 | 10553003 |
| 12217689 | 20833819 | 28472341 | 5324173 | 17199046 |
| 19350569 | 10948449 | 19647095 | 24315934 | 12958364 |
| 23148227 | 11493559 | 9774680 | 25692999 | 16678095 |
| 11257229 | 8898218 | 11278460 | 24581449 | 21768372 |
| 22189971 | 18031719 | 24240231 | 24898039 | 16678096 |
| 11545733 | 30590909 | 22964823 | 25780173 | 19837038 |
| 15030764 | 10913181 | 30590466 | 25304425 | 21725318 |
| 26404249 | 9743535 | 16046443 | 25806043 | 17108000 |
| 26060017 | 24751955 | 18445271 | 25734984 | 23260141 |
| 18438928 | 16531405 | 20054179 | 22137969 | 22653731 |
| 9794229 | 29185068 | 8970730 | 21854988 | 17141155 |
| 19020999 | 27558325 | 17956727 | 21910628 | 21554866 |
| 24924170 | 22432088 | 17475546 | 21417719 | 22017973 |
| 17142326 | 19699815 | 16054042 | 20716516 | 10990458 |
| 27272193 | 20924107 | 7531665 | 25755220 | 19561074 |
| 10816326 | 20203268 | 17011499 | 16040803 | 18093802 |
| 22189423 | 15701714 | 17060461 | 22426497 | 9214626 |
| 15574200 | 24584857 | 18518822 | 21572561 | 15935773 |
| 16164020 | 25940801 | 16380219 | 16364630 | 11842111 |
| 24880459 | 21725048 | 10603305 | 15879697 | 18667602 |
| 18676844 | 22307729 | 15860371 | 20101632 | 26316498 |
| 16804544 | 29593731 | 21082419 | 12045094 | 11438699 |
| 23334789 | 17934488 | 3466163 | 17482885 | 17166836 |
| 21734270 | 16732331 | 10369688 | 37508007 | 12885641 |
| 19800882 | 19796237 | 24648516 | 32707086 | 31883789 |
| 21203558 | 20887958 | 11058110 | 38425362 | 26111660 |
| 18184866 | 29431622 | 11006276 | 11051212 | 24344280 |
| 21706051 | 20581311 | 12145297 | 34471290 | 11278378 |

| | | | | |
|---|---|---|---|---|
| 8238642 | 10477748 | 15146457 | 25084529 | 20074548 |
| 28715957 | 9473669 | 30901624 | 32893956 | 32296183 |
| 10023774 | 17868694 | 11201742 | 10473610 | 9692923 |
| 26797144 | 9407135 | 11705709 | 10321247 | 18235502 |
| 29395074 | 9837938 | 20036641 | 10601307 | 17494760 |
| 17001303 | 17307141 | 32910366 | 19745164 | 20538072 |
| 21109473 | 12808044 | 12909588 | 16767699 | 18235501 |
| 11748222 | 12169098 | 16799563 | 30368668 | 14749374 |
| 23210908 | 10856295 | 12023038 | 27813118 | 11087735 |
| 11080163 | 10026224 | 19247433 | 12909369 | 12102554 |
| 12692302 | 8557660 | 12533514 | 23455478 | 15563450 |
| 9545647 | 8128486 | 7961848 | 9700206 | 26044572 |
| 9742218 | 28279197 | 10024240 | 24748541 | 23317503 |
| 27224062 | 10506173 | 16483939 | 29393141 | 16848710 |
| 11784712 | 10575001 | 9790763 | 9242461 | 31285543 |
| 9343427 | 16079250 | 18003636 | 11891283 | 20170512 |
| 29425100 | 12953056 | 11543634 | 31127867 | 24309898 |
| 11606575 | 9792688 | 23231787 | 9346935 | 16940157 |
| 12032137 | 26066539 | 19478182 | 19778628 | 23415904 |
| 9804823 | 28174279 | 19011233 | 28625565 | 30340023 |
| 30590535 | 9707407 | 9731536 | 21555521 | 10944470 |
| 9710638 | 29856954 | 12422217 | 24263804 | 15095019 |
| 28847827 | 23807634 | 29769719 | 12718547 | 25819840 |
| 31916624 | 16738128 | 12223416 | 29909984 | 31698146 |
| 25134449 | 12865409 | 15044604 | 11394904 | 20615952 |
| 10727212 | 10334923 | 29454968 | 12719432 | 24664998 |
| 9616160 | 26844272 | 15175244 | 1718748 | 21229319 |
| 9628887 | 21383261 | 29305086 | 15208306 | 23673625 |
| 19124460 | 30366904 | 7882974 | 25406032 | 21048217 |
| 18715871 | 12578837 | 19995937 | 9677374 | 21874273 |
| 24038671 | 17412710 | 16556915 | 20333297 | 21813643 |
| 31784983 | 23454892 | 22140376 | 30845223 | 14973125 |
| 23213474 | 16399079 | 31276219 | 18775313 | 12082086 |
| 23161582 | 30502085 | 12049641 | 22566699 | 26635000 |
| 12912983 | 17046230 | 17000704 | 16288044 | 25851604 |
| 11279118 | 23682772 | 16428860 | 23532176 | 8377829 |
| 26598493 | 20122914 | 9063746 | 11027676 | 9685409 |
| 11146551 | 24904170 | 24449907 | 25902869 | 18529014 |
| 18634786 | 15736167 | 16251970 | 28428259 | 9012831 |
| 27637550 | 15996652 | 24067371 | 7556058 | 11290329 |
| 27526206 | 8887666 | 23055042 | 11545740 | 23564352 |
| 15556292 | 24675081 | 11832950 | 8647288 | 26239904 |
| 16814779 | 16109395 | 9598310 | 28659385 | 10373551 |
| 15805470 | 14559152 | 9710624 | 21177766 | 18398435 |
| 11593007 | 10779340 | 15764604 | 1851997 | 31722427 |
| 25725067 | 15821743 | 18357469 | 22537386 | 24086678 |
| 23275342 | 29719408 | 26068709 | 9465039 | 16885160 |
| 7758584 | 11058098 | 23284264 | 19203578 | 10648599 |
| 32295885 | 15561719 | 18845538 | 15378014 | 19092802 |
| 12177006 | 32454406 | 16267379 | 25586960 | 31428936 |
| 19946469 | 23332760 | 7616957 | 23960073 | 9516461 |

| | | | | |
|---|---|---|---|---|
| 17900573 | 29989768 | 11409944 | 164477 | 7681362 |
| 12963706 | 21865368 | 12067245 | 21802064 | 15544030 |
| 25393282 | 16826530 | 11440964 | 17386955 | 21908933 |
| 30318146 | 21639948 | 10506113 | 23423488 | 26235620 |
| 23091057 | 18419792 | 16815959 | 21131358 | 15610735 |
| 24308962 | 30742913 | 16385575 | 15295589 | 16244323 |
| 12493763 | 19956441 | 16970545 | 20486779 | 21310851 |
| 12930902 | 17660942 | 16584116 | 22285895 | 24852694 |
| 9751713 | 23475188 | 20681653 | 17612493 | 22569073 |
| 11698644 | 14747765 | 15590648 | 16529745 | 19158095 |
| 17432114 | 29279276 | 15039226 | 24797360 | 20412774 |
| 12514734 | 31034780 | 15020646 | 20417621 | 9228007 |
| 20375010 | 12379743 | 21829704 | 18829576 | 23802099 |
| 15543136 | 8985388 | 18579364 | 15864339 | 21453770 |
| 14530263 | 12089062 | 19455135 | 10830966 | 22927400 |
| 9384587 | 8591043 | 22678861 | 22921398 | 27410263 |
| 9334247 | 9108473 | 12581329 | 15363492 | 16493415 |
| 27509850 | 15371627 | 22790079 | 27313835 | 11875109 |
| 12393887 | 10567268 | 22302197 | 18782350 | 23261442 |
| 24222088 | 15818401 | 18547992 | 18434550 | 23516131 |
| 25569233 | 17116881 | 23437201 | 20048743 | 20117946 |
| 19001379 | 17195095 | 19857756 | 17055782 | 10692429 |
| 29604308 | 9616213 | 20067622 | 29343764 | 26569053 |
| 10546895 | 3027969 | 27754373 | 18328803 | 17234883 |
| 12239342 | 10196249 | 11329262 | 9472019 | 8702689 |
| 24488492 | 11779719 | 26668369 | 19245654 | 20698033 |
| 9228079 | 10867028 | 9452426 | 2567185 | 9094314 |
| 12766176 | 11679155 | 26855069 | 16364253 | 8826975 |
| 12192049 | 11762996 | 12917011 | 12740371 | 11803135 |
| 12171929 | 15803138 | 27339894 | 16105857 | 26490400 |
| 27789755 | 9727492 | 21159775 | 19596898 | 19683471 |
| 14623887 | 9036860 | 1238395 | 16479585 | 26387753 |
| 8093561 | 9971843 | 7217084 | 14651849 | 16916938 |
| 22792322 | 20447565 | 10898110 | 19098422 | 15581595 |
| 11438732 | 9321539 | 11294878 | 11069105 | 25720603 |
| 12764197 | 7511174 | 26545797 | 12392881 | 23150577 |
| 20588296 | 8181167 | 16216911 | 12502487 | 21440577 |
| 10642537 | 2871380 | 26587646 | 15661536 | 24379912 |
| 17255108 | 15546794 | 25577493 | 16027121 | 26499800 |
| 6867732 | 9770555 | 21636783 | 24135279 | 19324998 |
| 17376010 | 16170679 | 11360992 | 21569822 | 24262987 |
| 17464936 | 10756043 | 2339109 | 20033380 | 26254015 |
| 15134803 | 19762681 | 240819 | 11025665 | 16061178 |
| 18729003 | 11573093 | 19661994 | 18791498 | 10446062 |
| 16679330 | 15177891 | 26976652 | 12831062 | 19372258 |
| 15174896 | 22160858 | 1123350 | 15778410 | 8891199 |
| 16374430 | 28912259 | 21586563 | 12539168 | 17785634 |
| 18417113 | 19701189 | 8075637 | 9560344 | 22155301 |
| 18035185 | 11031233 | 18707589 | 17395877 | 33262481 |
| 18449520 | 18949601 | 14640697 | 20045007 | 32529116 |
| 12921235 | 12724730 | 23800242 | 11397779 | 32277040 |

| | | | | |
|---|---|---|---|---|
| 32321524 | 7834621 | 3266253 | 18760377 | 9735376 |
| 31967327 | 10777214 | 16940415 | 16682204 | 19075563 |
| 28510041 | 9380030 | 10075741 | 10340377 | 16183742 |
| 20716671 | 16679317 | 15322122 | 7554031 | 11665719 |
| 33091573 | 21846477 | 15790560 | 22435563 | 20493860 |
| 16382109 | 956169 | 12727795 | 16360037 | 18075314 |
| 32142651 | 10074425 | 19617630 | 7948950 | 16582605 |
| 32283108 | 10224120 | 11991950 | 22435546 | 22854598 |
| 20600852 | 19474220 | 12573484 | 16101442 | 18804552 |
| 19617398 | 23512538 | 20399894 | 2147521 | 21737170 |
| 29737559 | 19203995 | 11120824 | 23425920 | 23416979 |
| 32438371 | 18607004 | 15541765 | 23150759 | 12726855 |
| 20457564 | 9704006 | 17901129 | 22435545 | 16360026 |
| 32680882 | 19366855 | 21124311 | 21541195 | 23296650 |
| 27825853 | 18089838 | 16545780 | 20004154 | 19181515 |
| 26551702 | 18336843 | 19106115 | 23263379 | 8617251 |
| 32711925 | 19255421 | 17952121 | 22083606 | 23239948 |
| 25904598 | 19158084 | 14978251 | 19230643 | 9108147 |
| 30101215 | 17934341 | 17595319 | 22260696 | 17301840 |
| 17404574 | 2843537 | 24261707 | 11859406 | 10887490 |
| 19079265 | 17709388 | 17266942 | 12214244 | 16314342 |
| 17914240 | 12223212 | 15340055 | 1653904 | 16258277 |
| 10618704 | 11114727 | 18288955 | 23238565 | 15603758 |
| 24743243 | 22167199 | 15501915 | 14599770 | 22263797 |
| 20808857 | 11502724 | 17284330 | 18424430 | 20679392 |
| 9821948 | 17223712 | 11903058 | 12214253 | 23002429 |
| 15078873 | 11259609 | 16197558 | 22588366 | 20354226 |
| 10224227 | 10395741 | 16289099 | 23721719 | 15254416 |
| 9799222 | 9504046 | 12573486 | 1535095 | 22980457 |
| 14512773 | 8910454 | 12611900 | 8985253 | 22410433 |
| 10934467 | 22903824 | 20619336 | 23747889 | 20061149 |
| 10982831 | 20395535 | 15598615 | 21807066 | 17581279 |
| 14555980 | 14597617 | 19471584 | 21187855 | 9363685 |
| 17164131 | 9712901 | 17944540 | 23638217 | 11595834 |
| 17494091 | 9488039 | 9099515 | 15947787 | 23802008 |
| 17975109 | 8937476 | 11343253 | 19125251 | 19075564 |
| 15180494 | 22659247 | 15004031 | 11089869 | 7488937 |
| 3619897 | 19574409 | 19129222 | 2142119 | 23303139 |
| 18216721 | 15837795 | 15897893 | 17935492 | 10037143 |
| 16258029 | 10398686 | 18848820 | 15865930 | 19586837 |
| 4051502 | 1844873 | 15489539 | 16115458 | 22160080 |
| 15748653 | 11470802 | 15496506 | 16082221 | 20515729 |
| 16399376 | 7557864 | 12120277 | 15686624 | 21903324 |
| 3977905 | 10620335 | 21226706 | 17151078 | 18677098 |
| 6548162 | 11097088 | 21420387 | 19582216 | 12207176 |
| 2834948 | 8396713 | 28760335 | 9261102 | 8948422 |
| 16179585 | 21856752 | 22073591 | 16398215 | 22522501 |
| 18383502 | 18827440 | 23150753 | 15136767 | 15147722 |
| 10759591 | 10961990 | 18047734 | 18508566 | 15082921 |
| 24467436 | 8120057 | 20813042 | 21722344 | 9681884 |
| 7889851 | 6095293 | 10362249 | 11832478 | 23519070 |

| | | | | |
|---|---|---|---|---|
| 9466696 | 20305300 | 29653253 | 31540582 | 9822657 |
| 22038529 | 12963843 | 10893433 | 32850791 | 18184567 |
| 8301538 | 19890334 | 16376858 | 31843468 | 28473751 |
| 18796370 | 10526213 | 9218780 | 20945369 | 29249655 |
| 17622968 | 19683496 | 12481981 | 16984440 | 24741076 |
| 16998506 | 23788429 | 20301396 | 7485389 | 26103054 |
| 22911014 | 23308042 | 8930409 | 17644256 | 29126901 |
| 20023404 | 8782982 | 22116691 | 34017408 | 21193401 |
| 16024119 | 15254415 | 25514926 | 15296246 | 29038536 |
| 22743550 | 8666399 | 24122582 | 16621453 | 27499160 |
| 12655298 | 16397206 | 22417847 | 34149889 | 19073886 |
| 22265392 | 21034966 | 22899908 | 9588880 | 19426232 |
| 23388117 | 22084390 | 7741998 | 31974341 | 22116087 |
| 20224865 | 19581929 | 23142810 | 32102389 | 29339377 |
| 18940270 | 20006736 | 19369394 | 7999272 | 21904389 |
| 12045216 | 20445224 | 15621726 | 28771809 | 8596936 |
| 19400965 | 21535261 | 3798106 | 27648692 | 1191089 |
| 18497029 | 10092119 | 23220880 | 12618512 | 16551244 |
| 8533012 | 10582242 | 26028978 | 21538580 | 30511409 |
| 21089513 | 8533157 | 15805248 | 17869227 | 29347993 |
| 19707781 | 15670890 | 27251275 | 23792675 | 28990585 |
| 8939883 | 9418909 | 20453058 | 25087956 | 22437870 |
| 21963855 | 23200932 | 18676830 | 24972246 | 28643244 |
| 19686080 | 21593588 | 28348404 | 21602788 | 30345906 |
| 12660173 | 19023332 | 17397528 | 18184568 | 23108542 |
| 23717325 | 15723711 | 20570901 | 24225153 | 27106177 |
| 15483403 | 18243065 | 16385346 | 24859235 | 26187180 |
| 15865942 | 16903208 | 15086769 | 27449752 | 16672367 |
| 17626635 | 22844074 | 12373339 | 28634229 | 26218440 |
| 23554604 | 12963833 | 26794609 | 19812359 | 21704641 |
| 10713716 | 17611403 | 16217483 | 23027865 | 20392245 |
| 22704343 | 22515271 | 33073427 | 23103546 | 23748100 |
| 21326949 | 23808152 | 22092795 | 18555543 | 29587428 |
| 18368919 | 15254433 | 10469310 | 26071486 | 16936699 |
| 20465793 | 22761618 | 27477081 | 25530759 | 22226352 |
| 12171872 | 23208375 | 11573241 | 15689566 | 15680226 |
| 10673629 | 21391908 | 23226515 | 28608766 | 15922018 |
| 15496581 | 21615334 | 10550323 | 26585419 | 2203193 |
| 23070009 | 30081750 | 2656805 | 28624624 | 7720105 |
| 22991823 | 28933638 | 27256813 | 27922662 | 18423897 |
| 17251377 | 30501132 | 17576240 | 29133590 | 16112078 |
| 22150313 | 30140075 | 15953044 | 28642704 | 16402899 |
| 20158568 | 23877423 | 18597619 | 12386268 | 23951310 |
| 7606818 | 20186705 | 25837671 | 29431615 | 10640766 |
| 7672812 | 19590688 | 11685189 | 26824050 | 26244871 |
| 21329706 | 11952781 | 28182006 | 26750873 | 24161035 |
| 19364923 | 7889863 | 25876136 | 25616441 | 25212687 |
| 7787878 | 17702526 | 15625461 | 23313043 | 15212335 |
| 14593728 | 12510015 | 25998853 | 20357764 | 21795548 |
| 7587079 | 15660110 | 32104287 | 28624623 | 21173082 |
| 21196225 | 9118888 | 22848228 | 19073885 | 29127110 |

| | | | | |
|---|---|---|---|---|
| 29427249 | 24979721 | 22403545 | 19834513 | 19898483 |
| 9733515 | 27694978 | 7753047 | 11875025 | 20463033 |
| 20621048 | 28329682 | 22715882 | 25475423 | 25080474 |
| 16005139 | 21613227 | 6321612 | 25176654 | 30587505 |
| 7564239 | 22306293 | 30886620 | 22652185 | 22083728 |
| 20847235 | 23792563 | 32116493 | 8123671 | 17380161 |
| 19524112 | 24058770 | 31439935 | 9694901 | 14730303 |
| 14239091 | 8841154 | 16525119 | 9582303 | 19111245 |
| 16602100 | 16778732 | 24077738 | 8672428 | 15545627 |
| 3769199 | 10516633 | 31175934 | 16854371 | 15668327 |
| 6774165 | 9010622 | 29606485 | 12011101 | 16921403 |
| 12235098 | 12811366 | 27408775 | 19014978 | 16805667 |
| 25646736 | 12492606 | 9741627 | 22724020 | 15343279 |
| 20923771 | 1867957 | 27852311 | 27547445 | 14993903 |
| 23558541 | 18187562 | 31126986 | 27047494 | 15688066 |
| 9291139 | 12950465 | 32094408 | 18787044 | 17943134 |
| 22156476 | 11186130 | 31351098 | 29100090 | 17128209 |
| 12821112 | 18781855 | 23443243 | 17522159 | 17060944 |
| 20420946 | 15886284 | 28476236 | 10340754 | 15339658 |
| 25736321 | 15051713 | 28588114 | 9719154 | 28339062 |
| 8898652 | 8875123 | 17126425 | 16396903 | 18004398 |
| 14715079 | 2049245 | 33203880 | 12766769 | 20610534 |
| 18316791 | 18622261 | 31708432 | 23064016 | 22941656 |
| 10479724 | 8818573 | 27009876 | 28566479 | 8513149 |
| 19679400 | 12895198 | 29908837 | 29194579 | 8621488 |
| 25236910 | 1782973 | 27879284 | 22510445 | 17715138 |
| 27173435 | 16920476 | 30328953 | 16880404 | 17765923 |
| 28698599 | 18719619 | 26239609 | 12414726 | 16436514 |
| 7697895 | 12920168 | 17186031 | 29100091 | 9199932 |
| 27380651 | 15625333 | 17382285 | 9675033 | 19525936 |
| 26161337 | 17724700 | 18483225 | 15838507 | 11809771 |
| 25322271 | 12185559 | 18277979 | 28739660 | 7915517 |
| 29553041 | 15530129 | 20117961 | 9361030 | 15377654 |
| 9405715 | 32760717 | 21305127 | 15327782 | 15225546 |
| 24790347 | 19793304 | 22387373 | 11203700 | 12919958 |
| 9046958 | 26923589 | 10969042 | 27597235 | 19168442 |
| 27068427 | 30479058 | 26979667 | 26604140 | 19202061 |
| 13211659 | 23901111 | 29149350 | 16971658 | 19586908 |
| 28423456 | 32451529 | 2143188 | 14656760 | 18509536 |
| 8001864 | 22885700 | 32344011 | 17537792 | 12086603 |
| 27047663 | 18179280 | 27363989 | 12060755 | 9765279 |
| 26956190 | 23661693 | 2856554 | 17785448 | 11877377 |
| 12788846 | 21505029 | 21742792 | 17462874 | 12364621 |
| 29224098 | 20720586 | 11460506 | 15141091 | 8175912 |
| 27163392 | 10904115 | 3021194 | 10021334 | 19793862 |
| 1594605 | 28345259 | 19008457 | 29945868 | 11027291 |
| 8839934 | 33670154 | 32339221 | 19809516 | 18619531 |
| 16024935 | 28258193 | 24611772 | 11138002 | 19854139 |
| 19129847 | 17051160 | 11076529 | 9425907 | 10608837 |
| 12050213 | 18560619 | 11731805 | 24357607 | 19652551 |
| 21325058 | 33599083 | 15122900 | 12783789 | 16617241 |

| | | | | |
|---|---|---|---|---|
| 17460694 | 24104880 | 1840259 | 20360045 | 16633336 |
| 12447691 | 12215251 | 8524414 | 15886194 | 19996105 |
| 10556074 | 12011431 | 27067600 | 15190204 | 11865064 |
| 20871615 | 11369231 | 17525341 | 16810316 | 16009723 |
| 10499802 | 12814551 | 15640246 | 11239454 | 19111657 |
| 11916980 | 21283629 | 15870257 | 9733514 | 19151707 |
| 10884395 | 12676925 | 21044075 | 20413593 | 19995904 |
| 17525342 | 9705271 | 10364196 | 2882507 | 2564316 |
| 7961795 | 15317757 | 18006705 | 23007646 | 24293646 |
| 10078208 | 17049555 | 11537053 | 12354784 | 21978893 |
| 12717439 | 2050703 | 7606819 | 26675481 | 9590181 |
| 10373536 | 21659603 | 9716408 | 7526206 | 7700386 |
| 16862143 | 17478428 | 20079829 | 17621610 | 16530042 |
| 8146338 | 9564049 | 18319725 | 20023648 | 17939684 |
| 18001825 | 15205463 | 23166356 | 1578192 | 19328070 |
| 11157767 | 19193796 | 14976165 | 12199140 | 15279788 |
| 9278511 | 22154951 | 15680327 | 8665503 | 12147700 |
| 11051553 | 26512707 | 19567472 | 16899510 | 18212045 |
| 19966300 | 3025664 | 7958836 | 16628214 | 15064416 |
| 17828269 | 17194776 | 27723717 | 15574326 | 8632903 |
| 7939630 | 12766152 | 12402037 | 8991084 | 16712457 |
| 15897895 | 16547522 | 15221963 | 8769649 | 18596042 |
| 10078207 | 20171170 | 21857671 | 8640237 | 20364141 |
| 20061803 | 2946935 | 15064730 | 19652550 | 18172165 |
| 21419344 | 19338310 | 11202906 | 21475307 | 10999600 |
| 10713044 | 20064462 | 17558410 | 15671039 | 19473992 |
| 16980960 | 10373512 | 21639834 | 20348101 | 9828139 |
| 16707425 | 10205172 | 9435225 | 10744741 | 7585968 |
| 20020535 | 18172500 | 18206974 | 2406247 | 12939256 |
| 3142690 | 16189514 | 2195549 | 11573085 | 20445207 |
| 21120944 | 11432836 | 15527801 | 27342858 | 18172690 |
| 12724401 | 16360315 | 10981963 | 16199878 | 17704056 |
| 18082599 | 7969176 | 17412408 | 22508508 | 17312392 |
| 9660782 | 14734534 | 16236519 | 12228710 | 16260474 |
| 24235147 | 18443037 | 14695167 | 15199141 | 12890688 |
| 15806145 | 19344625 | 20826806 | 18974355 | 10679321 |
| 21872579 | 19203579 | 8170954 | 9518481 | 20173098 |
| 22867704 | 2142452 | 10910365 | 22442688 | 16787914 |
| 15899892 | 18001824 | 1332913 | 24627472 | 16153896 |
| 9822679 | 15356634 | 9836640 | 20643585 | 10660545 |
| 18519686 | 22237204 | 20634189 | 15485915 | 14636568 |
| 19661379 | 20231364 | 12192000 | 1811480 | 12781359 |
| 15940266 | 26196677 | 21172801 | 15272308 | 15201865 |
| 14695475 | 16943440 | 16531125 | 14636569 | 9637771 |
| 11986308 | 11971963 | 12384589 | 20102227 | 17227144 |
| 15122316 | 18298799 | 19261749 | 15299030 | 20639885 |
| 8521392 | 7838523 | 12419324 | 20046100 | 15454491 |
| 16116421 | 15195100 | 9030781 | 16205630 | 17599047 |
| 15115758 | 18644834 | 12470949 | 17643122 | 15661742 |
| 9685493 | 8843195 | 7961977 | 15314187 | 16698308 |
| 2573657 | 17396150 | 8964493 | 17608804 | 9388480 |

| | | | | |
|---|---|---|---|---|
| 11032027 | 16814252 | 23680151 | 14566050 | 21042587 |
| 11157805 | 19907496 | 9636169 | 12209014 | 21161613 |
| 17616665 | 17611284 | 24811749 | 24100029 | 12526805 |
| 17959650 | 16087684 | 10608806 | 8247533 | 15117943 |
| 12019152 | 20729858 | 10448035 | 17118716 | 9590180 |
| 19109555 | 9052673 | 24220101 | 7905912 | 14729973 |
| 16908529 | 17088560 | 12191481 | 16595695 | 9207062 |
| 12414623 | 2243768 | 15377652 | 15451423 | 21319273 |
| 15096610 | 12824158 | 16142238 | 18660752 | 8640235 |
| 11163187 | 12857880 | 9889122 | 15096578 | 12205100 |
| 22357538 | 8622991 | 19033441 | 18156970 | 21504906 |
| 14500819 | 9766667 | 17954613 | 12419808 | 10839544 |
| 831811 | 12080054 | 18066086 | 9572863 | 15138768 |
| 21285353 | 10329681 | 14988723 | 6273595 | 11784855 |
| 16247472 | 23345434 | 19283071 | 8119945 | 19097996 |
| 12578958 | 12883740 | 15315825 | 12750383 | 20656690 |
| 11018012 | 21242293 | 12794064 | 15616588 | 20147522 |
| 21149266 | 17643121 | 9933573 | 10684855 | 10777662 |
| 9452416 | 12649176 | 23030715 | 19446481 | 7545954 |
| 16713580 | 26778126 | 14517836 | 11090622 | 23849169 |
| 21789020 | 18442975 | 18356527 | 18832153 | 11025664 |
| 14871897 | 17563354 | 11454867 | 15199523 | 27428775 |
| 17314514 | 23467123 | 22908299 | 16141202 | 17767920 |
| 8702565 | 10468606 | 7792600 | 23361318 | 20671765 |
| 19535328 | 11331310 | 10477523 | 15707391 | 18775730 |
| 11057907 | 18510930 | 9311737 | 4006916 | 20639400 |
| 25813721 | 23525106 | 12447382 | 17634560 | 7697716 |
| 9699634 | 10480872 | 18417535 | 17452773 | 19440044 |
| 16365875 | 16546998 | 19589784 | 11418864 | 17588522 |
| 7115720 | 14681192 | 24332808 | 17611581 | 11551919 |
| 10551855 | 8990123 | 17923702 | 12167711 | 16731526 |
| 11350926 | 10673031 | 15279789 | 11301010 | 17115032 |
| 17081985 | 18285803 | 21362556 | 11239453 | 2565339 |
| 12810625 | 15456891 | 17273969 | 15811850 | 17030982 |
| 22013166 | 8969240 | 24526736 | 11395493 | 21309033 |
| 12446782 | 12852856 | 8943031 | 9430682 | 17495531 |
| 21798247 | 19024604 | 16581787 | 6704953 | 18550849 |
| 21558276 | 25609649 | 7923193 | 26582912 | 21501958 |
| 10391891 | 10888888 | 16371510 | 10498869 | 12239151 |
| 8837778 | 23540691 | 16377563 | 9651580 | 19197159 |
| 15743907 | 12612651 | 19579266 | 22886304 | 14742437 |
| 25465621 | 11863428 | 15247280 | 22589541 | 14559997 |
| 23083810 | 26253028 | 19270065 | 22002537 | 22517901 |
| 9461304 | 15220350 | 16285702 | 16049003 | 15592449 |
| 10552928 | 20829486 | 10359610 | 9038370 | 18048416 |
| 11459832 | 23364835 | 20936109 | 8589730 | 21198351 |
| 16479174 | 10364235 | 19464297 | 12414651 | 8804307 |
| 12734188 | 18372919 | 14734805 | 11842105 | 15111055 |
| 9878247 | 9925639 | 22139841 | 11537315 | 9679063 |
| 12228248 | 11371615 | 11821419 | 15694335 | 16027118 |
| 12140561 | 20304803 | 26133775 | 24485656 | 16467875 |

| | | | | |
|---|---|---|---|---|
| 12504096 | 15782130 | 12704184 | 11106734 | 17942393 |
| 11278446 | 18309293 | 7774019 | 19444312 | 10973490 |
| 7957065 | 18480403 | 8621570 | 12556559 | 15196461 |
| 8589678 | 20932174 | 15775963 | 15361825 | 20424263 |
| 6602790 | 12537559 | 1531147 | 8566796 | 12654198 |
| 25933514 | 14983014 | 10208430 | 19788416 | 14743218 |
| 9450543 | 12370410 | 16537486 | 17157251 | 7715730 |
| 18931676 | 16818604 | 15659650 | 19483192 | 11331603 |
| 12145306 | 18283122 | 19160488 | 11447121 | 15342490 |
| 18519640 | 15650047 | 2139805 | 11114745 | 11550094 |
| 15280377 | 20009512 | 18644861 | 9295282 | 11454856 |
| 11836499 | 14605214 | 19759395 | 11314038 | 10464290 |
| 16431910 | 16794254 | 17178852 | 16122425 | 14636574 |
| 9427750 | 15279791 | 10563794 | 15650050 | 9121459 |
| 20195506 | 10859164 | 16223874 | 15989956 | 18449195 |
| 11063725 | 17553757 | 12773400 | 19465921 | 12782307 |
| 15226314 | 16601680 | 19556969 | 14583606 | 11604499 |
| 12402044 | 17638878 | 11726552 | 18025084 | 9351817 |
| 10526407 | 15364927 | 15538388 | 7715731 | 10950869 |
| 16912045 | 18077418 | 20061386 | 7596430 | 21199877 |
| 12709442 | 15235112 | 24079363 | 12814430 | 17102637 |
| 27611684 | 18317453 | 20705237 | 20212043 | 11583998 |
| 20360007 | 11077446 | 17101782 | 9045680 | 19629043 |
| 12607004 | 19185524 | 21056556 | 11333291 | 16288057 |
| 20858735 | 20729856 | 16793542 | 23521171 | 8946918 |
| 9050866 | 19793861 | 16311512 | 11053413 | 11931755 |
| 20413589 | 16622405 | 11741547 | 7594449 | 16438930 |
| 19423707 | 11325820 | 16478997 | 10206961 | 26802432 |
| 14724280 | 10320477 | 12046007 | 15936993 | 9751706 |
| 18594563 | 11687627 | 20724660 | 9267021 | 15618521 |
| 18848520 | 18418389 | 15314022 | 15310756 | 9168116 |
| 17124492 | 23715498 | 1393148 | 12576443 | 12861053 |
| 18158334 | 18239466 | 15960976 | 16510573 | 11298456 |
| 12607005 | 17804464 | 9822680 | 18285460 | 8358790 |
| 9649500 | 19261748 | 20383123 | 16930133 | 11252893 |
| 16086026 | 12379650 | 21145460 | 10608812 | 19898529 |
| 18948756 | 22973052 | 11721054 | 18662573 | 19605351 |
| 19202191 | 18343821 | 8610130 | 18566590 | 18948948 |
| 23230272 | 15542852 | 11166174 | 8521816 | 11336668 |
| 20937773 | 21699228 | 8769132 | 11864614 | 18158288 |
| 12697768 | 17636252 | 8056767 | 15220930 | 17899380 |
| 9756909 | 19908865 | 20154705 | 15583028 | 11877376 |
| 8896563 | 9806842 | 9812896 | 11278964 | 17459151 |
| 18769153 | 11390642 | 17889669 | 9037071 | 11709054 |
| 15149598 | 17097061 | 14966270 | 8918887 | 10724175 |
| 23748380 | 17173041 | 18280240 | 9109492 | 17974916 |
| 3257477 | 16912307 | 17189255 | 14561771 | 19494828 |
| 17525340 | 12607003 | 16278218 | 15003516 | 15574335 |
| 17887956 | 12676583 | 21364637 | 16689829 | 10713175 |
| 16873062 | 2308592 | 17963495 | 19525978 | 19683501 |
| 10764811 | 18077395 | 15811628 | 18305112 | 15567177 |

| | | | | |
|---|---|---|---|---|
| 17110379 | 32486270 | 16973150 | 15542856 | 12015981 |
| 9802988 | 23838442 | 19666824 | 18403408 | 14519196 |
| 10611322 | 31612241 | 23643939 | 20805471 | 24929628 |
| 21104395 | 12045100 | 23022961 | 19544440 | 27740627 |
| 10681541 | 11257218 | 19825829 | 24823357 | 24384374 |
| 15949439 | 24409201 | 23123965 | 24576030 | 25307053 |
| 19962312 | 23352243 | 23677624 | 28041631 | 29158817 |
| 10364241 | 17065211 | 16258936 | 1722028 | 21798082 |
| 11389439 | 21855102 | 16738056 | 32353859 | 26247089 |
| 12149244 | 26379663 | 19439460 | 31226023 | 24426196 |
| 16464007 | 25396298 | 21284982 | 21543844 | 30249788 |
| 10487762 | 25741013 | 19915529 | 11163209 | 30420806 |
| 9488720 | 23870315 | 11562345 | 22028656 | 28882953 |
| 15688006 | 23637409 | 20833363 | 19304306 | 32198194 |
| 9476899 | 24473128 | 26847180 | 30425656 | 25470695 |
| 14676842 | 25136083 | 27098840 | 21285253 | 20301485 |
| 10228148 | 25642836 | 27374873 | 29258163 | 22461740 |
| 19468298 | 19694547 | 23496208 | 32746883 | 31676439 |
| 15916964 | 25493356 | 15184648 | 26269761 | 14592533 |
| 21245467 | 16698996 | 17463250 | 23856032 | 23055695 |
| 16904321 | 11461707 | 22773810 | 24556840 | 31462513 |
| 15071507 | 16140752 | 23533263 | 23723077 | 29100061 |
| 10362363 | 17718913 | 11397666 | 23045246 | 28766509 |
| 11314011 | 26516900 | 25337673 | 25656897 | 32120838 |
| 20655466 | 19153231 | 22426421 | 10334664 | 30333156 |
| 9922454 | 11017109 | 23404247 | 9371269 | 9537421 |
| 11988839 | 22031933 | 19419954 | 20573930 | 18768914 |
| 25778702 | NA | 11751408 | 22334892 | 25110901 |
| 15574327 | 26771495 | 25115383 | 15607759 | 29298899 |
| 10203277 | 26246577 | 24535670 | 9380738 | 26725424 |
| 10097108 | 23573288 | 25984556 | 12587805 | 27809445 |
| 27257257 | 14673115 | 26139350 | 12727798 | 24404629 |
| 20932475 | 19625394 | 18761324 | 10963623 | 14673705 |
| 17488475 | 24478428 | 11729303 | 16280036 | 12522687 |
| 12024041 | 16051304 | 1597188 | 15535854 | 30307446 |
| 1055055 | 15103332 | 10622721 | 12714703 | 12910492 |
| 15122335 | 23555248 | 6182444 | 17380162 | 27457486 |
| 20019063 | 26487564 | 23746447 | 28431792 | 29057873 |
| 20841568 | 23342373 | 22902835 | 23519123 | 22474067 |
| 7671312 | 26861015 | 30030361 | 11040209 | 28098593 |
| 11076961 | 17005688 | 18682835 | 9585500 | 25328986 |
| 7983002 | 21866103 | 15867910 | 25429310 | 32557261 |
| 17428792 | 17339424 | 11602624 | 23619365 | 21811631 |
| 10672017 | 22383882 | 18220859 | 22327366 | 20657596 |
| 18174154 | 10975523 | 18374667 | 23085511 | 22281838 |
| 9660939 | 26468524 | 17659995 | 23376832 | 26561703 |
| 16122426 | 18089854 | 23977373 | 19996457 | 12655414 |
| 15468306 | 16446357 | 23592794 | 17307989 | 28404813 |
| 28356513 | 16477006 | 23154981 | 20603081 | 25418138 |
| 23565119 | 18052967 | 22960178 | 19555663 | 27995415 |
| 20713514 | 22007134 | 21741376 | 11967526 | 33685683 |

| | | | | |
|---|---|---|---|---|
| 1406652 | 12724731 | 11044454 | 16951403 | 2992548 |
| 19841733 | 15548681 | 8440384 | 9683613 | 18203897 |
| 24369348 | 9849491 | 19951991 | 9099683 | 10720158 |
| 31035700 | 18498133 | 16858407 | 15940673 | 15189125 |
| 30871156 | 2656050 | 228272 | 7577714 | 11307174 |
| 21633178 | 16709241 | 6371429 | 1848582 | 18420277 |
| 21636835 | 15638735 | 9767110 | 11519011 | 17283068 |
| 10914538 | 19219653 | 8652655 | 15851485 | 2167130 |
| 18803879 | 18820913 | 190267 | 4774123 | 10471130 |
| 2899130 | 18267032 | 19747065 | 15536089 | 10833329 |
| 3790257 | 11956089 | 18583509 | 19740703 | 16547353 |
| 12856180 | 17549067 | 18522490 | 10048303 | 17046972 |
| 20354580 | 19093176 | 7649494 | 2870496 | 17389926 |
| 12479567 | 19302291 | 7278682 | 1652755 | 10100195 |
| 21504868 | 16510598 | 18198219 | 1708392 | 11839807 |
| 27477280 | 8996164 | 15668660 | 14726604 | 10216279 |
| 9654093 | 17283124 | 11325678 | 19628033 | 17508906 |
| 16424607 | 18992248 | 19854035 | 8846925 | 14976195 |
| 29352142 | 18390668 | 7639730 | 14642406 | 17912575 |
| 30042655 | 14555507 | 3918580 | 11878747 | 10101267 |
| 31101865 | 18837291 | 12432931 | 18424738 | 15704532 |
| 30110365 | 18075467 | 17439363 | 19024248 | 9442035 |
| 30925886 | 18607850 | 6884990 | 8325534 | 8769129 |
| 33268902 | 19144510 | 11447214 | 19625220 | 17916561 |
| 33043019 | 19402749 | 19706381 | 10980461 | 15063746 |
| 32760721 | 15814641 | 9325339 | 18938145 | 2181276 |
| 32015325 | 15901346 | 932041 | 10802064 | 15317910 |
| 32256352 | 16785472 | 18428149 | 14749265 | 12775843 |
| 30921410 | 18035049 | 10544287 | 17463087 | 9112289 |
| 31740582 | 17716232 | 19375431 | 18680158 | 11964182 |
| 33495651 | 17695509 | 6814427 | 8986768 | 1542667 |
| 30808384 | 22100631 | 8567691 | 6300970 | 3013986 |
| 27987249 | 11972054 | 16887964 | 11121721 | 16047261 |
| 9831565 | 18987785 | 18656701 | 17583675 | 11761328 |
| 12899623 | 16413410 | 15925209 | 10653827 | 11956665 |
| 15173886 | 9075838 | 14677018 | 10914032 | 18479189 |
| 11891120 | 11939906 | 10569628 | 9873062 | 18409172 |
| 20644576 | 8534261 | 14704851 | 15585321 | 15026176 |
| 26353940 | 19403638 | 12895592 | 1618328 | 19013211 |
| 9851930 | 6418841 | 18598592 | 15680219 | 17016550 |
| 20639876 | 6122208 | 10842581 | 15780594 | 12569109 |
| 17339406 | 19660687 | 16698314 | 2448410 | 19014349 |
| 19252480 | 9584338 | 16430221 | 10666321 | 16962588 |
| 32182061 | 12432932 | 11264458 | 18562168 | 19822456 |
| 16267625 | 9085163 | 9737710 | 11215511 | 17526492 |
| 18357466 | 18032380 | 10191291 | 11460480 | 8877730 |
| 19339911 | 11090610 | 12627223 | 10705969 | 15817466 |
| 19077464 | 17081103 | 218223 | 16011463 | 374662 |
| 18172246 | 16574427 | 9627909 | 230492 | 8617728 |
| 19383847 | 16174820 | 15215856 | 16120612 | 18927507 |
| 1272473 | 15689384 | 2602371 | 3790723 | 9587024 |

| | | | | |
|---|---|---|---|---|
| 10064852 | 7276750 | 9462665 | 27889578 | 17895972 |
| 17713401 | 12231557 | 11412116 | 14570712 | 23576886 |
| 11017945 | 12062442 | 19415921 | 18332424 | 18215622 |
| 8386626 | 12831960 | 9242711 | 14585353 | 28334053 |
| 19674315 | 17392543 | 19413181 | 12368261 | 28859574 |
| 18354163 | 15314690 | 2246616 | 22267161 | 19436069 |
| 4355366 | 19268692 | 11001804 | 11237210 | 26287747 |
| 18498226 | 9370338 | 18164739 | 26670047 | 26287746 |
| 19464347 | 11716958 | 17927969 | 30447757 | 12040175 |
| 1730245 | 16339544 | 14993240 | 21167873 | 12164919 |
| 18718914 | 17296604 | 11533328 | 17298186 | 34244591 |
| 10880336 | 16219795 | 12815058 | 22483044 | 16435184 |
| 19403631 | 3919061 | 17534535 | 18184796 | 3126356 |
| 15670717 | 4387676 | 874074 | 17644144 | 16483879 |
| 17511631 | 7288293 | 9368067 | 16055064 | 24168112 |
| 9570154 | 19769461 | 9852097 | 24727796 | 15471987 |
| 1848655 | 3099851 | 3046314 | 24346713 | 17263796 |
| 458381 | 2497518 | 16716149 | 10473536 | 27041232 |
| 11251339 | 17049925 | 9789062 | 10613508 | 25405608 |
| 698243 | 6311078 | 17477829 | 20188650 | 17384686 |
| 8386433 | 17198385 | 19521349 | 20457613 | 1782695 |
| 17612623 | 19803417 | 10716626 | 14676271 | 19272414 |
| 14580199 | 19167960 | 16835236 | 19968948 | 27127795 |
| 5637427 | 10448523 | 15680232 | 24966374 | 25690510 |
| 17311055 | 2174886 | 2175712 | 25113167 | 24726665 |
| 10484769 | 15104204 | 21996254 | 23202739 | 21226711 |
| 16251722 | 9133619 | 20534694 | 16908410 | 21345369 |
| 2610352 | 2876046 | 12440980 | 18474609 | 20183829 |
| 11929995 | 8301225 | 14500912 | 24533017 | 23897649 |
| 16202558 | 9730949 | 16459311 | 12563289 | 28203483 |
| 15837518 | 14999406 | 12522256 | 11837891 | 17076662 |
| 4086942 | 4690137 | 3091479 | 30402088 | 9928160 |
| 15016776 | 12679481 | 16648838 | 27257628 | 17637475 |
| 6295699 | 10444342 | 26215792 | 27766264 | 15885916 |
| 19825219 | 11861418 | 29456534 | 17496911 | 10980593 |
| 17365173 | 6113262 | 30076771 | 9839441 | 24568968 |
| 6758698 | 16319061 | 15467723 | 28473628 | 27410235 |
| 9278416 | 16990795 | 33178221 | 27573877 | 27592409 |
| 18524888 | 17276401 | 31986264 | 25781681 | 22076657 |
| 8656081 | 8968737 | 32513989 | 15673435 | 27359210 |
| 8413223 | 8806705 | 20410124 | 23641197 | 17143559 |
| 7429333 | 15976321 | 30705620 | 30686771 | 22308501 |
| 19557870 | 18757362 | 20505079 | 27847432 | 29183756 |
| 18541719 | 1377680 | 17084704 | 22017584 | 25152404 |
| 19619139 | 4386961 | 24044036 | 19345194 | 15934937 |
| 9501215 | 12209373 | 30364226 | 26468183 | 25017118 |
| 11413152 | 19450180 | 22510460 | 22405502 | 18272487 |
| 12432933 | 18617006 | 17088211 | 27239350 | 16819983 |
| 16911361 | 12730456 | 19602257 | 21162125 | 28196718 |
| 17960332 | 18394665 | 28183735 | 15219735 | 24602615 |
| 18209571 | 9535873 | 29410531 | 25729352 | 15885774 |

| | | | | |
|---|---|---|---|---|
| 19276245 | 22761566 | 23775697 | 12919045 | 18854196 |
| 25449850 | 19606501 | 9131163 | 7743460 | 7951064 |
| 33136286 | 12106454 | 192727 | 9299423 | 10692424 |
| 11445798 | 21953180 | 3753461 | 22877991 | 15172163 |
| 4152527 | 21278789 | 9526094 | 2163656 | 22809994 |
| 8127060 | 21078976 | 9370318 | 2153142 | 23022039 |
| 10515893 | 21934092 | 8063717 | 7768880 | 6088233 |
| 29483667 | 19035562 | 9370322 | 6257301 | 6145585 |
| 8099811 | 19244115 | 9838049 | 7574485 | 18791037 |
| 4152248 | 14988435 | 1718687 | 8037660 | 17917066 |
| 18515354 | 25415055 | 19250975 | 9561269 | 2988417 |
| 2294991 | 18425414 | 1311951 | 21554199 | 9131118 |
| 238530 | 1323041 | 21979151 | 1323040 | 11286640 |
| 15717202 | 11740936 | 9654085 | 9370334 | 8777586 |
| 16618936 | 8382477 | 8405660 | 10882337 | 15979148 |
| 7937585 | 9370317 | 2364071 | 8869881 | 330281 |
| 5289242 | 17095752 | 6712967 | 1362247 | 1897971 |
| 8810901 | 11171073 | 993673 | 21282087 | 476064 |
| 6113726 | 19401146 | 8883840 | 20559679 | 23240538 |
| 21942574 | 19362164 | 1628249 | 20503434 | 22750097 |
| 1711780 | 16859663 | 10208837 | 8499439 | 9370331 |
| 23727593 | 5581578 | 5580660 | 11324699 | 1323036 |
| 7680650 | 18204095 | 7378426 | 17412762 | 9553082 |
| 7215948 | 8866672 | 8274015 | 1128165 | 1495422 |
| 11893509 | 9386267 | 1495442 | 20178759 | 10396601 |
| 6995243 | 1989575 | 10893425 | 10601694 | 16339116 |
| 22539939 | 7107629 | 6354980 | 23054682 | 1495441 |
| 24584707 | 9370329 | 1546171 | 20167241 | 9370315 |
| 19112489 | 22136116 | 2498871 | 1329260 | 9144082 |
| 29357412 | 17673461 | 8200054 | 2499328 | 23010477 |
| 32030122 | 8468530 | 23143232 | 18955040 | 17171187 |
| 20960469 | 2505652 | 16899548 | 22319379 | 17558022 |
| 32715478 | 5277094 | 15821158 | 9370335 | 7215553 |
| 8402000 | 12824553 | 17318530 | 2663077 | 17940275 |
| 12833157 | 17294083 | 8699417 | 9468529 | 22215515 |
| 18087673 | 677084 | 3071807 | 19413994 | 17079146 |
| 15208781 | 8440692 | 1819750 | 9370312 | 2832400 |
| 16175503 | 1333282 | 22981911 | 8993541 | 8391253 |
| 17661906 | 7961445 | 11330037 | 6263342 | 19318427 |
| 15485997 | 22285183 | 7537411 | 22892679 | 15481814 |
| 2446635 | 23775696 | 23611148 | 9370321 | 9458812 |
| 15821736 | 14759225 | 8391435 | 10729607 | 205412 |
| 15790351 | 23712958 | 18525025 | 6251897 | 12743757 |
| 25167330 | 2545264 | 2143679 | 17881348 | 6113006 |
| 15731287 | 18204094 | 9765874 | 9370313 | 17132865 |
| 15986374 | 12749687 | 1550861 | 20176101 | 19324408 |
| 17010589 | 22014644 | 21214572 | 8382961 | 16015482 |
| 20378641 | 17157506 | 22374091 | 2118266 | 18341203 |
| 29459677 | 236033 | 6388643 | 10761925 | 23394527 |
| 21210154 | 2083674 | 16495223 | 10358924 | 1932343 |
| 29030052 | 9370323 | 1717219 | 8765148 | 9547571 |

| | | | | |
|---|---|---|---|---|
| 21077828 | 15361348 | 23945590 | 16524428 | 29577047 |
| 19539604 | 19422785 | 12943534 | 20201954 | 12435631 |
| 19540930 | 10905630 | 27217160 | 19717472 | 10949293 |
| 3440872 | 23295697 | 29666272 | 27428965 | 15145825 |
| 12038971 | 8709678 | 7635144 | 28632878 | 11805083 |
| 2096696 | 6297743 | 23871722 | 7757071 | 11839797 |
| 4092051 | 2833930 | 17986282 | 31949887 | 24991833 |
| 19666474 | 9370319 | 8016100 | 2573431 | 32205204 |
| 6288375 | 22468920 | 25270028 | 20381640 | 30406125 |
| 23350810 | 18621144 | 10581255 | 28911859 | 17667842 |
| 14626658 | 9370326 | 16100110 | 32276449 | 18852458 |
| 9590628 | 8786821 | 6366476 | 29761529 | 19893639 |
| 639820 | 15878874 | 21334936 | 18828673 | 21953136 |
| 10396999 | 21303393 | 11058895 | 15695812 | 23008150 |
| 18508126 | 16719778 | 10944123 | 24051395 | 26253919 |
| 19450542 | 4005284 | 19576565 | 23631851 | 23943603 |
| 4369816 | 1323034 | 11733556 | 3103658 | 24011563 |
| 21703569 | 22960354 | 11901181 | 27789274 | 31863285 |
| 22103853 | 3707974 | 12872255 | 24942883 | 24458711 |
| 15922587 | 9370330 | 15945070 | 23133399 | 17503467 |
| 10191259 | 5334818 | 11228641 | 23153495 | 25883951 |
| 16717392 | 3037250 | 18452889 | 22399345 | 21835778 |
| 19657568 | 10759022 | 17166182 | 18593557 | 16485879 |
| 8488570 | 22910056 | 24685145 | 17376426 | 15833859 |
| 1495418 | 16246718 | 18455129 | 23847349 | 15573099 |
| 2936199 | 10395968 | 10980531 | 31695792 | 21163869 |
| 23765576 | 19328771 | 8808595 | 19393343 | 22710719 |
| 3707712 | 21818839 | 14973782 | 19557177 | 23940611 |
| 880944 | 2543404 | 28816422 | 17350578 | 20725040 |
| 14604010 | 20651826 | 29459785 | 20691899 | 25703554 |
| 6268930 | 20153716 | 29297247 | 16990131 | 22872737 |
| 17182612 | 6705975 | 12480927 | 19620969 | 25241263 |
| 1323038 | 18295604 | 5288798 | 16914500 | 10901230 |
| 22609101 | 479108 | 10788335 | 19578358 | 9483513 |
| 7031820 | 10427554 | 14973778 | 27234298 | 9378992 |
| 10215861 | 8382767 | 20080937 | 22617791 | 19028820 |
| 15893598 | 2160964 | 12217961 | 24766805 | 11929849 |
| 23220394 | 7492326 | 9525984 | 20549139 | 8156598 |
| 6264966 | 12573444 | 29874875 | 21139080 | 19008118 |
| 8389973 | 19889969 | 10835346 | 1451437 | 10673356 |
| 9370314 | 17063928 | 22304930 | 28042609 | 19324969 |
| 3818635 | 22960381 | 20637498 | 27148389 | 10937998 |
| 17012796 | 17069352 | 15148656 | 22128289 | 26830228 |
| 2833508 | 5083874 | 12887896 | 23633457 | 16858403 |
| 9083101 | 22628558 | 17273964 | 25564569 | 12453919 |
| 16838328 | 27304503 | 10527672 | 25018647 | 15317753 |
| 15134747 | 11687968 | 15741281 | 21355094 | 9233789 |
| 17928411 | 21336310 | 11836223 | 19490906 | 15165456 |
| 17646670 | 27591812 | 9399852 | 22088887 | 19342379 |
| 3300655 | 29362483 | 30705290 | 17304241 | 10959075 |
| 15273989 | 31920721 | 27720922 | 29565815 | 8175923 |

| | | | | |
|---|---|---|---|---|
| 23016862 | 1280824 | 2227617 | 28826372 | 23399566 |
| 12006103 | 9196040 | 15065877 | 8839927 | 9054771 |
| 19135507 | 24696235 | 19528880 | 29882869 | 3283935 |
| 25451943 | 7988557 | 18022819 | 19689262 | 27013343 |
| 28844715 | 7665574 | 8681959 | 32633718 | 28930145 |
| 24243012 | 9374471 | 12949720 | 29354093 | 29163511 |
| 1890850 | 10514501 | 19840950 | 25748677 | 22824096 |
| 23400783 | 17968323 | 3218790 | 11831846 | 26926090 |
| 21359530 | 8232552 | 16274220 | 8016083 | 25254104 |
| 20086245 | 8181059 | 23793029 | 24363178 | 25326323 |
| 16125000 | 11574262 | 30843452 | 1653609 | 18755000 |
| 19461653 | 11574261 | 18950740 | 31790802 | 8634065 |
| 30578699 | 17244752 | 22002721 | 31954874 | 12446022 |
| 25311867 | 11242034 | 26305592 | 31676443 | 16132226 |
| 23162553 | 19302040 | 18694559 | 31812582 | 16888915 |
| 27622013 | 17442919 | 20512146 | 28118532 | 9831245 |
| 28477742 | 24550720 | 18376416 | 31616242 | 9286353 |
| 30504141 | 22307082 | 21266464 | 30980044 | 15491976 |
| 29660367 | 19229310 | 11980706 | 31736978 | 16047947 |
| 21654544 | 22465036 | 23150559 | 20811799 | 12115225 |
| 12355441 | 30065109 | 28800946 | 26186194 | 23380452 |
| 33363734 | 12893990 | 23532844 | 24438557 | 28527011 |
| 18803764 | 14646693 | 22179047 | 32111819 | 16838012 |
| 31134055 | 12730278 | 22152675 | 31819986 | 15817569 |
| 31088566 | 15608127 | 15224133 | 28888937 | 30054974 |
| 11698225 | 11990381 | 22863007 | 26165754 | 17267393 |
| 24780758 | 11602529 | 18772192 | 24627487 | 26447148 |
| 20536554 | 12810652 | 26982032 | 29945215 | 27105113 |
| 18794151 | 10815927 | 18413257 | 31602316 | 19851446 |
| 20064376 | 12960109 | 16682973 | 19430479 | 29925282 |
| 9872990 | 10197614 | 22121117 | 10799542 | 21525168 |
| 1496401 | 12181437 | 21422230 | 16497175 | 16600869 |
| 8628273 | 9466980 | 12403812 | 24239284 | 25762440 |
| 9857039 | 10220571 | 20375344 | 29597279 | 22922464 |
| 7608146 | 9950149 | 10764818 | 32469225 | 10100484 |
| 1638633 | 21566147 | 16760425 | 33664446 | 16278232 |
| 9724754 | 9242408 | 21725307 | 32159237 | 14615802 |
| 9326223 | 21036394 | 15107855 | 33377319 | 29286006 |
| 9295335 | 18794901 | 18371931 | 30654597 | 21800163 |
| 8496154 | 20685892 | 10806194 | 33571544 | 21936831 |
| 8524272 | 21543532 | 8798640 | 32155444 | 19509293 |
| 9287210 | 16227996 | 19625297 | 32422320 | 28870987 |
| 8780698 | 21346250 | 14623329 | 26271607 | 29072575 |
| 1468582 | 21252237 | 29186351 | 1608291 | 31867277 |
| 9305869 | 19923220 | 26411921 | 33505321 | 23395095 |
| 8550573 | 15755449 | 32273485 | 32130973 | 28327630 |
| 17604604 | 21699959 | 21664428 | 33170317 | 29371938 |
| 10022928 | 19682329 | 17054399 | 34281182 | 30073421 |
| 2153461 | 12547239 | 25985275 | 29167338 | 32768523 |
| 10428508 | 5075227 | 23596439 | 29256392 | 33329574 |
| 10644731 | 19381358 | 33454021 | 27180971 | 33462414 |

| | | | | |
|---|---|---|---|---|
| 24780002 | 17112607 | 21423409 | 4061122 | 20303741 |
| 24607545 | 15781663 | 17573714 | 190272 | 23898905 |
| 34073720 | 10646883 | 25458568 | 6961921 | 28910500 |
| 12612292 | 21924373 | 23960241 | 12016260 | 23583456 |
| 27300434 | 28323937 | 21070191 | 18499582 | 27562463 |
| 27507853 | 9727023 | 20368621 | 24808179 | 25277212 |
| 30101371 | 10951588 | 20697302 | 15728179 | 29251630 |
| 23550303 | 3283656 | 28522374 | 26965621 | 23622250 |
| 9378538 | 25323927 | 23611944 | 19139765 | 17468766 |
| 32317288 | 10468914 | 20400852 | 3630977 | 10086361 |
| 12589646 | 29433126 | 22334035 | 17215125 | 20407820 |
| 31285550 | 16785999 | 20010955 | 25108285 | 9253712 |
| 27282309 | 18668205 | 16251272 | 12730697 | 9488659 |
| 28614305 | 15982921 | 19554514 | 11717312 | 19018142 |
| 26078352 | 3283542 | 22817889 | 18039658 | 11055975 |
| 33800494 | 29610148 | 26091043 | 23475612 | 18342376 |
| 34226685 | 17540175 | 22453014 | 22236406 | 10189350 |
| 28740119 | 3491291 | 19718025 | 27068984 | 8621626 |
| 32882916 | 6308607 | 22842228 | 26224785 | 10864911 |
| 23975423 | 26878173 | 22622578 | 6805319 | 10615383 |
| 21484256 | 12670889 | 21946352 | 168823 | 9568714 |
| 31283845 | 12866375 | 30987166 | 3477815 | 10827130 |
| 30642555 | 12114746 | 25059483 | 24288038 | 11101507 |
| 26358421 | 3043188 | 24732412 | 23066022 | 25903473 |
| 15686623 | 11278720 | 25603176 | 7730305 | 21205967 |
| 19753302 | 11278702 | 25738837 | 10672230 | 27568792 |
| 12771921 | 17726008 | 12618595 | 8187868 | 17488487 |
| 33232793 | 22539006 | 16399907 | 13018271 | 22193159 |
| 16216411 | 10970876 | 15616152 | 3052428 | 20031384 |
| 27915330 | 30210299 | 24628039 | 949837 | 26655797 |
| 24758178 | 11050117 | 22225631 | 15234968 | 25863248 |
| 19424592 | 10756053 | 27441728 | 7961626 | 27036018 |
| 14668814 | 25011106 | 21331042 | 12234803 | 18695042 |
| 17880691 | 22721863 | 22545159 | 7730304 | 19826040 |
| 16231422 | 15308747 | 15986483 | 8506365 | 19141645 |
| 17377501 | 19157421 | 10617567 | 7060582 | 20029046 |
| 21991364 | 27352031 | 19561075 | 9765290 | 19202062 |
| 22917536 | 10490655 | 11260262 | 3724458 | 15944709 |
| 15689376 | 24069510 | 21266536 | 17558466 | 19833767 |
| 9513715 | 16613900 | 7426196 | 10781873 | 18066065 |
| 23840669 | 18820302 | 3004475 | 19696787 | 18521080 |
| 23154416 | 18203756 | 2511019 | 20371350 | 20351064 |
| 18562482 | 23431031 | 8939939 | 8393662 | 19258499 |
| 12907752 | 24662006 | 21480869 | 16435220 | 20406979 |
| 11146549 | 24615633 | 6403642 | 6822523 | 17135268 |
| 23404106 | 20716963 | 7327552 | 16807786 | 17656095 |
| 16678816 | 21212100 | 16452169 | 26536169 | 17823410 |
| 18413260 | 23351786 | 194920 | 19496715 | 19221490 |
| 18681747 | 20452318 | 3944267 | 24812413 | 18538733 |
| 17540168 | 22615490 | 15071125 | 11114738 | 20018759 |
| 14602780 | 22141737 | 28538136 | 22042966 | 19597470 |

| | | | | |
|---|---|---|---|---|
| 20404092 | 32464637 | 23238060 | 28438858 | 28086984 |
| 17135249 | 32839770 | 24622826 | 30084000 | 19166931 |
| 19377482 | 19692591 | 17086380 | 26116534 | 18466115 |
| 19034270 | 33019591 | 23176034 | 18469807 | 26678875 |
| 20080624 | 23321557 | 21778025 | 21460185 | 20505359 |
| 19273599 | 32574107 | 28003096 | 18253061 | 29409688 |
| 18949056 | 28956771 | 12163693 | 27856613 | 22221393 |
| 18701644 | 33937724 | 20446114 | 24528886 | 27416781 |
| 19536137 | 21703540 | 24959120 | 26039999 | 30619462 |
| 21177881 | 33015593 | 15693941 | 18719102 | 20083114 |
| 16111679 | 33116300 | 11701706 | 24472220 | 20356743 |
| 18579786 | 31115493 | 11854458 | 12062040 | 29233870 |
| 20212154 | 32995797 | 26785480 | 19211685 | 27412492 |
| 15077171 | 17451827 | 18429699 | 27882935 | 25012593 |
| 17942906 | 32818486 | 22971926 | 12598900 | 29158945 |
| 17784791 | 21187859 | 26890602 | 20693423 | 23630460 |
| 17569667 | 32728199 | 21646373 | 21238923 | 23087426 |
| 19120703 | 32979938 | 26686024 | 25288737 | 21779718 |
| 27903835 | 32726803 | 30778219 | 15729359 | 17440456 |
| 25403569 | 32699849 | 26009982 | 27034160 | 29898976 |
| 27720676 | 21597473 | 19052657 | 10366627 | 33510058 |
| 27623250 | 30449619 | 20725615 | 11387206 | 23853735 |
| 17383918 | 15504727 | 16628003 | 18216145 | 33511116 |
| 20816094 | 16261263 | 19470756 | 32555321 | 7615345 |
| 17557076 | 15103690 | 23751779 | 33422265 | 27187935 |
| 27226634 | 14645660 | 25770769 | 32691695 | 15291819 |
| 24564666 | 16803896 | 22209643 | 32376634 | 25598354 |
| 24838397 | 15155839 | 19643170 | 32307550 | 30459650 |
| 18179882 | 12805287 | 19206551 | 32492406 | 22287959 |
| 24089531 | 16452680 | 22064246 | 32733001 | 32932623 |
| 9837812 | 18614672 | 22107733 | 32838362 | 24742457 |
| 27626371 | 18317590 | 20398356 | 32228226 | 33239064 |
| 25678554 | 17941718 | 22720979 | 33845483 | 19909229 |
| 17344420 | 17005849 | 23428231 | 32048163 | 20842175 |
| 22982022 | 12629174 | 19516051 | 32619549 | 33061891 |
| 22356826 | 15963952 | 21593791 | 32132184 | 25078115 |
| 24344204 | 15774771 | 22098780 | 32511476 | 27443914 |
| 21057504 | 19179283 | 21974862 | 30190671 | 11403878 |
| 27215383 | 12196911 | 21979174 | 21108791 | 29018191 |
| 19463981 | 15728187 | 28438689 | 19575671 | 33050345 |
| 16200211 | 14566944 | 29991871 | 16402914 | 28603493 |
| 18614015 | 17032905 | 18391788 | 20083100 | 19903816 |
| 20858599 | 14697674 | 17229764 | 21784065 | 11908751 |
| 20818383 | 16452991 | 23520208 | 15731351 | 12827358 |
| 19752196 | 12361670 | 25667580 | 16845469 | 15734728 |
| 23260140 | 17188889 | 19273616 | 12815623 | 19184645 |
| 16218961 | 16055563 | 22231519 | 14973296 | 21784059 |
| 19028688 | 17067279 | 21172653 | 15944192 | 28167679 |
| 20226757 | 18674600 | 29375318 | 28561066 | 8074175 |
| 18391175 | 18445122 | 25483983 | 21628530 | 32604946 |
| 33106987 | 19494120 | 19543238 | 20542007 | 17186029 |

| | | | | |
|---|---|---|---|---|
| 18161746 | 31495888 | 24382962 | 30862117 | 30578919 |
| 17110941 | 10854423 | 22652455 | 25823658 | 25092323 |
| 16823444 | 12034848 | 30207631 | 24459210 | 12077124 |
| 25501551 | 26627236 | 11299310 | 18311132 | 11013305 |
| 22705006 | 20368362 | 26170736 | 16403804 | 9660774 |
| 30315846 | 16473935 | 26064108 | 29693488 | 32696532 |
| 28892558 | 18079701 | 11549701 | 21964294 | 24288332 |
| 19352614 | 32358495 | 24068427 | 23644458 | 22465940 |
| 12891546 | 22492724 | 18599270 | 15328537 | 22366074 |
| 16364488 | 15210332 | 31091547 | 19587107 | 2335522 |
| 28983452 | 33010169 | 27237973 | 18847334 | 10748047 |
| 22570745 | 23906714 | 10884055 | 19917303 | 23273843 |
| 18242193 | 16751180 | 33613258 | 30936491 | 19805370 |
| 20303879 | 22453236 | 14764632 | 15864272 | 17872378 |
| 32584474 | 15882621 | 29413903 | 20016026 | 21796212 |
| 28341729 | 26865925 | 18198212 | 23858833 | 8790411 |
| 32685191 | 17643379 | 26220343 | 17607736 | 10588945 |
| 27117316 | 29069470 | 24333629 | 23882024 | 2384150 |
| 26950144 | 21855803 | 22550093 | 16456071 | 25104388 |
| 17328863 | 19029798 | 31667577 | 24724793 | 17873880 |
| 28572459 | 29955842 | 28667055 | 23224777 | 30471425 |
| 20418096 | 32235701 | 23349191 | 21353301 | 24735479 |
| 23322901 | 23901102 | 12673048 | 20713077 | 25271621 |
| 25871831 | 20887785 | 11435939 | 20628458 | 24491228 |
| 9245493 | 17872503 | 28487995 | 21793770 | 9013544 |
| 26485378 | 15610069 | 23284041 | 23322373 | 31009661 |
| 22819548 | 15907797 | 27667570 | 16697548 | 12569201 |
| 23434765 | 11038252 | 12748652 | 25739888 | 27647924 |
| 14570043 | 16459330 | 16012053 | 23865044 | 33246156 |
| 15923610 | 18505913 | 27543160 | 32700336 | 19996111 |
| 21189866 | 16547273 | 30992313 | 7654171 | 28923174 |
| 19321346 | 9388184 | 34501340 | 3881803 | 7925343 |
| 20104017 | 18430728 | 34786213 | 10600673 | 18621681 |
| 28618167 | 15012602 | 32403258 | 388618 | 10602018 |
| 27542226 | 21548952 | 10675335 | 15869601 | 19502589 |
| 26795388 | 12911774 | 17446932 | 11771674 | 11673457 |
| 22948112 | 11207308 | 20716577 | 4381909 | 28322867 |
| 34039996 | 17471233 | 16115815 | 11340051 | 29960034 |
| 31572065 | 10356359 | 17227891 | 15490415 | 9717719 |
| 26209534 | 12947119 | 16478798 | 4291593 | 21796211 |
| 17631638 | 18927239 | 11285237 | 14641005 | 16685654 |
| 19909241 | 8019414 | 11792809 | 10477257 | 10097111 |
| 30216699 | 12466265 | 12370805 | 21576599 | 10067858 |
| 17974973 | 10762064 | 21828285 | 23759795 | 10487205 |
| 33188728 | 9242908 | 14755334 | 23415802 | 10207619 |
| 16729019 | 17389618 | 22103516 | 11089551 | 10557099 |
| 21452186 | 22634633 | 18371421 | 10655068 | 11163208 |
| 23045548 | 13271447 | 27865926 | 16385454 | 10498912 |
| 21676658 | 8752328 | 10984438 | 9614081 | 10527909 |
| 22118460 | 11158046 | 24123709 | 10377398 | 32404993 |
| 11972036 | 24515297 | 12960086 | 17428920 | 29099489 |

| | | | | |
|---|---|---|---|---|
| 33932560 | 9367159 | 11553680 | 9971736 | 23817417 |
| 29436617 | 10716940 | 21161717 | 27941249 | 20400695 |
| 12759238 | 15902656 | 21283639 | 28074012 | 12010778 |
| 15952880 | 10802647 | 12666113 | 8019699 | 29247122 |
| 15503154 | 16554828 | 16295699 | 29425059 | 32398875 |
| 16022590 | 29056325 | 19061375 | 27288456 | 26505736 |
| 12716939 | 31035587 | 17567803 | 12919684 | 30381825 |
| 12783850 | 18215151 | 20067580 | 19043080 | 19273625 |
| 17911161 | 26516076 | 19342987 | 19536196 | 32192578 |
| 19956200 | 26705830 | 11585624 | 25090446 | 15978547 |
| 19524505 | 20811577 | 19770513 | 23934149 | 32407669 |
| 12628165 | 10952417 | 21168495 | 16484492 | 28583370 |
| 15229644 | 11714087 | 21092735 | 20870751 | 16785567 |
| 18426756 | 10528039 | 16498630 | 21839365 | 21595737 |
| 12728264 | 24759575 | 21657851 | 20130528 | 26982353 |
| 19088304 | 25660022 | 21868473 | 23502960 | 21715307 |
| 15896322 | 11357143 | 16623828 | 20537521 | 29935220 |
| 18680555 | 15784165 | 23152062 | 21125407 | 25229003 |
| 15651302 | 17360849 | 22399755 | 22958489 | 9792917 |
| 15054086 | 18842113 | 17629414 | 22334613 | 22014308 |
| 15505146 | 9512493 | 23035104 | 31796734 | 17277314 |
| 23383003 | 21620960 | 22536415 | 18796515 | 12835412 |
| 22432004 | 16847462 | 16504406 | 26993153 | 11087758 |
| 15040260 | 15209374 | 20547125 | 27437668 | 18508192 |
| 23685749 | 12531180 | 19350383 | 27871366 | 23568971 |
| 12618436 | 12660731 | 22884909 | 32153505 | 20943776 |
| 7565688 | 21779440 | 17917587 | 25620207 | 15375155 |
| 12368238 | 12048182 | 23045704 | 25727005 | 11672424 |
| 27372738 | 21440011 | 20202079 | 27320729 | 10785607 |
| 2841963 | 9838078 | 21771809 | 29381406 | 22307589 |
| 21769672 | 16187290 | 11059815 | 32439107 | 11017125 |
| 2557116 | 15505410 | 21640756 | 7796919 | 16492677 |
| 22512244 | 17496910 | 23027386 | 33324646 | 31286677 |
| 2545232 | 19363522 | 19932745 | 29587204 | 1978318 |
| 1661577 | 28554312 | 22564823 | 32549377 | 25105124 |
| 1755382 | 29348254 | 19770517 | 15616590 | 12446672 |
| 2546599 | 21673720 | 17482149 | 33811820 | 12716889 |
| 20657170 | 10094148 | 9108119 | 28803844 | 26308067 |
| 30124109 | 22930444 | 18930956 | 21191810 | 26327396 |
| 27660392 | 12861073 | 15649943 | 30930090 | 26582078 |
| 29793167 | 22771695 | 22694955 | 31687975 | 25126564 |
| 28467092 | 9872457 | 29499229 | 20388082 | 11514575 |
| 29778464 | 17698296 | 9153397 | 28298525 | 11788602 |
| 9714779 | 16602711 | 27931246 | 15367596 | 18695242 |
| 9416770 | 20304963 | 25002992 | 23575689 | 18316376 |
| 26590421 | 17706365 | 21914775 | 22590492 | 26586472 |
| 27037577 | 18267965 | 23341784 | 16446378 | 9843703 |
| 18474859 | 20484625 | 25957321 | 21681369 | 16883319 |
| 4331040 | 21373950 | 8863824 | 20018632 | 15322102 |
| 8640223 | 21957482 | 28009282 | 24285838 | 24311738 |
| 16365287 | 18637713 | 22065085 | 20970516 | 19184091 |

| | | | | |
|---|---|---|---|---|
| 15383652 | 22442671 | 15153069 | 2433278 | 15879121 |
| 21170887 | 9489617 | 15699391 | 15609003 | 15572664 |
| 23908456 | 18086946 | 17242337 | 25406093 | 10625698 |
| 17164835 | 32526773 | 20206331 | 15536068 | 11698415 |
| 28634454 | 33065209 | 17916383 | 19005037 | 10861044 |
| 17938243 | 32562843 | 16400613 | 15546853 | 10629034 |
| 17803961 | 33375371 | 18632736 | 22260796 | 11297525 |
| 18834332 | 10320667 | 18319072 | 2459516 | 1447167 |
| 16122728 | 1336455 | 24264046 | 16325348 | 9639556 |
| 16386727 | 10712923 | 25498144 | 21959103 | 15604236 |
| 9603781 | 27477490 | 19270536 | 20383137 | 12746452 |
| 17222338 | 26398287 | 11729304 | 20826676 | 9159118 |
| 17849436 | 19536897 | 33082294 | 8190100 | 12600988 |
| 16201968 | 19584267 | 9529250 | 11257115 | 12239223 |
| 18519637 | 22155636 | 33395426 | 10196191 | 15492006 |
| 17403901 | 29543893 | 33458558 | 12488455 | 10711350 |
| 10938085 | 23729401 | 33442700 | 11500364 | 11108712 |
| 30927255 | 15947886 | 32869019 | 12048245 | 11304531 |
| 2615857 | 19740705 | 32125455 | 11782488 | 14965240 |
| 15078215 | 19850741 | 32227760 | 15814901 | 9272731 |
| 27078878 | 15469821 | 23943763 | 12730329 | 15795236 |
| 12270951 | 7565706 | 32662421 | 15572660 | 11496823 |
| 3479638 | 9528081 | 32938769 | 7622446 | 9759836 |
| 22886028 | 16464983 | 32489508 | 12960148 | 14570902 |
| 15261132 | 15640247 | 32917722 | 11588191 | 10825383 |
| 31726893 | 12437990 | 33083157 | 11120810 | 9873047 |
| 15279543 | 19854871 | 32592996 | 10891480 | 8621725 |
| 30902655 | 18333962 | 33244168 | 11408617 | 9927426 |
| 3703020 | 27288453 | 32511376 | 10835426 | 9679149 |
| 18443746 | 27507650 | 32169673 | 15677321 | 12600999 |
| 28229088 | 22939629 | 33072893 | 12628924 | 11856330 |
| 8632302 | 21931736 | 32653452 | 10744767 | 12393899 |
| 3561384 | 19623215 | 33082293 | 11959661 | 10444401 |
| 15549298 | 22905747 | 32706371 | 10593906 | NA |
| 16141383 | 17500595 | 32404436 | 12242299 | 11916537 |
| 12919933 | 26153216 | 32970989 | 12495932 | 12070163 |
| 3588607 | 24058414 | 32944968 | 3141589 | 10716930 |
| 17077318 | 24164323 | 33801464 | 15488193 | 11701463 |
| 23539623 | 19262167 | 34127972 | 9328344 | 11082445 |
| 28939686 | 15136563 | 33391280 | 11708838 | 15826941 |
| 5323048 | 11238932 | 33372174 | 11080204 | 11454875 |
| 10989432 | 15752761 | 26579391 | 11292831 | 11245462 |
| 30823446 | 10611234 | 33335518 | 12167697 | 12896977 |
| 15093568 | 24920159 | 32529952 | 15064721 | 12855697 |
| 7936305 | 23752268 | 17490702 | 12496361 | 12181435 |
| 25030255 | 28295012 | 33101306 | 12351703 | 12446733 |
| 16541364 | 9885572 | 19430490 | 10783132 | 11859076 |
| 31744820 | 15917271 | 15580300 | 11907026 | 8900202 |
| 17462537 | 20463145 | 34788596 | 17492052 | 7797459 |
| 21508345 | 23116402 | 15681410 | 7731720 | 12738761 |
| 25595279 | 21110228 | 26260141 | 8633244 | 12414794 |

| | | | | |
|---|---|---|---|---|
| 10942386 | 8622647 | 11032810 | 10751227 | 11053438 |
| 10226075 | 10022874 | 12498787 | 11751459 | 10431683 |
| 11673483 | 11839761 | 9607940 | 12690113 | 10196167 |
| 10781029 | 15156153 | 12782630 | 9528766 | 10753939 |
| 9169451 | 11441089 | 12588875 | 10958792 | 12970364 |
| 12595573 | 11058585 | 8887643 | 12226077 | 10706690 |
| 9379049 | 9218480 | 9727040 | 11400324 | 10827000 |
| 11245588 | 11038347 | 406657 | 9065414 | 12218141 |
| 8808177 | 10854065 | 11846562 | 8627578 | 9755064 |
| 9439849 | 11376011 | 7592979 | 12649265 | 9831561 |
| 10488088 | 11956220 | 9430721 | 15153095 | 12917107 |
| 10330161 | 8756648 | 12839832 | 8969228 | 11827958 |
| 9725212 | 14654098 | 11490018 | 15151905 | 12810613 |
| 11306453 | 10788495 | 10325235 | 12529253 | 15805288 |
| 10644746 | 11971024 | 9688849 | 7923353 | 15657416 |
| 11732999 | 9733801 | 8940180 | 9422727 | 12738797 |
| 11278709 | 14499342 | 11395491 | 11560921 | 10693929 |
| 9808624 | 8700529 | 11157082 | 15640156 | 10026206 |
| 8626650 | 11571230 | 9139689 | 11315998 | 9569230 |
| 11283246 | 9852158 | 12208764 | 12842874 | 14681544 |
| 11520792 | 11950876 | 11463794 | 12670941 | 8626761 |
| 15262987 | 12810757 | 8765993 | 10096607 | 9891015 |
| 9016798 | 11278353 | 15688010 | 7499206 | 9885231 |
| 12589052 | 12145292 | 17190838 | 14676298 | 9973202 |
| 10807788 | 14984580 | 9687510 | 12032150 | 11487658 |
| 9361191 | 9675184 | 11500497 | 11303030 | 12621039 |
| 10931853 | 11574474 | 15213298 | 10477597 | 12446726 |
| 11696358 | 12491767 | 7836388 | 15491994 | 10199807 |
| 9006914 | 11416000 | 10918063 | 11238126 | 11124968 |
| 9094716 | 10704466 | 11574420 | 11500363 | 9669024 |
| 8717044 | 8910314 | 7559496 | 14963045 | 14729955 |
| 11812784 | 14557275 | 10666409 | 11274965 | 12808055 |
| 15322221 | 7744823 | 14532295 | 12796499 | 12524221 |
| 10951185 | 12960165 | 10993748 | 8571671 | 10706116 |
| 14668344 | 15650183 | 12225966 | 10747974 | 14988405 |
| 12649327 | 9130707 | 11006268 | 10066767 | 8995385 |
| 7761832 | 14749369 | 11880369 | 11406578 | 16760378 |
| 9207092 | 11352924 | 10642513 | 11503142 | 15967790 |
| 8668348 | 11017917 | 11463795 | 7533300 | 12239105 |
| 7744815 | 11123233 | 11035106 | 12181454 | 11296227 |
| 10891587 | 8621389 | 8687465 | 10438924 | 12181434 |
| 8939929 | 11983682 | 9391009 | 15078882 | 11171588 |
| 9346921 | 15201137 | 12721328 | 12169624 | 14688255 |
| 10713965 | 7896797 | 10463587 | 12746434 | 12724418 |
| 11124982 | 14970203 | 10487749 | 12832465 | 10318869 |
| 12496377 | 10777559 | 10969079 | 8887554 | 12193412 |
| 8798679 | 11879201 | 11328854 | 1313322 | 10024371 |
| 9642252 | 11454682 | 11350959 | 14702343 | 10788492 |
| 15155836 | 11591753 | 12623839 | 14575867 | 8710867 |
| 10636870 | 11090049 | 10976102 | 12801936 | 9724043 |
| 12584202 | 10428835 | 8429046 | 8617731 | 10650934 |

| | | | | |
|---|---|---|---|---|
| 9039124 | 12110168 | 15085196 | 18234890 | 17374610 |
| 9651336 | 25483003 | 17043353 | 7592771 | 10094440 |
| 10098484 | 16144902 | 14765107 | 19289301 | 22418790 |
| 11895866 | 18579680 | 10224053 | 19880519 | 17360901 |
| 9316442 | 18485326 | 20145241 | 15918042 | 16764822 |
| 12562867 | 15823095 | 19706601 | 22000980 | 16006741 |
| 1908778 | 11756451 | 11591434 | 20965158 | 18928334 |
| 8713997 | 16100120 | 15452130 | 9933599 | 18633768 |
| 10391889 | 26853464 | 16399501 | 16916793 | 16938762 |
| 7608156 | 17977534 | 10889017 | 10421804 | 18032528 |
| 11487632 | 33414460 | 20817729 | 20124538 | 15758173 |
| 17464208 | 30108193 | 22031849 | 10978336 | 18639564 |
| 8393279 | 34064003 | 20541520 | 21836131 | 18384375 |
| 15220929 | 12011103 | 23300015 | 16916659 | 19729586 |
| 11752209 | 34102611 | 10318878 | 20880945 | 22226696 |
| 7713937 | 32839400 | 22154484 | 19120039 | 16275016 |
| 9057106 | 29090099 | 10397761 | 19958302 | 19278673 |
| 12060777 | 4064003 | 22684109 | 20847275 | 7908738 |
| 8382677 | 30885289 | 21240525 | 21555452 | 17142880 |
| 1683468 | 30060809 | 15266058 | 20209124 | 17553693 |
| 10542136 | 27812877 | 16831124 | 10197817 | 18792810 |
| 15546818 | 19416851 | 21696544 | 12065627 | 14608012 |
| 11259513 | 19029065 | 22984430 | 20198379 | 9038229 |
| 14724213 | 18216770 | 20670887 | 20351659 | 9676432 |
| 16903857 | 17928229 | 15899885 | 15584216 | 8957963 |
| 9395450 | 17621255 | 11358865 | 17077377 | 9259266 |
| 7985086 | 22865681 | 9488713 | 17973628 | 9602135 |
| 14715266 | 17065982 | 19126544 | 18778441 | 2058745 |
| 11606615 | 17646659 | 18835813 | 14756806 | 18599790 |
| 11818496 | 20855588 | 18286597 | 17556193 | 18192214 |
| 12554779 | 17347648 | 14693395 | 17584746 | 11335745 |
| 16303171 | 18046414 | 17568567 | 18085326 | 20226894 |
| 10781004 | 17581637 | 18084013 | 17354259 | 10575032 |
| 11181532 | 20595232 | 17597646 | 8841406 | 21204411 |
| 16189297 | 20732877 | 15056938 | 16830328 | 19247214 |
| 10217295 | 17290267 | 22351881 | 19559077 | 9645961 |
| 17452640 | 21473868 | 14741405 | 14575242 | 18180303 |
| 9620655 | 19015276 | 21865888 | 17613521 | 16728402 |
| 28924389 | 10770267 | 19772970 | 11080175 | 18045290 |
| 30487431 | 9106493 | 15240857 | 22371606 | 18675942 |
| 27378692 | 8402897 | 18041756 | 16144830 | 19396608 |
| 12110169 | 20103737 | 20080142 | 21600882 | 7745608 |
| 15902259 | 21459323 | 18094064 | 16467535 | 19476549 |
| 17360776 | 19759400 | 17869437 | 12393272 | 18771483 |
| 29761784 | 10500120 | 19369586 | 19841066 | 22179534 |
| 16505380 | 21816276 | 17101276 | 20009079 | 21976671 |
| 16728472 | 21664250 | 15273283 | 18810510 | 15199055 |
| 16342160 | 22436747 | 17311278 | 18571009 | 18534229 |
| 19608404 | 15680331 | 20937660 | 17709376 | 16322073 |
| 18849530 | 10805775 | 19801649 | 12599191 | 11259344 |
| 12529645 | 17296605 | 19666841 | 22334710 | 12439736 |

| | | | | |
|---|---|---|---|---|
| 21383205 | 31611699 | 26482876 | 30065554 | 11171046 |
| 22449172 | 17719247 | 19196800 | 21076395 | 28007913 |
| 29934975 | 8146197 | 28235572 | 20951942 | 12427739 |
| 16480962 | 7543139 | 28723567 | 28109176 | 9182576 |
| 17765940 | 35727133 | 27449104 | 24122795 | 23076213 |
| 17186017 | 16208375 | 21683758 | 23435428 | 24689035 |
| 15135306 | 17182562 | 17574003 | 18433497 | 28289053 |
| 19250197 | 1423621 | 31076567 | 19940144 | 12154000 |
| 8242225 | 26473606 | 18259611 | 28820180 | 25910937 |
| 16516475 | 11520798 | 27681418 | 17055431 | 21636859 |
| 17680028 | 23530057 | 26207953 | 21670757 | 16006559 |
| 17041621 | 11704864 | 16870170 | 15044469 | 11682481 |
| 19143648 | 11402335 | 16428285 | 25848864 | 19050761 |
| 16469695 | 29237129 | 30886364 | 33506952 | 12716911 |
| 25310982 | 28712664 | 221813 | 32726355 | 9450544 |
| 6998725 | 24324645 | 21455117 | 32695122 | 25956888 |
| 29608137 | 11057902 | 17974478 | 33596266 | 22124154 |
| 23791173 | 24069422 | 22649723 | 33337934 | 16893970 |
| 23334421 | 22383895 | 14751757 | 24493213 | 19295129 |
| 24449267 | 10964259 | 23634849 | 23483280 | 23830865 |
| 23685455 | 17452290 | 21047732 | 29180617 | 28535874 |
| 24799285 | 17666482 | 11830511 | 29107536 | 21192934 |
| 9489702 | 24872083 | 9774969 | 28658158 | 11387210 |
| 25435019 | 15964803 | 11423969 | 20301631 | 20844008 |
| 22066971 | 20167242 | 10391251 | 20301338 | 20826718 |
| 19375514 | 23045339 | 17139329 | 27132995 | 16966330 |
| 11861617 | 27351946 | 29892481 | 30337552 | 11744618 |
| 9784967 | 21511232 | 7799943 | 28293384 | 12588761 |
| 15246961 | 22653837 | 17167474 | 20301360 | 11161716 |
| 32726801 | 28322461 | 21807935 | 30158522 | 10938010 |
| 29517999 | 11149600 | 22969750 | 16940241 | 21148427 |
| 23405894 | 18329679 | 20530740 | 30588060 | 22592917 |
| 31996678 | 24463099 | 14739072 | 2722838 | 7657594 |
| 29568499 | 27693079 | 21193034 | 11465067 | 18206965 |
| 12172553 | 28448947 | 22079269 | 11532180 | 10067868 |
| 18991293 | 20802499 | 30456354 | 1988037 | 15928081 |
| 19143635 | 8222576 | 38260464 | 14597963 | 14525795 |
| 1847722 | 25330987 | 27641100 | 2112956 | 26221892 |
| 9429890 | 18467435 | 19932708 | 17687269 | 15247219 |
| 11395409 | 15971998 | 21189261 | 16945988 | 15180964 |
| 15778376 | 29997323 | 17161604 | 9353388 | 9858476 |
| 19710469 | 21317437 | 4400078 | 18838507 | 9169492 |
| 8676075 | 20378719 | 19826765 | 8168173 | 17686471 |
| 15546391 | 25390014 | 25860609 | 1676665 | 16627366 |
| 9973453 | 21448409 | 29902437 | 15761113 | 30086460 |
| 29915297 | 24820868 | 31825083 | 21843514 | 16336951 |
| 23922331 | 17574010 | 31258331 | 26655500 | 18450586 |
| 18515054 | 11533250 | 23797870 | 15448146 | 11494124 |
| 11406366 | 20051527 | 29468141 | 26136364 | 23718729 |
| 17015696 | 22353746 | 30619249 | 23000962 | 10688886 |
| 28851713 | 17720805 | 22688187 | 19181962 | 20507983 |

| | | | | |
|---|---|---|---|---|
| 20332118 | 16974068 | 20813203 | 17276402 | 23592840 |
| 25644401 | 24785348 | 10331420 | 9405464 | 18723443 |
| 21212275 | 17178724 | 16484616 | 15548136 | 22145046 |
| 24662486 | 15207703 | 15184502 | 25387128 | 17658244 |
| 11108718 | 19494114 | 16679383 | 19923922 | 15692085 |
| 23029280 | 9393975 | 24746698 | 17993608 | 19715393 |
| 25331892 | 29222111 | 19228841 | 12490545 | 2656356 |
| 19435802 | 14514674 | 15920022 | 28183800 | 9453379 |
| 11090059 | 28418925 | 19718476 | 10022890 | 8878137 |
| 11335727 | 9461619 | 11228166 | 11678628 | 8948389 |
| 11704645 | 16403913 | 21586748 | 9804796 | 1444060 |
| 24931163 | 26063728 | 22689825 | 12244099 | 12806616 |
| 18952368 | 21169383 | 21245381 | 22264731 | 1595584 |
| 9299537 | 15545625 | 20432469 | 15854902 | 1416048 |
| 7896817 | 22101521 | 23851566 | 21873429 | 18289917 |
| 11696015 | 11278553 | 28946938 | 20422004 | 12401878 |
| 20400538 | 12844492 | 17065532 | 24966171 | 12937841 |
| 16479592 | 15026417 | 29051140 | 16787925 | 10227052 |
| 19965691 | 21653826 | 9722576 | 20434959 | 11585356 |
| 16909199 | 22854047 | 12086892 | 19914243 | 8017322 |
| 18787075 | 27052191 | 26846344 | 11927607 | 3372162 |
| 10544009 | 12654612 | 16760434 | 17060906 | 3019190 |
| 15051508 | 10347193 | 12214271 | 14525763 | 9627692 |
| 15016650 | 11719508 | 10048588 | 29401587 | 15358783 |
| 25157100 | 11278468 | 11238453 | 27695625 | 15911617 |
| 17004325 | 10961983 | 15371454 | 23341459 | 26364851 |
| 20129920 | 19038867 | 21524749 | 15637071 | 19700356 |
| 26217013 | 23732519 | 19883397 | 8769777 | 21114891 |
| 12529448 | 19325137 | 12717443 | 19834490 | 17892308 |
| 22763125 | 22287577 | 23401740 | 16373578 | 11215515 |
| 24300896 | 9580552 | 29545238 | 14656735 | 12860264 |
| 10327068 | 12480817 | 18509061 | 18936167 | 4390543 |
| 15308628 | 10196157 | 15918795 | 12771128 | 20110595 |
| 20463056 | 15261145 | 18059339 | 15131009 | 17045981 |
| 18840614 | 15659776 | 20631299 | 19661463 | 19707742 |
| 14564009 | 28927665 | 17519230 | 15919658 | 6885824 |
| 11513746 | 9393862 | 29880492 | 28271280 | 1492101 |
| 9398617 | 15217908 | 17371830 | 18617643 | 18226574 |
| 20110358 | 18440775 | 10102632 | 10865940 | 3954771 |
| 15466206 | 18451337 | 23401860 | 12714333 | 11279066 |
| 28278510 | 12855698 | 18559514 | 20445537 | 14695536 |
| 28536097 | 22762016 | 23805312 | 11280761 | 16690356 |
| 28851877 | 17303569 | 11525641 | 19661918 | 12571671 |
| 12407018 | 18332134 | 22773844 | 19995915 | 18563633 |
| 18981713 | 22438576 | 18440854 | 19478092 | 11832420 |
| 18180305 | 21711246 | 20413783 | 20061392 | 6107306 |
| 11015619 | 29065929 | 28478454 | 15001553 | 15117956 |
| 17721515 | 10336480 | 28424170 | 18805968 | 17163662 |
| 17908694 | 20966350 | 25582201 | 21905169 | 15631778 |
| 15342917 | 16456544 | 21653897 | 9756915 | 19940234 |
| 11122379 | 17372230 | 18377662 | 29100366 | 16472406 |

| | |
|---|---|
| 3134526 | 17372190 |
| 12419480 | 24416395 |
| 6668142 | 23506894 |
| 15782407 | 17344318 |
| 16239338 | 11831458 |
| 12663279 | 23506888 |
| 9159205 | 22289350 |
| 11472746 | 24710731 |
| 19373259 | 22057392 |
| 1597321 | |
| 10468579 | |
| 12816923 | |
| 10377254 | |
| 17964036 | |
| 11012887 | |
| 31825581 | |
| 16763894 | |
| 28202503 | |
| 31867158 | |
| 25071589 | |
| 15128933 | |
| 20541252 | |
| 20736230 | |
| 18689389 | |
| 20418485 | |
| 7690968 | |
| 11111101 | |
| 22782502 | |
| 23923049 | |
| 21870057 | |
| 15205382 | |
| 24247221 | |
| 16399349 | |
| 24202443 | |
| 16186133 | |
| 17721433 | |
| 24726990 | |
| 16002434 | |
| 24975273 | |
| 22213316 | |
| 24854954 | |
| 19667142 | |
| 23999061 | |
| 21872797 | |
| 25163748 | |
| 21459326 | |
| 18800146 | |
| 20091054 | |
| 18483421 | |
| 16881513 | |
| 18957414 | |
