## Supplementary File S2 for "CENTRA: Knowledge-Based Gene Contexuality Graphs Reveal Functional Master Regulators by Centrality and Fractality"

0432ccr
10depart
10e
130b
13s
17depart
18a
1cancer
1cell
1depart
1department
1divis
1institut
1laboratori
1urolog
2443200200591x
27a
2cell
2depart
302b
33b
3depart
3T3
4depart
5472can
5depart
6depart
7depart
7e3
8290cd
8depart
9depart
aachen
aarhus
abil
abolish
abrog
absenc
abund
acad
academ
academi
acceptor
accord
accumul
achiev
acid
across
act
acta
action
activ
adapt
addit
address
adelaid
administr
adolfogarcia
adult
advanc
advisori
affect
affili
affin
age
agent
agt
aid
aim
air
albert
alberta
allow
alon
along
alpha
also
alter
altern
although
america
american
amgen
amino
among
amount
amsterdam
analys
analysi
analyz
and
anderson
angel
anim
anoth
anti
antwerp
appar
appear
appli
applic
approach
approv
approxim
apr
arabidopsi
arbor
arch
are
area
argentina
arkadia
arm
around
articl
aspect
assay
assembl
assess
assign
associ
astrazeneca
athen
atlas
attach
attenu
auckland
aug
augusta
australia
austria
author
avail
avenu
babraham
background
baltimor
balu
bangalor
barbara
Barbara
barcelona
base
basi
basic
bath
baylin
bec
becom
become
been
behalf
behavior

beij
belgium
benefit
berg
berkeley
berlin
bern
beroukhim
beta
bethesda
better
between
beuren
bind
bingen
bioactiv
biochem
biochemistri
bioengineering
bioinformat
biol
biolog
biomed
biomedical
biophi
biophys
bioscienc
biostatist
biotechnolog
block
blot
board
bochum
bodi
body
bohr
bologna
bond
bonn
bosco
boston
both
bound
bovin
bowen
box
branch
braunschweig
brazil
break
bristol
broad
bronx
brook
brown
brussel
bueno
build
but
butyr
california
call
cambridg
campbel
campus
can
canada
cancer
candid
capabl
capac
cardiff
carolina
carri
carrier
carter
cascad
case
cat
catalog
catalonia
caus
cdna
cedex
cell
cellular
center
centr
central
certain
cerveau
chain
chang
changsha
chapel
character
characterist
charitÃ©
chem
chemic
chemistri
chen
chicago
Chien
child
children
chile
chimer
china
chines
chip
cincinnati
citi
clark
class
classic
classif
clear
cleavage
cleveland
clin
clinic
clinicopatholog
clock
clone
close
cnrs
Cobra1
code
cohort
coimbra
collabor
collect
colleg
cologn
coloni
colorado
columbia
combin
commens
comment
common
communiti
compar
comparison
compet
complet
complex
compon
composit
comprehens
compris
comput

computing
concentr
concern
conclude
conclus
condit
conduct
confer
confirm
conflict
congenit
connect
consequ
conserv
consid
consist
consortium
constitut
construct
consult
contain
content
context
contract
contrast
contribut
control
convers
convert
cooper
coordin
copenhagen
copyright
cord
cordeli
core
corporate
correl
correspond
council
coupl
coval
cpt
creighton
critic
cross
crucial
cruzi
crystal
CSIR
csir
cultur
curat
current
custom
daegu
dalla
dana
danver
data
database
dataset
davi
day
dec
decad
declar
decreas
dedic
defect
defici
defin
definit
degrad
delaware
deliver
della
delta
demonstr
denmark
densiti
dentistri
depart
depend
deplet
deriv
des
descart
describ
descript
design
despite
detail
detect
determin
dev
develop
DFG
diagnost
diderot
die
diego
differ
ding
dion
direct
director
discov
discoveri
discuss
disord
display
dispo
disposit
disrupt
distal
distinct
distribut
divers
divis
doi
doi:
domain
dongguan
donor
dose
download
downstream
dresden
drive
driven
dublin
due
duke
dunde
earli
east
ecollect
edinburgh
educ
effect
effici
einstein
either
electron
element
elena
elev
elimin
elsevi
elucid
emboj
emerg

emeritus
employ
enabl
encod
end
endocrinolog
endocrinologist
endogen
eng
engag
engel
engin
engl
enhanc
enrich
enter
entri
environment
epub
equal
erasmus
essen
essenti
establish
ester
ethics
euclid
european
evalu
even
event
evid
evolut
ewe
examin
exampl
exchang
exert
exhibit
exist
expans
experi
experiment
explain
explor
expos
express
extens
extent
extract
faïenceri
facilit
factor
faculti
fail
fairlamb
famili
farber
fast
fate
featur
feb
feder
feed
feinberg
ferrara
filter
final
financi
find
finland
first
fish
five
florenc
florida
fluid
fluoresc
focus
fold
follow
for
forc
form
format
found
foundat
four
fragment
franc
francisco
fred
frederick
free
freiburg
frequent
from
front
fts
fuch
fudan
full
fulli
fulton
fumar
function
fungoid
furthermor
furthermore
fxiii
gÃ©nÃ©tiqu
gÃ¶ttingen
gabriel
gain
galveston
gamma
Gaslini
gene
genentech
general
generat
genet
genom
georgia
german
germani
getz
ghent
gibb
gilman
giovanni
given
gku1267
glasgow
global
golub
gothenburg
graduat
grafton
grant
graz
great
greater
greatwal
greec
green
gregorio
greifswald
grenobl
griffith
groningen
gross
group

growth
guangdong
guangzhou
guo
gupta
gustav
hÃ´pital
hairi
hall
hamon
hampton
hangzhou
hannov
harbin
harbor
harvard
has
hasegawa
have
haven
hay
head
health
healthi
heidelberg
help
helsinki
hepatolog
heuvel
high
higher
highlight
hill
hippel
histori
hoadley
hong
hopkin
hornbeck
hospit
hospital
houston
howev
ht2a
huang
hubei
hugh
hum
human
hunan
hundr
hybrid
hypothesi
ibaraki
icahn
ident
identif
identifi
identified
iii
illinoi
illkirch
immunopatholog
impact
impair
implic
import
impos
improv
inc
includ
incorpor
increas
incub
independ
index
indexed
india
indic
individu
induc
induct
influence
inform
information:
inh
inhibit
initi
inlb
inner
innov
insight
institut
institutet
intak
integr
interact
interest
interf
intermedi
intern
intersect
intervene
intervention
intracellular
investig
involv
iowa
ircc
isol
israel
issu
itali
its
J1538
jackson
jacksonvill
jacob
jan
jan43databas
japan
jbcm801400200
jccr200911025
jccr201003022
jena
jiangsu
jinan
jmolcel201104017
jmolcel201108039
john
johnson
jolla
jone
jonsson
joseph
journal
jstem201006012
jul
jun
kakusan
kanazawa
karolinska
kda
kentucki
ketter
kevanshokatucsfedu
key
kiel
kim
kingdom
kink
knockdown
knowledg

knowledgebas
known
kobe
kong
korea
kornhaus
kroemerorangefr
kyoto
kyowa
kyungpook
lab
label
laboratori
lack
ladanyi
laforin
lai
laird
lancet
lander
lane
langon
lara
larg
larizza
larsson
last
late
later
latham
lau
lausann
lawrenc
layer
lead
least
leav
led
lee
leicester
leiden
len
length
lesion
less
lett
leuven
level
lewi
life
light
like
limit
lin
lindau
line
linehan
link
linker
literatur
littl
liu
live
local
locat
london
long
loop
lopez
los
loss
loui
low
lower
ltd
ltedaro
lund
luxembourg
lyon
mÃ¡laga
m1234
m617
maastricht
madan
made
madison
madrid
maharashtra
main
maintain
mainten
major
make
male
mammalian
manag
mani
manitoba
manner
map
mar
marburg
mari
mark
marker
marra
marseill
martinsri
maryland
mass
massachusett
massagu
match
matter
matur
may
mayo
mcb00624
mcgill
mckay
measur
mechan
mechanist
med
mediat
medic
medicin
medium
medlin
medline]
meibergdreef
meier
mel
melbourn
member
merck
method
meyerson
miami
mice
michael
michigan
microm
midkin
might
mill
miller
mim
min
ministri
mishima
mix
mmol

model
modif
modifi
modul
moell
mol
molecul
molecular
monash
monotherapi
month
montr
moor
moreov
morgan
morri
morton
motif
mount
mous
much
multipl
munich
municip
murin
murray
mutat
myer
nagoya
nakao
name
nanchang
nanj
nanomateri
nanoparticl
nant
nar
nashvill
nat
nation
natl
natur
near
nearest
necessari
neck
need
negat
neighbor
nephrolog
net
netherland
network
neurologi
nevankroganucsfedu
new
newli
nice
nih
nijmegen
nine
node
nogo
non
normal
north
norway
not
nottingham
nov
novarti
novel
now
npc
null
number
observ
obtain
occur
oct
often
ohio
oic
one
ontario
open
order
organ
ortho
osaka
oslo
ospedaliera
other
ottawa
oulu
our
outcom
overal
overexpress
oxford
PA1
pac
padova
padua
pain
pair
palczewski
pandey
paramet
pari
park
parker
parkvill
part
partial
particl
particular
partner
parvin
pasteur
patel
patholog
pathway
patient
pattern
paulo
PCR
pdt
peke
penni
pennsylvania
peopl
per
perform
period
perou
peter
pharmaceut
pharmacolog
phenotyp
phi
philadelphia
phornbeckcellsignalco
m
phosphositeplusÂ®
phosphositeplus
physic
physiolog
pick
pig
pit
pitiÃ©
pittsburgh

pkciota
plant
platania
play
plus
pmc4383998
pmcid
pmcid:
pmid
pmid:
point
pokfulam
poland
polymer
poor
por
portland
porto
portug
posit
possibl
post
potent
potenti
power
pre
precis
precursor
predomi
predomin
prefer
prepar
presenc
present
press
pretreat
prevent
previous
primari
princeton
princip
pro
probabl
proc
process
produc
product
professor
profil
program
progress
prolifer
promot
properti
propos
protect
protein
proteom
provid
provinc
proxim
psi
pubert
public
publish
pulpa
purifi
putat
qbi
qcrg
qingdao
qualiti
queen
rabbit
radboud
rang
rant
raph
rapid
rare
rat
rate
rather
rathmel
ratio
reaction
real
reanalyz
recalibr
receiv
recent
recherch
recognit
recombin
recruit
reduc
reduct
redund
reed
reflect
regard
regina
region
regul
regulatori
rehovot
relat
relationship
releas
relev
reliabl
remain
remark
remov
repeat
report
republ
republi
requir
res
research
reserv
residu
resolut
resourc
respect
respond
respons
restrict
result
results
retriev
rev
reveal
revers
review
rewir
rich
rickett
right
road
robertson
robinson
roch
rockvill
role
rome
root
roussi
rout
royal
rzburg
sÃ£o

s00281
s00424
s0140
S0955
s1097
s41586
saint
saitama
saksena
salamanca
salem
salpÃªtriÃ¨r
salt
sampl
san
sander
sansom
sant
santa
santiago
sastremssmedu
sato
scale
scatter
schmidt
school
schultz
schwarz
sci
scienc
scientifiqu
scotland
scripp
seattl
second
secret
section
select
self
sendai
sens
sensor
seoul
sep
sequenc
serono
serv
set
seven
sever
shandong
shanghai
share
shelton
shen
shenzhen
shi
shift
short
show
shown
sick
side
siegel
siena
signal
signific
similar
simon
sinai
sinc
singapor
singl
singleton
site
six
size
sjonc1209954
skrzypek
sleep
sloan
small
smith
societi
sofferenza
softwar
solut
song
sougnez
sourc
south
southern
southwestern
space
spain
spanish
speci
special
specif
specimen
spectra
spectrometri
spellman
spread
spring
squibb
sri
stabil
stage
standard
standford
stanford
state
statement
statist
status
step
still
stimul
stockholm
strain
strategi
street
stress
strike
strong
structur
studi
subject
subsequ
subset
substrat
subtyp
subunit
success
succursal
suffici
suggest
suita
summar
sun
supplement
support
suppress
surfac
surgery
surprising
surviv
suscept
sustain
sw3
swansea
sweden

switch
switzerland
sydney
symptom
system
taichung
taipei
taiwan
takahashi
taken
talk
tam
tamper
tanaka
tang
tanpakushitsu
target
tati
tcga
tcpobop
technisch
technolog
term
termin
terminus
test
texa
thaliana
that
the
their
therapeut
therebi
therefor
these
this
thoma
thompson
thought
three
through
thus
tianjin
tight
time
tip
tiqu
tissu
tochigi
togeth
tokio
tokyo
tongji
tool
tooth
toronto
total
toward
tran
transcript
transduct
transfect
transfer
transit
translat
transloc
transmit
transport
treat
treatment
trend
triangl
triest
trigger
tromsø
tsinghua
tsmedizin
tsukuba
ttingen
tube
tucson
tuebingen
tulan
turkey
turku
turn
tweak
two
type
umr
unclear
under
undergo
understand
understood
unexpect
union
uniprotkb
uniqu
unitÃ©
unit
univers
universidad
université
universit
universitario
unknown
unrel
upon
uppsala
uptake
usa
usa.
use
usual
util
utrecht
valid
valu
van
vanderbilt
variabl
variant
variat
varieti
various
vault
verhaak
versus
via
victoria
vienna
vill
vitro
vivo
wale
wang
was
washington
water
week
weinstein
weisenberg
weizmann
well
wellcom
were
western
wheeler
wherea
whether
which
whose

wide
wild
wiley
wilkinson
will
willem
william
winnipeg
wisconsin
with
within
without
women
wong
wood
work
worldwid
wuhan
wwwphosphositeorg
xenopus
xiangya
xiii
yale
yang
yangzhou
yao
year
yenepoya
yet
yield
york
young
zhang
zhejiang
zhou
zhu
zone
zurich
